# Controlling Selectivity of Modular Microbial Biosynthesis of Butyryl-CoA-Derived Designer Esters

**DOI:** 10.1101/2021.10.19.465013

**Authors:** Jong-Won Lee, Cong T. Trinh

## Abstract

Short-chain esters have broad utility as flavors, fragrances, solvents, and biofuels. Controlling selectivity of ester microbial biosynthesis has been an outstanding metabolic engineering problem. Here, we present a generalizable framework to enable the *de novo* fermentative microbial biosynthesis of butyryl-CoA-derived designer esters (e.g., butyl acetate, ethyl butyrate, butyl butyrate) with controllable selectivity. Using the modular design principles, we generated the butyryl-CoA-derived ester pathways as exchangeable production modules compatible with an engineered chassis cell for anaerobic production of designer esters. We designed these modules derived from an acyl-CoA submodule (e.g., acetyl-CoA, butyryl-CoA), an alcohol submodule (e.g., ethanol, butanol), a cofactor regeneration submodule (e.g., NADH), and an alcohol acetyltransferase (AAT) submodule (e.g., ATF1, SAAT) for rapid module construction and optimization by manipulating replication (e.g., plasmid copy number), transcription (e.g., promoters), translation (e.g., codon optimization), pathway enzymes, and pathway induction conditions. To further enhance production of designer esters with high selectivity, we systematically screened various strategies of protein solubilization using protein fusion tags and chaperones to improve the soluble expression of multiple pathway enzymes. Finally, our engineered ester-producing strains could achieve 19-fold increase in butyl acetate production (0.64 g/L, 96% selectivity), 6-fold increase in ethyl butyrate production (0.41 g/L, 86% selectivity), and 13-fold increase in butyl butyrate production (0.45 g/L, 54% selectivity) as compared to the initial strains. Overall, this study presented a generalizable framework to engineer modular microbial platforms for anaerobic production of butyryl-CoA-derived designer esters from renewable feedstocks.

## Introduction

Esters are industrial platform chemicals with versatile applications as flavors, fragrances, solvents, and biofuels (Lee and Trinh, 2020). Microbial biosynthesis of esters from lignocellulosic biomass can potentially offer an alternative promising solution to the current petroleum-based process that is neither renewable nor sustainable (Chubukov et al., 2016; Seo et al., 2019). For bioenergy applications, short-chain (C6-C10) esters have recently been attracting attention as drop-in fuels due to its favorable properties such as high energy density (Layton and Trinh, 2016b), high hydrophobicity (Tai et al., 2015), and good compatibility with current infrastructures including engines, transport, and storage density (Contino et al., 2013a; Jenkins et al., 2013). For instance, ethyl valerate (C7) (Contino et al., 2013b), butyl butyrate (C8) (Chuck and Donnelly, 2014; Jenkins et al., 2013), butyl valerate (C9) (Contino et al., 2013a), and pentyl valerate (C10) (Contino et al., 2013a) are good fuel additives for gasolines while butyl butyrate (C8) (Chuck and Donnelly, 2014; Jenkins et al., 2013) and ethyl octanoate (C10) (Chuck and Donnelly, 2014) are considered as an alternative jet fuel.

In nature, eukaryotic cells utilize alcohol acetyltransferases (AATs) to condense an alcohol and acetyl-CoA to make acetate esters in a thermodynamically favorable reaction, as often found in plants and fruits for generating scents (D’Auria, 2006) or in fermenting yeasts for making flavors (van Wyk et al., 2018). Inspired by nature, microbial biomanufacturing platforms (e.g., *Escherichia coli)* have been engineered to make these acetate esters directly from fermentable sugars (Chacon et al., 2019; Horton and Bennett, 2006; Horton et al., 2003; Layton and Trinh, 2014; Layton and Trinh, 2016a; Lee and Trinh, 2019; Rodriguez et al., 2014; Vadali et al., 2004). Remarkably, the substrate promiscuity of AATs also enables microbial biosynthesis of acylate esters beyond acetate esters including propionate esters (Layton and Trinh, 2016a), lactate esters (Lee and Trinh, 2019; Seo et al., 2021), butyrate esters (Layton and Trinh, 2014), pentanoate esters (Layton and Trinh, 2016a), and hexanoate esters (Layton and Trinh, 2016a). Therefore, harnessing diversity of AATs, acyl-CoAs, and alcohols can result in the *de novo* microbial biosynthesis of a vast library of esters from renewable feedstocks for useful applications.

To enable a systematic and rapid generation of microbial biocatalysts to produce various esters, a modular cell engineering framework has recently been developed (Garcia and Trinh, 2019a; Garcia and Trinh, 2019b; Garcia and Trinh, 2020; Trinh et al., 2015; Wilbanks et al., 2018). Each ester production strain can be assembled from an engineered modular chassis cell and exchangeable ester producing pathways known as production modules. Nevertheless, experimental implementation has been challenging due to the intrinsic complexity of the production modules requiring expression of multiple heterologous enzymes derived from bacteria, yeasts, and plants (Layton and Trinh, 2014; Layton and Trinh, 2016a; Layton and Trinh, 2016b; Lee and Trinh, 2019).

Critical to the effective microbial biosynthesis of a target designer ester is the availability of efficient and robust AATs and precursor metabolite pathways that are compatible with a microbial host (Seo et al., 2021). Selective microbial biosynthesis of acylate esters other than acetate esters is very challenging due to low availability of target acyl-CoAs and alcohols, a high intracellular pool of competing substrates (i.e., non-target acetyl-CoA and alcohols), and inefficient AATs. For instance, the microbial ester production is much less efficient for a butyryl-CoA-derived acylate ester (e.g., butyl butyrate (Feng et al., 2021; Layton and Trinh, 2014)) than for an acetate ester (e.g., isobutyl acetate (Tai et al., 2015; Tashiro et al., 2015), isoamyl acetate (Tai et al., 2015)), due to low product selectivity. In particular, equipped with a butyrate ester pathway, an engineered *E. coli* can generate two acyl-CoAs (i.e., acetyl-CoA, butyryl-CoA) and two alcohols (i.e., ethanol, butanol) from fermentable sugars that can be condensed to form two possible acetate esters (i.e., ethyl acetate (EA), butyl acetate (BA)) and two possible butyrate esters (i.e., ethyl butyrate (EB), butyl butyrate (BB)). Furthermore, effective microbial production of acylate esters has been hampered by the required expression of multiple heterologous enzymes that are not compatible with the host. Specifically, low solubility of eukaryotic AATs in a microbial host is a commonly observed problem (Tai et al., 2015; Zhu et al., 2015). Currently, innovative strategies to produce designer esters with high selectivity and efficiency in a microbial host are very limited.

In this study, we presented systematic design and engineering approaches to tackle the current challenges of microbial biosynthesis of designer esters. As a proof-of-study, we demonstrated the microbial biosynthesis of designer butyryl-CoA-derived esters (i.e., BA, EB, BB) with high selectivity in an engineered modular *E. coli* cell. Specifically, we first developed a combinatorial modular design of the butyryl-CoA-derived ester biosynthesis pathways for rapid construction and optimization. Next, we optimized the culture conditions for expression of multiple pathway enzymes including culture temperatures and inducer concentrations to enhance and balance metabolic fluxes toward the synthesis of the target esters. To further improve the compatibility of the engineered pathways with the modular cell, we screened combinatorial strategies of protein solubilization including codon optimization, the use of fusion tags, and/or coexpression of chaperones to improve the soluble expression of multiple pathway enzymes (e.g., AATs). Finally, we characterized the engineered ester-producing strains under anaerobic conditions with pH-adjustment to achieve the enhanced production of designer esters with high selectivity. Overall, this study presents a generalizable framework for engineering modular microbial platforms for anaerobic production of butyryl-CoA-derived designer esters from renewable biomass feedstocks.

## Results

### Designing a general framework to build exchangeable ester production modules for the *de novo* microbial biosynthesis of designer butyryl-CoA-derived esters

To generate the *de novo* microbial biosynthesis of butyryl-CoA-derived esters from fermentable sugars in *E. coli* (Fig 1a), three major pathways are required including i) acyl-CoA synthesis pathway (butyryl-CoA, acetyl-CoA), ii) alcohol synthesis pathway (ethanol, butanol), and iii) ester synthesis pathway (AAT). Biosynthesis of acetyl-CoA is endogenous and is essential for cell functions. We modularized these fermentative ester pathways into four submodules for rapid pathway construction and optimization including i) submodule 1 (SM1) carrying *E. coli atoB* (*atoB*_Ec_), *Clostridium acetobutylicum Hbd* (*hbd*ea), *C. acetobutylicum crt* (*crt*_Ca_), and *Treponema denticola ter* (ter_Td_) for butyryl-CoA synthesis, ii) submodule 2 (SM2) consisted of *Zymomonas mobilis pdc* (*pdc_Zm_*) and *adhB* (*adhB*_Zm_) or *C. acetobutylicum adhE2* (*adhE2*_Ca_) for alcohol synthesis, iii) submodule 3 (SM3) carrying *Candida boidinii fdh* (*fdh*_Cb_) for NADH regeneration, and iv) submodule 4 (SM4) carrying *Saccharomyces cerevisiae ATF1* (*ATF1*_Sc_, specific for acetate ester synthesis) or *Fragaria* x *ananassa* (cultivated strawberry) *SAAT* (*SAAT*_Fa_, specific for acylate ester synthesis) for ester synthesis (Fig. 1b). In the design, the parts were chosen based on our previous function validation of the butyryl-CoA and alcohol (ethanol, butanol) pathways (Layton and Trinh, 2014), the substrate specificity of AATs against acetate and acylate esters (Layton and Trinh, 2016a), and the expression of a NAD^+^-dependent formate dehydrogenase (Fdh) for enhanced intracellular NADH availability (Lim et al., 2013; Shen et al., 2011). Each exchangeable ester module can be assembled from two parts: i) plasmid “pCore” carrying SM1, a common pathway for biosynthesis of butyryl-CoA-derived esters and ii) plasmid “pDerivatization” carrying SM2-SM3-SM4, variable pathways in butyryl-CoA-derived esters using plasmids with various copy numbers (Fig. 1c, Table S1). Figs. 2a, 3a, and 4a show how each submodule can be assembled to build the biosynthesis pathways of BA, EB, and BB, respectively with a list of enzymes presented in Table S2. By transforming a combination of pCore and pDerivatization into the modular *E. coli* strain, TCS083 *ΔfadE* (DE3) (Layton and Trinh, 2014), we generated a set of initial strains, EcJWBA1-6, EcJWEB1-6, and EcJWBB1-6 for optimizing the designer biosynthesis of BA (Fig. 2b), EB (Fig. 3b), and BB (Fig. 3b), respectively (Table 1).

**Figure 1.**
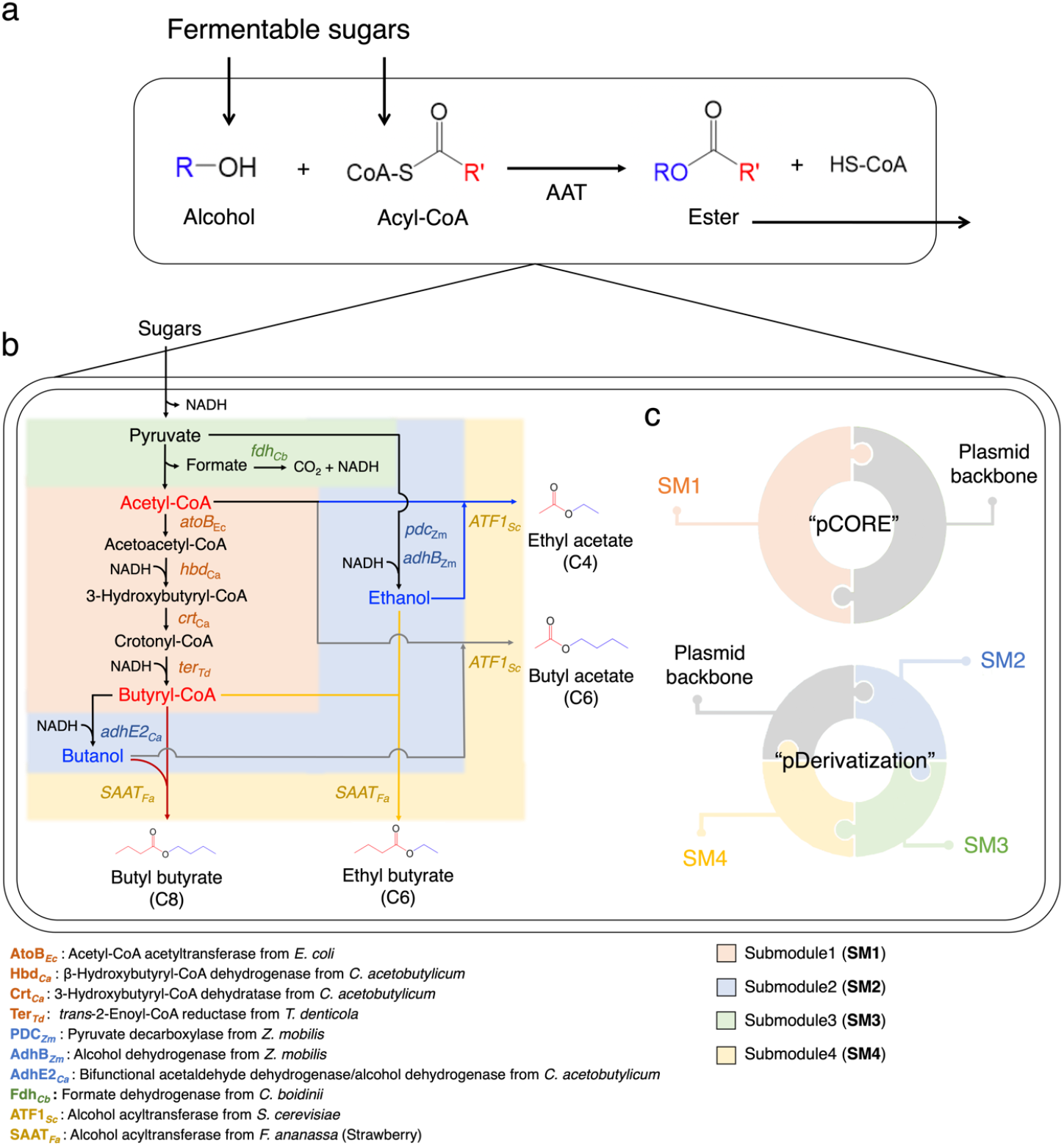
Design of the *de novo* modular microbial biosynthesis of butyryl-CoA-derived esters. **(a)** Biosynthesis of esters by an alcohol acyltransferase (AAT). **(b)** Modular biosynthetic pathways of butyryl-CoA-derived esters. Distinct biosynthesis pathways of each butyryl-CoA-derived ester are presented with colored lines as follows: Ethyl acetate (in blue), Butyl acetate (in grey), Ethyl butyrate (in yellow), and Butyl butyrate (in red). **(c)** Schematic representation of modular plasmid assembly to build the butyryl-CoA-derived ester pathways.

**Figure 2.**
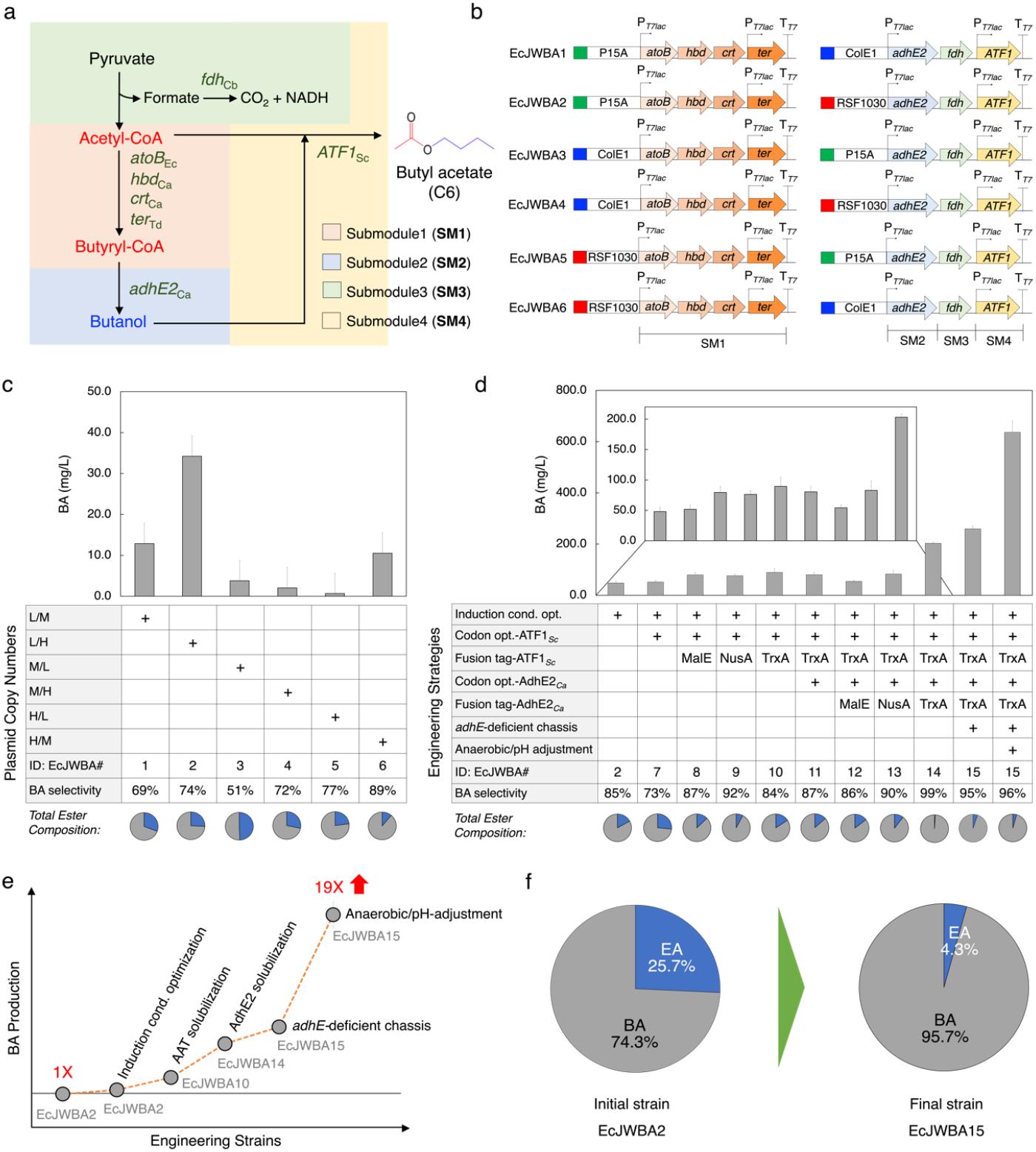
*De novo* microbial biosynthesis of butyl acetate (BA) production from glucose. (**a**) Modular biosynthesis pathway of BA. (**b**) Schematic of six initial strains carrying BA production modules with different copy numbers. The copy number of origins of replication are as follows: P15A (in green), ~10; ColE1 (in blue), ~40; RSF1030 (in red), ~100 (Lee and Trinh, 2019). (**c-f**) *De novo* BA production from glucose. Endogenous BA production in (**c**) six initial strains (EcJWBA1~EcJWBA6). (**d**) Improved BA production in EcJWBA2~EcJWBA15. (**e**) Summary of BA production. (**f**) Comparison of BA selectivity between initial (EcJWBA2) and final (EcJWBA15) strains. For pie charts, EA is shown in blue and BA in gray. Error bars represent the standard deviation of at least two biological replicates. Abbreviations: cond.: conditions, opt.: optimization.

**Figure 3.**
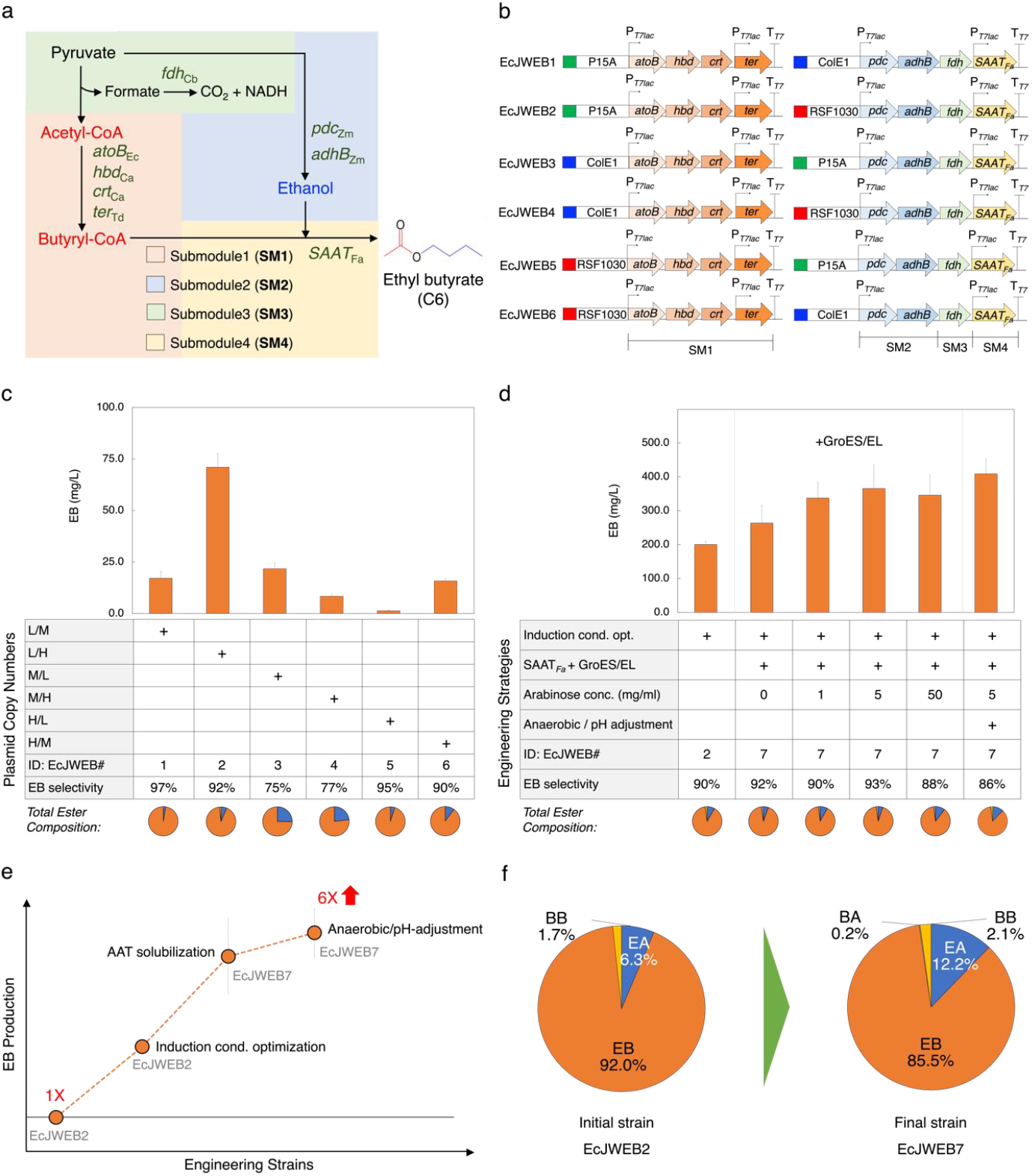
*De novo* microbial biosynthesis of ethyl butyrate (EB) production from glucose. (**a**) Modular biosynthesis pathway of EB. (**b**) Schematic of six initial strains carrying EB production modules with different copy numbers. The copy number of origins of replication are as follows: P15A (in green), ~10; ColE1 (in blue), ~40; RSF1030 (in red), ~100 (Lee and Trinh, 2019). (**c-f**) *De novo* EB production from glucose. Endogenous EB production in (**c**) six initial strains (EcJWEB1~EcJWEB6). (**d**) Improved EB production in EcJWEB7; (**e**) Summary of EB production. (**f**) Comparison of EB selectivity between initial (EcJWEB2) and final (EcJWEB7) strains. For pie charts, EA is shown in blue, BA in gray, EB in orange, and BB in yellow. Error bars represent the standard deviation of at least two biological replicates. Abbreviations: cond.: conditions, opt.: optimization.

**Table 1.**
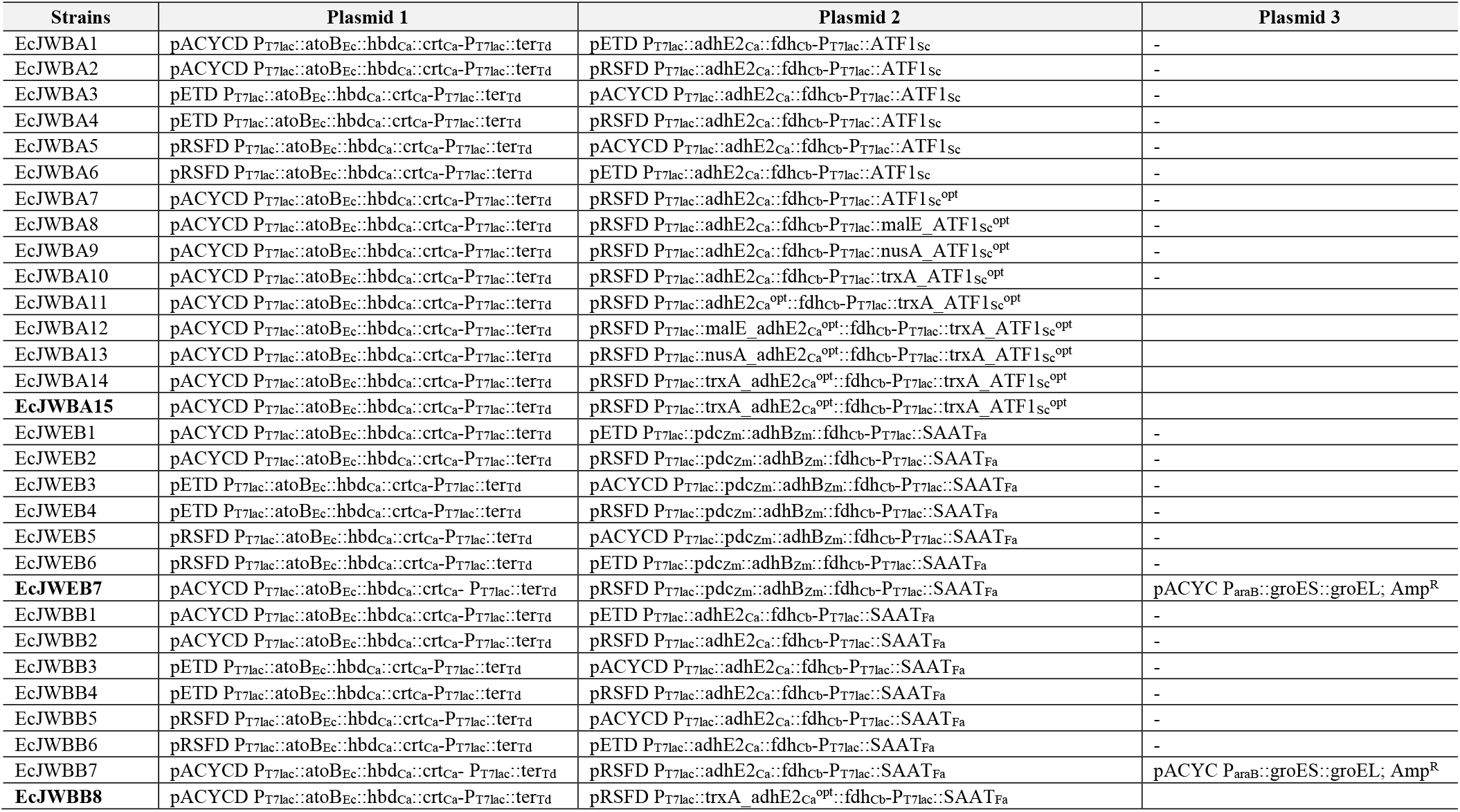
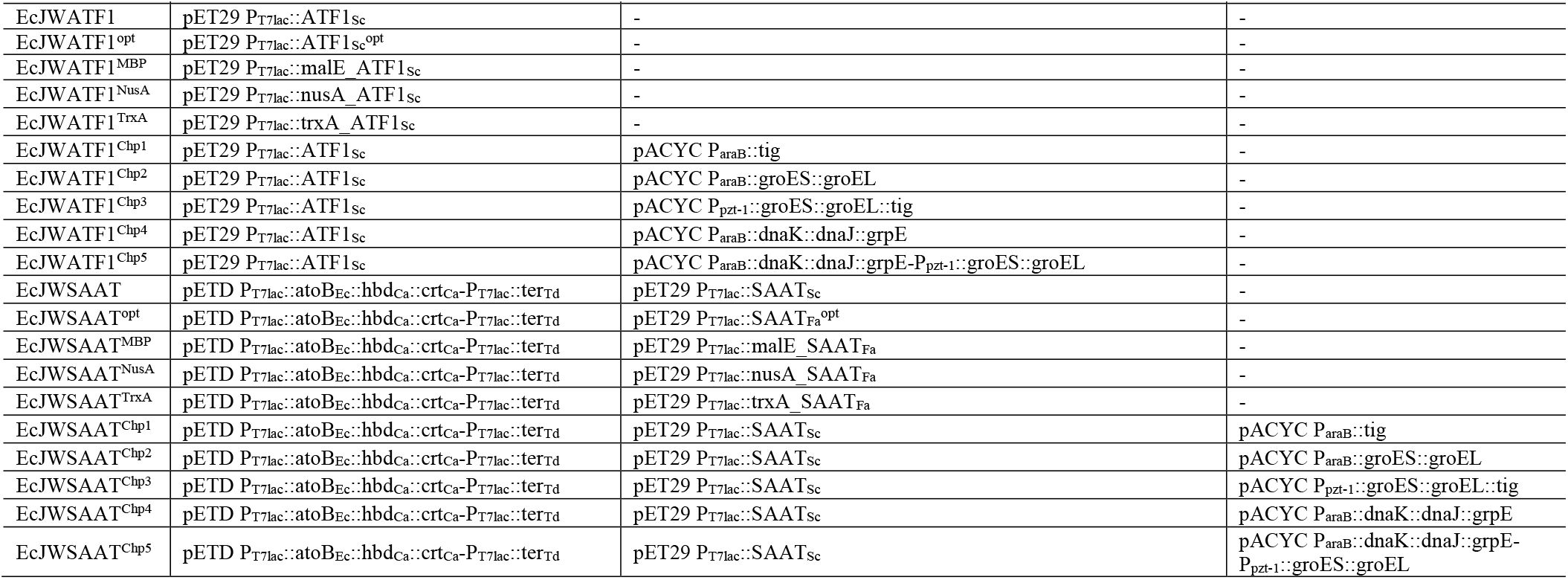
A list of strains and plasmids used in this study. Except for EcJWBA15, TCS083 *ΔfadE* (DE3) (Layton and Trinh, 2014) was used as a host strain. For EcJWBA15, TCS095 (DE3) (Wilbanks et al., 2018) was used as a host strain. Key strains are in bold.

### Establishing the *de novo* microbial biosynthesis of designer butyryl-CoA-derived esters in initial strains

Following the construction of initial strains, we characterized them in capped conical tubes to validate the constructed biosynthesis pathways of butyryl-CoA-derived esters and to identify the best combination of pCore and pDerivatization. Our results show that these pathways are functional in the chassis cell as evidenced by the protein expression via SDS-PAGE analysis (Fig. S1) and production of the target products (Figs. 2c, 3c, and 4c). We found that EcJWBA2 (Fig. 2c), EcJWEB2 (Fig. 3c), and EcJWBB2 (Fig. 4c) carrying the pCore with low copy number and the pDerivatization with high copy number achieved the highest ester production among the six initial strains characterized for each compound, indicating that higher alcohol production and/or AAT expression are required for efficient ester synthesis. For BA production, EcJWBA2 produced 34.2 ± 6.6 mg/L of BA with the selectivity of 74.3% (Fig. 2c, Table S4). For EB production, EcJWEB2 produced 71.0 ± 6.6 mg/L of EB with the selectivity of 92.0% (Fig. 3c, Table S5). For BB production, EcJWBB2 produced 33.5 ± 2.9 mg/L of BB with the selectivity of 20.8% (Fig. 4c, Table S6).

**Figure 4.**
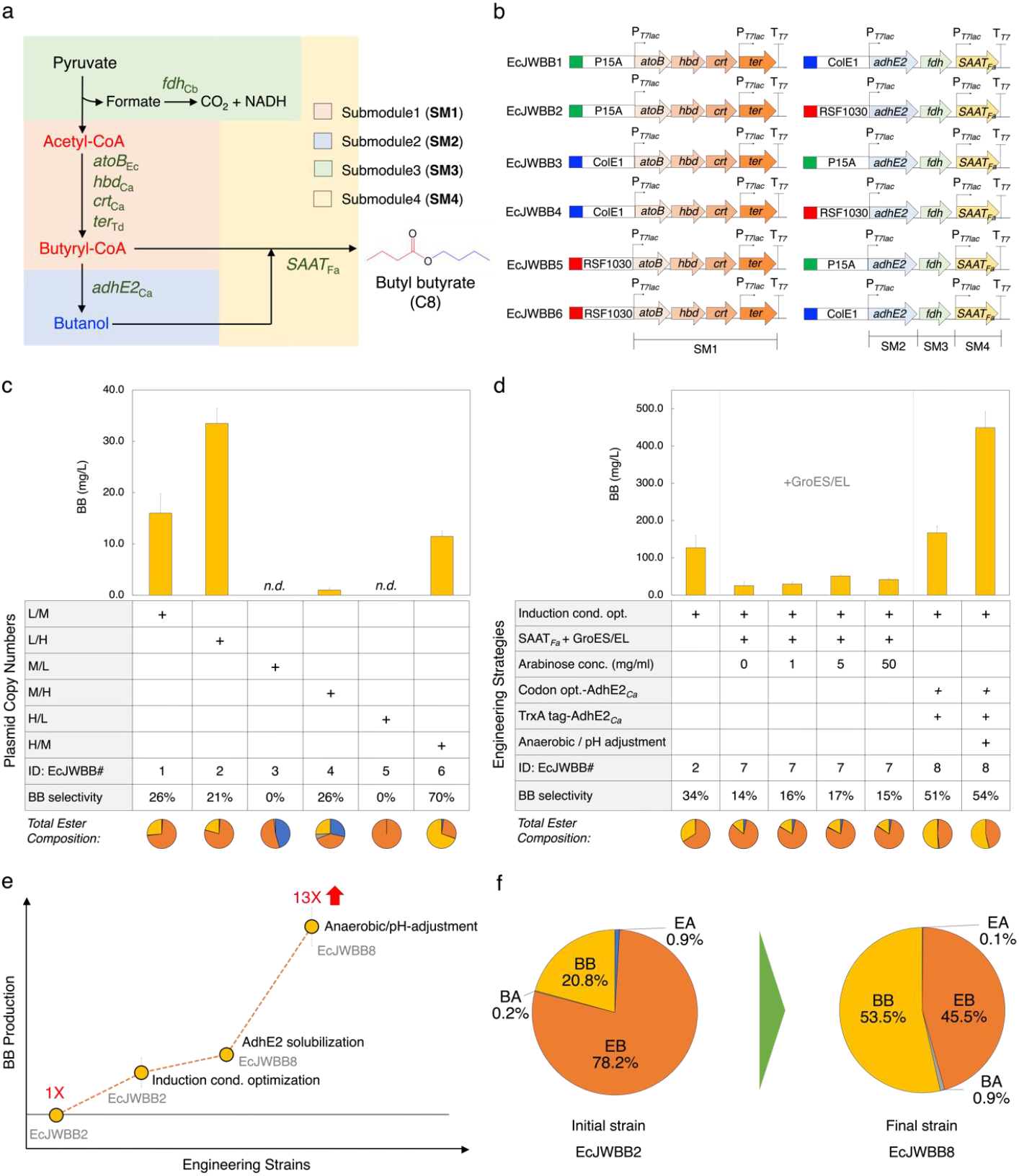
*De novo* microbial biosynthesis of butyl butyrate (BB) production from glucose. (**a**) Modular biosynthesis pathway of BB. (**b**) Schematic of initial six strains carrying BB production modules with different copy numbers. The copy number of origins of replication are as follows: P15A (in green), ~10; ColE1 (in blue), ~40; RSF1030 (in red), ~100 (Lee and Trinh, 2019). (**c-f**) *De novo* BB production from glucose. Endogenous BB production in (**c**) six initial strains (EcJWBB1~EcJWBB6). (**d**) Improved BB production in EcJWBB7~EcJWBB8. (**e**) Summary of BB production. (**f**) Comparison of BB selectivity between initial (EcJWBB2) and final (EcJWBB8) strains. For pie charts, EA is shown in blue, BA in gray, EB in orange, and BB in yellow. Error bars represent the standard deviation of at least two biological replicates. Abbreviations: cond.: conditions, opt.: optimization, *n.d.*: not detected.

After validating the synthetic pathways of butyryl-CoA-derived esters, we next optimized induction conditions with the best identified ester producers, EcJWBA2, EcJWEB2, and EcJWBB2. To optimize induction conditions, we tested various induction conditions using a combination of two different temperatures (28°C and 37°C), and three different concentrations of the inducer (0.01, 0.1, and 1.0 mM IPTG). The results show that the titer of BA, EB, and BB was improved by 1.4, 2.8, and 3.8-fold at the optimized induction conditions, respectively (Figs. 2d, 3d, 4d, and S2, Table S7-S9). Specifically, for BA production, EcJWBA2 produced 48.0 ± 7.1 mg/L of BA with the selectivity of 83.1% when it was induced by 0.1 mM of IPTG at 28°C (Figs. 2d, and S2a, Table S7). For EB production, EcJWEB2 produced 200.4 ± 9.4 mg/L of EB with the selectivity of 89.6% when it was induced by 0.1 mM of IPTG at 28°C (Figs. 3d, and S2b, Table S8). For BB production, EcJWBB2 produced 127.4 ± 32.5 mg/L of BB with the selectivity of 34.0% when it was induced by 0.1 mM of IPTG at 37°C (Figs. 4c, and S2c, Table S9). Collectively, we established the *de novo* microbial biosynthesis of butyryl-CoA-derived esters and identified the induction conditions for enhanced production of these esters to be used in the subsequent experiments. The ester production and selectivity, however, are relatively low, especially for the biosynthesis of EB and BB, and hence require further optimization.

### Alleviating poor AAT expression as a rate limiting step for ester microbial biosynthesis through a comprehensive evaluation of various protein solubilization strategies

Since the protein bands of ATF1_Sc_ and SAAT_Fa_ are weaker than the other protein bands (Figs. S1, and S3) and these eukaryotic AATs are prone to poor expression in *E. coli* (Tai et al., 2015), we hypothesized that the AAT flux is one of the rate limiting steps and hence improving soluble expression of AATs would enhance the ester production. To examine the effect of AAT solubilization on ester production, we chose three strategies including i) codon optimization (Gorochowski et al., 2015; Rosano and Ceccarelli, 2009); ii) the use of fusion partners such as maltose binding protein (MBP) (Waugh, 2016), N-utilization substrate A (NusA) (Raran-Kurussi and Waugh, 2014), or thioredoxin 1 (TrxA) (Lavallie et al., 1993); and iii) co-expression of molecular chaperones (DnaK/DnaJ/GrpE, GroES/GroEL, or Trigger factor (Tf)) (Thomson et al., 2013) (Fig. 5a).

**Figure 5.**
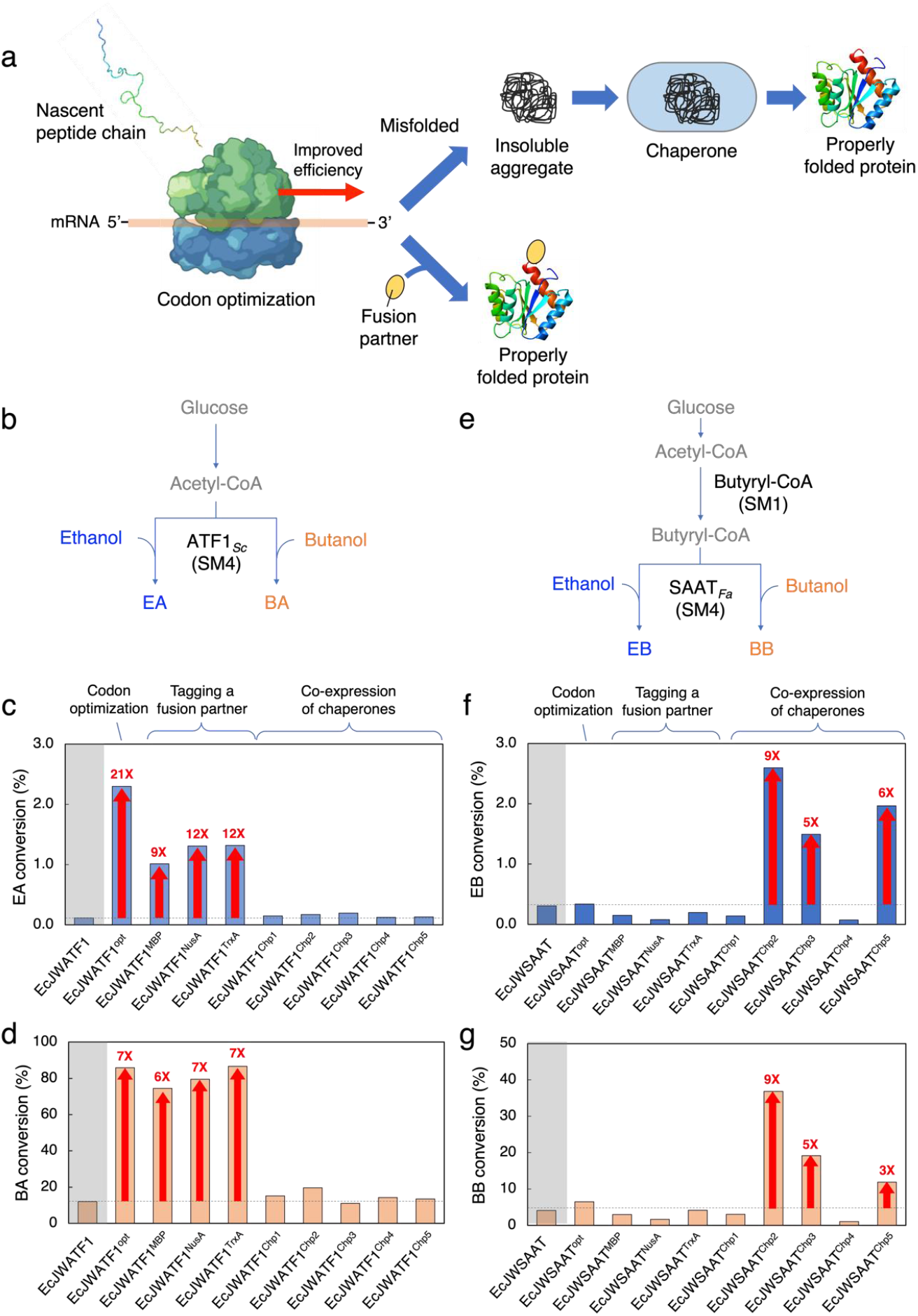
Protein solubilization of AATs. (**a**) Schematic presentation of protein solubilization strategies used in this study. (**b-d**) Bioconversion of an alcohol into an ester by ATF1_Sc_ derivatives. (**b**) Proposed bioconversion pathway of an alcohol (ethanol/butanol) into an ester (ethyl acetate (EA)/butyl acetate (BA)) by ATF1*_sc_* in *E. coli.* Conversion of (**c**) ethanol into EA, and (**d**) butanol into BA in engineered *E. coli.* (**e-g**) Bioconversion of an alcohol into an ester by SAAT_Fa_ derivatives. (**e**) Proposed bioconversion pathway of an alcohol (ethanol/butanol) into an ester (ethyl butyrate (EB)/butyl butyrate (BB)) by SAAT*_Fa_* in *E. coli.* Conversion of (**b**) ethanol into EB and (**c**) butanol into BB in the engineered *E. coli*. Grey box indicates a negative control. Error bars represent the standard deviation of three biological replicates. Each ester conversion (%) was calculated by (target ester produced)/(target ester produced + corresponding alcohol substrate remained)*100 (mole/mole).

To test whether AAT is a rate limiting step in isolation, we engineered the chassis cell harboring only the acyl-CoA and AAT submodules while alcohols can be externally doped. We started by generating the plasmids that harbor wildtype AATs, codon optimized AATs, fusion partner tagged AATs. For BA production, the plasmids carrying wildtype *ATF1*_Sc_, codon optimized *ATF1*_Sc_ (*ATF1*_Sc_^opt^), and N’-terminus MBP-, NusA-, or TrxA-tagged ATF1_Sc_ *(malE_ATF1*_Sc_, *nusA_ATF1*_Sc_, or *trxA_ATF1*_Sc_) were constructed and introduced into TCS083 *ΔfadE* (DE3) (Layton and Trinh, 2014), resulting EcJWATF1, EcJWATF1^opt^, EcJWATF1^MBP^, EcJWATF1^NusA^, EcJWATF1^TrxA^, respectively (Table 1). For EB and BB production, the plasmids carrying wildtype SAAT (*SAAT*_Fa_), codon optimize SAAT_Fa_ (*SAAT*_Fa_^opt^). and N’-terminus MBP-, NusA-, or TrxA-tagged SAAT_Fa_ (*malE_SAAT*_Fa_; *nusA_SAAT*_Fa_; or *trxA_SAAT*_Fa_) were constructed and introduced into TCS083 *ΔfadE* (DE3) (Layton and Trinh, 2014) with the pACYCDuet-1 carrying the SM1 (butyryl-CoA pathway), resulting EcJWSAAT, EcJWSAAT^opt^, EcJWSAAT^MBP^, EcJWSAAT^NusA^, EcJWSAAT^Trx^A, respectively (Table 1). To co-express chaperones with AATs, the chaperone plasmid set comprising of five different plasmids carrying various chaperones were introduced into EcJWATF1 and EcJWSAAT, resulting in EcJWATF1^Chp1^~EcJWATF1^Chp5^ and EcJWSAAT^Chp1^~EcJWSAAT^Chp5^, respectively (Table 1).

We next characterized the engineered strains in conical tubes with 2 g/L of alcohol doping including ethanol and butanol to evaluate the conversion of an alcohol (ethanol/butanol) into an ester (EA/BA) by ATF1_Sc_ (Fig. 5b) or EB/BB by SAAT_Fa_ (Fig. 5e), respectively. The cultures were induced by 0.1 mM of IPTG, 0.5 mg/ml of L-arabinose (if applicable) and/or 5 ng/ml of tetracycline (if applicable), and the protein expressions were confirmed by SDS-PAGE analysis (Fig. S4). The characterization results show that the protein solubilization strategies enhanced the conversion of an alcohol into an ester (Fig. 5c-d, and 5f-g). Interestingly, we found that different protein solubilization strategies worked effectively for different AATs. In particular, the codon optimization and use of a fusion partner (i.e., MBP, NusA, or TrxA) for ATF1_Sc_ improved the BA conversion while co-expression of chaperones (i.e., GroES/EL, GroES/EL/Tf, or DnaK/DnaJ/GrpE/GroES/EL) with SAATFa enhanced the EB/BB conversion.

For BA conversion, EcJWATF1^opt^, EcJWATF1^MBP^, EcJWATF1^NusA^, EcJWATF1^TrxA^ achieved 85.9%, 74.6%, 79.5%, and 86.6% of BA conversion, resulting in 7.1, 6.2, 6.6, and 7.2-fold improvement as compared to EcJWATF1 (12.0%), respectively (Fig. 5d, and Table S10). With 2 g/L butanol doping, the BA production could reach up to 2.26 ± 0.22 g/L, and the selectivity of BA over other esters was as high as 98.1%. As compared to BA production under similar characterization conditions, improvement in EB production was less prominent, only reaching up to 0.46 ± 0.09 g/L with a selectivity of 86.3%. In particular, EcJWSAAT^Chp2^, EcJWSAAT^Chp3^, and EcJWSAAT^Chp5^ achieved 2.6%, 1.5%, and 2.0% of EB conversion, leading to 8.5, 4.7, and 6.4-fold improvement as compared to EcJWSAAT (0.3%), respectively (Fig. 5f, and Table S11). The metabolic burden in protein expressions and different catalytic efficiency between SAATFa and ATF1_Sc_ likely contributed to the differences in strain performance. For BB conversion, the BB production was reasonably high, reaching up to 1.71 ± 0.26 g/L with a selectivity of 79.2%. In particular, EcJWSAAT^Chp2^, EcJWSAAT^Chp3^, and EcJWSAAT^Chp5^ achieved 36.8%, 19.1%, and 11.9% of BB conversion, resulting in 9.0, 4.7, and 2.9-fold improvement as compared to EcJWSAAT (4.1%), respectively (Fig. 5g, and Table S11). Even though ATF1_Sc_ and SAATFa have specificity towards longer-chain alcohols and acyl-CoAs, respectively, we also observed production of EA as a minor byproduct. For the EA conversion, EcJWATF1^opt^, EcJWATF1^MBP^, EcJWATF1^NusA^, EcJWATF1^TrxA^ achieved 2.3%, 1.0%, 1.3%, and 1.3% of EA conversion, resulting in 20.5, 9.0, 11.7, and 11.8-fold improvement as compared to EcJWATF1 (0.1%), respectively (Fig. 5c, and Table S10).

Overall, our results clearly indicated that AAT solubilization plays a critical role in controlling ester production and selectivity by isolated investigation of the AAT submodule with alcohol doping experiments. The next step is to evaluate the effect of most solubilized AATs on the *de novo* microbial biosynthesis of designer esters directly from fermentable sugars.

### Enhancing AAT solubility improves the endogenous production of BA and EB but not BB

To evaluate whether the AAT solubilization improves the *de novo* ester microbial biosynthesis from glucose, we constructed and characterized various BA, EB, and BB-producing strains. For BA production, we first built four pRSFDuet-1 plasmids carrying SM2(*adhE2*_Ca_)-SM3(/*dh*^opt^)-SM4(*ATF1*_Sc_^opt^, *malE_ATF1*_Sc_^opt^, *nusA_ATF1*_Sc_^opt^, or *trxA_ATF1*_Sc_^opt^), respectively (Table S1), and then introduced them into the chassis cell TCS083 *ΔfadE* (DE3) with the pACYCDuet-1 plasmid carrying the SM1 (butyryl-CoA pathway) to generate EcJWBA7~EcJWBA10, respectively (Table 1). For EB/BB production, we additionally introduced the plasmid carrying *groES* and *groEL* into EcJWEB2 and EcJWBB2, resulting in EcJWEB7 and EcJWBB7, respectively (Table 1). We characterized the engineered ester production strains in conical tubes for the endogenous ester production from glucose. For the expression of chaperones in EcJWEB2 and EcJWBB2, we also tested three different concentrations of L-arabinose as an inducer.

The characterization results show that the ATF1_Sc_ solubilization indeed enhanced the endogenous BA production. Specifically, EcJWBA7~EcJWBA10 produced 51.7 ± 7.1 mg/L, 79.3 ± 9.8 mg/L, 76.1 ± 6.2 mg/L, and 89.5 ± 14.8 mg/L of BA, resulting in 1.1, 1.7, 1.6, and 1.9-fold improved BA production as compared to the EcJWBA2 (48.0 ± 7.1 mg/L), respectively (Fig. 2d, and Table S12). Notably, because EcJWBA8~10 expressing ATF1_Sc_^opt^ with N’-terminus fusion partner such as MBP, NusA, and TrxA achieved higher BA production than that of EcJWBA7 expressing ATF1_Sc_^opt^ alone, we could confirm that there is a synergistic effect between codon optimization and the use of a fusion partner in BA production with ATF1_Sc_.

Similarly, we also observed the improvement in the endogenous EB production. When the cell cultures were induced by 0 mg/ml, 0.1 mg/ml, 0.5 mg/ml and 5.0 mg/ml of L-arabinose, respectively, EcJWEB7 produced 263.9 ± 51.8 mg/L, 337.6 ± 46.1 mg/L, 365.7 ± 69.2 mg/L, and 346.2 ± 59.8 mg/L of EB, resulting in 1.3, 1.7, 1.8, and 1.7-fold improved EB production as compared to EcJWEB2 (200.4 ± 9.4 mg/L) (Fig. 3d, and Table S13). However, unlike the EB production, chaperone expression negatively affected the endogenous BB production. When the cultures were induced by 0 mg/ml, 0.1 mg/ml, 0.5 mg/ml and 5.0 mg/ml of L-arabinose, respectively, EcJWBB7 produced 25.6 ± 10.7 mg/L, 30.0 ± 4.8 mg/L, 51.4 ± 3.4 mg/L, and 42.1 ± 4.1 mg/L of BB, achieving 0.2, 0.2, 0.4, and 0.3-fold decreased BB production as compared to EcJWBB2 (127.4 ± 32.5 mg/L) (Fig. 4d, and Table S14). Overall, the AAT solubilization is critical for the *de novo* microbial biosynthesis of designer esters. However, the limitation of other enzymatic steps besides the AAT condensation might interfere with the ester biosynthesis due to the complexity of the engineered pathways.

### Co-solubilization of AdhE2_Ca_ and AAT improved the endogenous production of BA and BB

Due to the low residual butanol in our BA/BB production experiments, we hypothesized that the low availability of butanol, one of the intermediates for butyl esters synthesis (Table S4-S14), might have affected the endogenous production of BB and EB. The bi-functional aldehyde/alcohol dehydrogenase AdhE2_Ca_ is known for its critical role in butanol production, and its low solubility can significantly reduce *in vivo* activities as compared to the *in vitro* activities (Shen et al., 2011). We tested whether the co-solubilization of AdhE2_Ca_ and AAT improved the *de novo* microbial biosynthesis of BA/BB by alleviating the limitation of butanol.

For BA production, we first constructed four pRSFDuet-1 plasmids carrying SM2(*adhE2*_Ca_^opt^, *malE_adhE2*_Ca_^opt^, *nwsA*_*adhE2*_Ca_^opt^, or *trxA*_*adhE2*_Ca_^opt^)-SM3*fdh*^opt^)-SM4(*trxA*_*ATF1*_Sc_^opt^), respectively (Table S1) and introduced them into the chassis cell TCS083 *ΔfadE* (DE3) with the pACYCDuet-1 plasmid carrying the SM1 (butyryl-CoA pathway) to generate EcJWBA11~EcJWBA14, respectively (Table 1). Next, we characterized these strains in conical tubes for BA production. The expression of the pathway enzymes was confirmed by SDS-PAGE analysis (Fig. S5). The results show EcJWBA11~EcJWBA14 produced 80.3 ± 9.0 mg/L, 54.2 ± 4.9 mg/L, 82.8 ± 15.5 mg/L, and 203.0 ± 5.7 mg/L of BA, respectively (Fig. 2d, Table S15). Remarkably, EcJWBA14 achieved 2.3-fold improved BA production (203.0 ± 5.7 mg/L) as compared to EcJWBA10 (89.5 ± 14.8 mg/L), indicating that solubilization of pathway enzymes using a fusion partner can be a simple, but useful extendable pathway optimization strategy in metabolic engineering.

To strengthen this result, we also evaluated whether the use of TrxA fusion partner with AdhE2_Ca_^opt^ can improve BB production. We built the pRSFDuet-1 plasmid carrying SM2(*trxA*_*adhE2*_Ca_^opt^)-SM3(*fdh*^opt^)-SM4(*SAAT*_Fa_) (Table S1) and introduced it into the chassis cell TCS083 *ΔfadE* (DE3) with the pACYCDuet-1 plasmid carrying the SM1 (butyryl-CoA pathway) to generate EcJWBB8 (Table 1). By characterizing EcJWBB8 in conical tubes, the results, indeed, show that EcJWBB8 achieved 1.3-fold improved BB production (167.3 ± 18.2 mg/L) as compared to EcJWBB2 (127.4 ± 32.5 mg/L) (Fig. 4d, and Table S15). Notably, EcJWBB8 achieved ~1.5-fold improved BB selectivity (50.6%) as compared to EcJWBB2 (34.0%), resulting in ~1.7-fold improved butanol/ethanol ratio (g/g of butanol to ethanol) (from 0.04 to 0.07) and ~0.6-fold reduced EB production (from 246.2 ± 72.6 mg/L to 156.3 ± 22.4 mg/L) (Tables S15). This result suggests that there is a substrate competition between ethanol and butanol in the enzymatic reaction of ATF1Sc, which can be alleviated by either engineering AATs with alcohol substrate preference or tuning the selective alcohol production.

Overall, the results highlight the critical limitation of AdhE and AAT enzymatic steps negatively affects the designer ester biosynthesis due to poor enzyme expression. Combinational solubilization of multiple pathway enzymes is feasible to alleviate the enzyme expression of a large, complex metabolic pathway.

### Anaerobic conditions helped boost the endogenous production of butyryl-CoA-derived designer esters

Although BA and BB production were improved to some extent via co-solubilization of AdhE2_Ca_ and AAT, residual butanol titer was still low, remaining at the titers of 0.18 ± 0.00 g/L for EcJWBB8 and 0.20 ± 0.01 g/L for EcJWBA14 (Tables S9, and S15). Given that the abundant alcohol production is important for ester synthesis due to the high *K*M value of AATs (Lee and Trinh, 2019; Tai et al., 2015), butanol production needs to be further improved for higher production of butyl esters. Because strict anaerobic conditions are important for alcohol production (Bond-Watts et al., 2011; Shen et al., 2011), we characterized the final strains, EcJWBA14, EcJWEB7, and EcJWBB8, in anaerobic bottles with pH-adjustment to evaluate their performance in production of C4-dereived esters. The culture pH was adjusted to around 7 with 10 M NaOH every 24 hours to maintain the optimum growth pH of *E. coli* (Philip et al., 2018).

The characterization results of EcJWBA14, EcJWEB7, EcJWBB8 showed 12.9, 5.8, and 13.4-fold improvement in titers, 4.8, 3.7, and 4.6-fold improvement in yields, and 6.5, 1.4, and 3.4- fold improvement in productivity as compared to the initial strains, EcJWBA2, EcJWEB2, and EcJWBB2, respectively. Specifically, EcJWBA14 produced 441.4 ± 40.9 mg/L of BA (9.2% of maximum theoretical yield) with 91.7% of selectivity (Figs. 2d, and Table S16), EcJWEB7 produced 408.9 ± 44.3 mg/L of EB (8.5% of maximum theoretical yield) with 85.5% of selectivity (Figs. 3d, 3f, and Table S16), and EcJWBB8 produced 449.6 ± 43.0 mg/L of BB (10.0% of maximum theoretical yield) with 53.5% of selectivity (Figs. 4d, 4f, and Table S16). In comparison with the direct fermentative production of butyryl-CoA-derived esters by *E. coli* in previous studies (Layton and Trinh, 2014), EcJWBA14 achieved 882.8, 1839.0, and 3937.0-fold improved TRY (titer, productivity, and yield) in BA production, EcJWEB7 achieved 3.1, 3.1, and 11.0-fold improved TRY in EB production, and EcJWBB8 achieved 12.2, 12.2, and 44.1-fold improved TRY in BB production (Table S18).

### Use of an endogenous *adhE-deficient* chassis further enhanced BA production

The modular cell TCS083 *ΔfadE* (DE3) is designed to be auxotrophic and required to metabolically couple with a butyryl-CoA-derived ester module (Layton and Trinh, 2014; Trinh et al., 2015). We hypothesized the promiscuity of endogenous alcohol dehydrogenases might have interfered with the butyryl-CoA-derived ester modules, competing for ester biosynthesis. For instance, the endogenous bifunctional aldehyde/alcohol dehydrogenase *adhE* favors the formation of ethanol over butanol (Atsumi et al., 2008). To demonstrate the optimization of BA production, we replaced TCS083 *ΔfadE* (DE3) with TCS095 (DE3) that is an *adhE*-deficient chassis cell (Wilbanks et al., 2018). We generated EcJWBA15 by introducing the BA pathway into TCS095 (DE3) (Table 1). The characterization results of EcJWBA15 in conical tubes showed that EcJWBA15 achieved higher BA production than EcJWBA14 by 1.28-fold with a titer of 259.5 ± 11.6 mg/L and a selectivity of 94.8 (Fig. 2d and Table S17). Finally, by characterizing EcJWBA15 in anaerobic bottles with pH-adjustment, we could achieve 636.3 ± 44.8 mg/L of BA (23.0% of maximum theoretical yield) with a high selectivity (95.7%) (Figs. 2d, 2f, and Table S17).

## Discussion

In this study, we reported the development of a generalizable framework to engineer a modular microbial platform for anaerobic production of butyryl-CoA-derived esters from fermentable sugars. Using the modular design approach, each ester production strain can be generated from an engineered modular (chassis) cell and an exchangeable ester production module in a plug-and-play fashion. The study focused on engineering exchangeable ester production modules to be compatible with the chassis cell for efficient biosynthesis of designer esters with controllable selectivity, including BA, EB, and BB. To build these modules, we arranged a set of 11 heterologous genes, derived from bacteria, yeasts, and plants, into four submodules SM1-SM4 to facilitate rapid module construction and optimization via manipulation of gene replication, transcription, translation, post-translation, pathway enzymes, and pathway induction conditions (Fig. S7). Our modular cell engineering approach achieves the highest production of esters (i.e., BA, EB, and BB) ever reported in *E. coli* with controllable selectivity.

For the past two decades, controlling selectivity of designer esters has been an outstanding metabolic engineering problem, mainly due to the complexity of the engineered pathways that require simultaneous expression of multiple heterologous enzymes causing deficient supply of precursor metabolites (i.e., alcohols and acyl-CoAs) for ester condensation. While the metabolic pathways directed towards biosynthesis of acetyl-CoA, butyryl-CoA, ethanol, and butanol are well known and can be tuned by manipulating gene replication (i.e., plasmid copy numbers) and transcription (e.g., RBSs, promoters) in many native and engineered ethanol/butanol producers (Nielsen et al., 2009; Shen et al., 2011; Sillers et al., 2008; Trinh et al., 2008; Zhang et al., 1995), extension of these pathways for ester biosynthesis has been problematic due to poor AAT expression and specificity. Aiming at these critical issues in this study, our initial combinatorial strategies to control the selectivity of butyryl-CoA-derived ester biosynthesis are proven to be effective by using ATF1_Sc_ specific for acetate ester biosynthesis (e.g., BA) and SAAT_Fa_ specific for butyrate ester biosynthesis (e.g., EB and BB) (Layton and Trinh, 2016a), together with optimization of the pathway gene replication and transcription for sufficient supply of precursor metabolites. However, the ester titer and selectivity were still insufficient since the problem of proper expression of pathway enzymes remained, which is difficult to solve. Based on the Protein-Sol, a web tool for predicting protein solubility from sequence (Hebditch et al., 2017), AATs are predicted to have the lowest solubility among the engineered pathway enzymes followed by AdhE2_Ca_ (Fig. S6). The prediction is consistent with the SDS-PAGE analysis as observed in our study (Fig. S4) and by others (Zhu et al., 2015), explaining the production phenotypes of the engineered strains (Figs. 2e, 3e, and 4e).

In solving the AAT expression problem, we found that implementing a comprehensive screening of protein solubilization strategies including codon optimization (Gorochowski et al., 2015; Rosano and Ceccarelli, 2009), the use of fusion tags (Lavallie et al., 1993; Raran-Kurussi and Waugh, 2014; Waugh, 2016), co-expression of chaperones (Thomson et al., 2013), and/or the combination thereof is simple and effective. Remarkably, fusion tags improve ATF1_Sc_ solubilization while chaperones enhance expression of SAAT_Fa_, which is intriguing to discover but is not trivial to predict or explain. In general, we expect that solubilization with fusion tags are enzyme specific; however, use of chaperones alone can be very unspecific and might not be as effective, especially when multiple enzymes are expressed simultaneously. Our study highlights the significance of modulating the translation and post-translation for multiple pathway enzymes, that cannot be effectively addressed by optimization of gene replication and transcription alone as commonly practiced in the fields of metabolic engineering and synthetic biology. We demonstrated the combinatorial protein solubilization strategy can be a powerful tool to improve microbial production of biochemicals and biofuels with (eukaryotic) aggregate-prone enzymes in the bacterial chassis cells like *E. coli*.

The engineered strains achieved 19-fold in BA production with 96% selectivity, 6-fold in EB production with 86% selectivity, and 13-fold in BB production with 54% selectivity, as compared to the initial strains. Unlike the microbial biosynthesis of BA and EB, tuning the BB selectivity is intrinsically challenging due to the following reasons: i) the butanol biosynthesis is limiting due to low solubility of AdhE_Ca_ and ii) AdhE_Ca_ is promiscuous and can reduce both acetyl-CoA and butyryl-CoA (Shen et al., 2011). While our strategy to enhance co-solubilization of AdhE_Ca_ together with SAAT_Fa_ helped improve BB production and selectivity, EB is always produced as a significant byproduct. Should high selectivity be desirable for specific applications, two engineering strategies can be further exploited to overcome this problem: i) improving specificity of AdhE_Ca_ towards butyryl-CoA and ii) decoupling butanol and butyl butyrate production using a microbial co-culture system. Furthermore, without external supply of butanol, production of BA and BB directly from glucose was much lower likely due to metabolic burden required for expressing multiple pathway enzymes.

One distinct advantage of microbial production of esters is that they have low solubility in an aqueous phase and hence are very beneficial for fermentation. Even though the butyryl-CoA-derived esters are inhibitory to microbes (Wilbanks and Trinh, 2017), their toxicity is significantly alleviated by implementing *in situ* fermentation and extraction (Layton and Trinh, 2014) (Rodriguez et al., 2014; Tai et al., 2015). Besides beneficial detoxification by extraction, we also found that maintaining anaerobic culture conditions at neutral pH control improves ester production. Anaerobic production of butyryl-CoA-derived esters from fermentable sugars are favorable because i) high product yields can be achieved due to higher reduction of esters than glucose and ii) scale-up for anaerobic processes is much simpler and more economical (Layton and Trinh, 2014).

In conclusion, we developed a generalizable framework to engineer a modular microbial platform for anaerobic production of butyryl-CoA-derived designer esters. Using the principles of modular design, we engineered the *de novo* modular fermentative pathways of biosynthesis of BA, EB, and BB from fermentable sugars in *E. coli* with controllable selectivity. In addition to the conventional strategies of replication and transcription manipulation, implementing various protein solubilization strategies on aggregate-prone pathway enzymes to control enzyme (post)-translation is very crucial to enhance ester production and selectivity. We envision the modular microbial ester synthesis platform presented is expected to accelerate the biosynthesis of diverse natural esters with various industrial applications.

## Methods

### Strains and plasmids

The list of strains and plasmids used in this study are presented in Table 1. Briefly, *E. coli* TOP10 strain was used for molecular cloning. Except for EcJWBA15, TCS083 *ΔfadE* (DE3) (Layton and Trinh, 2014) was used as a host strain. For EcJWBA15, TCS095 (DE3) (Wilbanks et al., 2018) was used as a host strain. A set of duet vectors including pACYCDuet-1, pETDuet-1, and pRSFDuet-1 was used as plasmid backbones for constructing a library of BA, EB, and BB production modules. The codon-optimized *S. cerevisiae ATF1* (ATF1_Sc_^opt^), cultivated strawberry *(F. ananassa) SAAT* (SAAT_Fa_^opt^), *Candida boidinii fdh* (fdh_Cb_^opt^), and *C. acetobutylicum adhE2* (adhE2_Ca_^opt^) were synthesized by the U.S. Department of Energy (DOE) Joint Genome Institute (JGI). The list of codon optimized gene sequences is presented in Table S3.

### Culture conditions

For molecular cloning and seed cultures, lysogeny broth (LB) was used. For ester production, TBD_50_ medium, terrific broth (TB) with 50 g/L glucose was used (without supplementation with glycerol). For all cultures, 30 μg/mL chloramphenicol (Cm), 50 μg/mL kanamycin (Kan), and/or 100 μg/mL ampicillin (Amp) were added to the medium where applicable.

For seed cultures, 1% (v/v) of stock cells were grown overnight in 5 mL of LB medium with appropriate antibiotics. For ester production in capped conical tubes, seed cultures were prepared as described in seed cultures. About 1% (v/v) of seed cultures were inoculated in 500 mL baffled flasks containing 50 ml of TBD_50_ medium with appropriate antibiotics. The cells were aerobically grown in shaking incubators at 28°C or 37°C, 200 rpm and induced at an O.D._600_ of 0.6~0.8 with various concentrations of IPTG, arabinose (if applicable), and/or 5 ng/ml of tetracycline (if applicable). After 2 hours of induction, the cultures in the baffled flasks were distributed into 15 mL conical centrifuge tubes (Cat. #339650, Thermo Scientific, MA, USA) with a working volume of 5 mL. Then, each tube was overlaid with 1 mL hexadecane (20% (v/v)) for *in situ* ester recovery and capped to generate anaerobic conditions. Finally, the tubes were grown for another 18 hours on a 75° angled platform in shaking incubators at 28°C or 37°C, 200 rpm. The remained cultures in the baffled flasks were induced for further 2 hours and then the cells were harvested for SDS-PAGE analysis.

For ester production in strict anaerobic bottles with pH-adjustment, the induced cultures were prepared as described in ester production in conical tubes with a working volume of 100 mL. To generate the anaerobic state, the induced cultures were transferred into anaerobic bottles. Then, each anaerobic bottle was overlaid with 20% (v/v) of hexadecane for *in situ* ester recovery and sealed with a rubber stopper inside the anaerobic chamber. The headspace of the anaerobic bottles was vacuumed and replaced by an anaerobic mix of 90% N_2_, 5% H_2_, and 5% CO_2_ inside the anaerobic chamber. Finally, the anaerobic bottles were grown for another 90 hours in shaking incubators at 28°C or 37°C, 200 rpm. The culture medium and hexadecane overlay samples were taken through the rubber stopper via a syringe and needle by maintaining the ratio of 5:1. The culture pH was adjusted to around 7 using 10 M NaOH every 24 hours.

### Protein expression and SDS-PAGE analysis

The cells were collected from the culture by centrifugation and resuspended in 1X PBS (Phosphate Buffered Saline) buffer (pH 7.4) at the final O.D._600_ of 10. Cell pellets were disrupted using the B-PER complete reagent (Cat. #89822, Thermo Scientific, MA. USA), according to the manufacturer’s instruction. Total and soluble fractions were separated by centrifugation for 20 min at 4°C. The resulting samples were mixed with 6X SDS (sodium dodecyl sulfate) sample buffer, heated at 95°C for 5 min, and analyzed by SDS-PAGE (SDS-polyacrylamide gel electrophoresis) using Novex™ 14% Tris-Glycine protein gels (Cat. #XP00145BOX, Thermo Scientific, MA, USA). Protein bands were visualized with Coomassie Brilliant Blue staining.

### Determination of cell concentrations

The optical density was measured at 600 nm using a spectrophotometer (GENESYS 30, Thermo Scientific, IL, USA). The dry cell mass was obtained by multiplication of the optical density of culture broth with a pre-determined conversion factor, 0.48 g/L/O.D.

### High performance liquid chromatography (HPLC)

Metabolites and doped alcohols were quantified by using the Shimadzu HPLC system (Shimadzu Inc., MD, USA) equipped with the Aminex HPX-87H cation exchange column (BioRad Inc., CA, USA) heated at 50°C. A mobile phase of 10 mN H2SO4 was used at a flow rate of 0.6 mL/min. Detection was made with the reflective index detector (RID).

### Gas chromatography coupled with mass spectroscopy (GC/MS)

All esters were quantified by GC/MS. For GC/MS analysis, the hexadecane overlays were used for quantification of esters. To prepare samples, the hexadecane overlays were first centrifuged at 4,800 x g for 5 min and diluted with hexadecane containing internal standard (isoamyl alcohol) in a 1:1 (v/v) ratio. Then, 1 μL of samples were directly injected into a gas chromatograph (GC) HP 6890 equipped with the mass selective detector (MS) HP 5973 using an autosampler. For the GC system, helium was used as the carrier gas at a flow rate of 0.5 mL/min and the analytes were separated on a Phenomenex ZB-5 capillary column (30 m x 0.25 mm x 0.25 μm). The temperature of the oven was programmed from the initial value of 50°C followed by a heating rate of 1°C/min to 58°C and then heated at a ramp of 25°C/min to 235°C. Finally, the oven is heated to 300°C a ramp of 50°C/min and held for 2 min. The injection was performed using the splitless mode with an initial injector temperature of 280°C. For the MS system, a selected ion monitoring (SIM) mode was deployed to detect analytes. The SIM parameters for detecting esters were as follows: i) for ethyl acetate, ions 45.00, and 61.00 detected from 4.15 to 5.70 min, ii) for isoamyl alcohol (internal standard), ions 45.00, and 88.00 detected from 5.70 to 7.20 min, iii) for ethyl butyrate, ions 47.00, and 116.00 detected from 7.20 to 7.75 min, iv) for butyl acetate, ions 61.00, and 116.00 detected from 7.75 to 11.25 min, vi) for butyl butyrate, ions 101.00, and 116.00 detected from 11.25 to 12.50 min.

## Acknowledgments

This research was financially supported in part by the NSF CAREER award (NSF#1553250) and the DOE subcontract grant (DE-AC05-000R22725) by the Center of Bioenergy Innovation, the U.S. Department of Energy Bioenergy Research Center funded by the Office of Biological and Environmental Research in the DOE Office of Science, and the U.S. Department of Energy Joint Genome Institute. The authors would like to thank the Center of Environmental Biotechnology at UTK for using the GC/MS instrument.

## Author contributions

CTT conceived and supervised this study. JWL and CTT designed the experiments, analyzed the data, and drafted the manuscript. JWL performed the experiments. Both authors read and approved the final manuscript.

