## Supplementary material for "Controlling Selectivity of Modular Microbial Biosynthesis of Butyryl-CoA-Derived Designer Esters": Table S1-S18, Figures S1-S7

**Supplementary Table S1.** A list of plasmids used in this study. The copy number of duet vectors are as follows: pACYCDuet-1, ~10; pETDuet-1, ~40; pRSFDuet-1, ~100[4]. Abbreviations: F, forward; R, reverse; BB, backbone; MCS, multiple cloning site.

| **Plasmid** | **Primers (5'→3')** | **Cloning strategies** |
|---|---|---|
| ***Submodule1 (SM1): Butyryl-CoA synthesis*** | | |
| pACYCDuet-1 $P_{T7lac}$::atoB$_{Ec}$::hbd$_{Ca}$::crt$_{Ca}$-$P_{T7lac}$::ter$_{Td}$ | CrCoA_F: CAGCAGCCATCACCATCATCACCACAGCCAGGATCCGATGAAAAATTGTGTCATCGTCAG<br>CrCoA_R: CTATCTATTTTTGAAGCCTTCAATTT<br>MCS1_MCS2 linker_F: GCTTTCATAGAGAAAAGAAAAATTGAAGGCTTCAAAAATAGATAGGAATTCGAGCTCGGC<br>MCS1_MCS2 linker_R: ATTGTTCCTAACCATTGGTTTTACAATCATCATATGTATATCTCCTTCTTATACTTAACT<br>ter$_{Td}$_F: AGTTAAGTATAAGAAGGAGATATACATATGATGATTGTAAAACCAATGGTTAGG<br>ter$_{Td}$_R: TCAAATTTCGCAGCAGCGGTTTCTTTACCAGACTCGAGTTAAATCCTGTCGAACCTTTCT<br>BB_Duet_F: CGGATCCTGGCTGTGG<br>BB_Duet_R: CTCGAGTCTGGTAAAGAAACC | Biosynthesis pathway of Butyryl-CoA in pDL2[5] was subcloned into the plasmids with various copy numbers by Gibson Assembly method[6]. Specifically, atoB$_{Ec}$- hbd$_{Ca}$-crt$_{Ca}$ and Ter$_{Td}$ were subcloned into the MCS1and MCS2 of the duet vectors, respectively. |
| pETDuet-1 $P_{T7lac}$::atoB$_{Ec}$::hbd$_{Ca}$::crt$_{Ca}$-$P_{T7lac}$::ter$_{Td}$ | | |
| pRSFDuet-1 $P_{T7lac}$::atoB$_{Ec}$::hbd$_{Ca}$::crt$_{Ca}$-$P_{T7lac}$::ter$_{Td}$ | | |
| ***Submodule2/3/4 (SM2-SM3-SM4) for BA synthesis: Butanol+NADH+ATF1*** | | |
| pACYCDuet-1 $P_{T7lac}$::adhE2$_{Ca}$::fdh$_{Cb}^{opt}$-$P_{T7lac}$::ATF1$_{Sc}$ | adhE2$_{Ca}$_F: GCCATCACCATCATCACCACAGCCAGGATCCGATGAAAGTTACAAATCAAAAAGAACTAA<br>adhE2$_{Ca}$_R: TATATCTCCTTTTAAAATGATTTTATATAGATATCCTTAAGTTCAC<br>fdh$_{Cb}^{opt}$_F: AGGATATCTATATAAAATCATTTTAAAAGGAGATATAATGAAAATTGTGCTGGTTTTGTA<br>fdh$_{Cb}^{opt}$_R: TTGTCGACCTGCAGGCGCGCCGAGCTCGAATTCTTATTTTTTATCGTGTTTACCATACGC<br>MCS1_MCS2 linker_BA_F: TAAGAATTCGAGCTCGGC<br>MCS1_MCS2 linker_BA_R: GATTTCATTCATCATATGTATATCTCCTTCTTATACTTAACT<br>ATF1$_{Sc}$_F: ATATTAGTTAAGTATAAGAAGGAGATATACATATGATGAATGAAATCGATGAGAAAAATC<br>ATF1$_{Sc}$_R: CGTTCAAATTTCGCAGCAGCGGTTTCTTTACCAGACTCGAGCTAAGGGCCTAAAAGGAGA<br>BB_Duet_F: CGGATCCTGGCTGTGG<br>BB_Duet_R: CTCGAGTCTGGTAAAGAAACC | Five PCR fragments including i) adhE2$_{Ca}$ from gDNA of *C. acetobutylicum*, ii) fdh$_{Cb}^{opt}$ from pET29 $P_{T7lac}$::fdh$_{Cb}^{opt}$, iii) MCS1_MCS2 linker from pACYCDuet-1, iv) ATF1$_{Sc}$ from , and v) plasmid backbone with various copy numbers were assembled by Gibson Assembly method[6]. Specifically, adhE2$_{Ca}$-fdh$_{Cb}$ and ATF1$_{Sc}$ were subcloned into the MCS1and MCS2 of the duet vectors, respectively. |
| pETDuet-1 $P_{T7lac}$::adhE2$_{Ca}$::fdh$_{Cb}^{opt}$-$P_{T7lac}$::ATF1$_{Sc}$ | | |
| pRSFDuet-1 $P_{T7lac}$::adhE2$_{Ca}$::fdh$_{Cb}^{opt}$-$P_{T7lac}$::ATF1$_{Sc}$ | | |
| pRSFDuet-1 $P_{T7lac}$::adhE2$_{Ca}$::fdh$_{Cb}^{opt}$-$P_{T7lac}$::ATF1$_{Sc}^{opt}$ | ATF1$_{Sc}^{opt}$_F: ATGAACGAAATCGACGAAAAAAATCAG<br>ATF1$_{Sc}^{opt}$_R: CGGACCCAGTAACAAAGCTTTATA<br>BB_BA_ATF1$_{Sc}^{opt}$_F: GAGCTTTGTAGCATATATAAAGCTTTGTTACTGGGTCCGCTCGAGTCTGGTAAAGAAAC<br>BB_BA_ATF1$_{Sc}^{opt}$_R: GCGCCTGATTTTTTTCGTCGATTTCGTTCATCATATGTATATCTCCTTCTTATACTTAAC | ATF1$_{Sc}$ in pRSFDuet-1 $P_{T7}$::adhE2$_{Ca}$::fdh$_{Cb}^{opt}$-$P_{T7}$::ATF1$_{Sc}$ was replaced with ATF1$_{Sc}^{opt}$ from pET29 $P_{T7}$::ATF1$_{Sc}^{opt}$ by Gibson Assembly method[6]. |

**Supplementary Table S1.** (Continued)

| **Plasmid** | **Primers (5'→3')** | **Cloning strategies** |
|---|---|---|
| pRSFDuet-1<br>P$_{T7lac}$::adhE2$_{Ca}$::fdh$_{Cb}^{opt}$-<br>P$_{T7lac}$::malE_ATF1$_{Sc}^{opt}$ | malE_ATF1$_{Sc}^{opt}$_F:<br>ATATTAGTTAAGTATAAGAAGGAGATATACATATGATGAAAATCGAAGAAGGTAAACTGG<br>malE_ATF1$_{Sc}^{opt}$_R:<br>CCTGCTGAACTGGCGCCTGATTTTTTTCGTCGATTTCGTTCTTGGTGATACGAGTCTGC<br>BB_BA_ FP_ATF1$_{Sc}^{opt}$_F:<br>AACGAAATCGACGAAAAAAATCAG<br>BB_BA_FP_ATF1$_{Sc}^{opt}$_R:<br>CATATGTATATCTCCTTCTTATACTTAACTAATATACT | The start codon of ATF1$_{Sc}^{opt}$ in pRSFDuet-1 P$_{T7}$::adhE2$_{Ca}$::fdh$_{Cb}^{opt}$-P$_{T7}$::ATF1$_{Sc}^{opt}$ was replaced with the *malE* from gDNA of *E. coli* MG1655 by Gibson Assembly method[6]. |
| pRSFDuet-1<br>P$_{T7lac}$::adhE2$_{Ca}$::fdh$_{Cb}^{opt}$-<br>P$_{T7lac}$::nusA_ATF1$_{Sc}^{opt}$ | nusA_ATF1$_{Sc}^{opt}$_F:<br>ATATTAGTTAAGTATAAGAAGGAGATATACATATGATGAACAAAGAAATTTTGGCTGTAG<br>nusA_ATF1$_{Sc}^{opt}$_R:<br>CACTCCTGCTGAACTGGCGCCTGATTTTTTTCGTCGATTTCGTTCGCTTCGTCACCGAAC<br>BB_BA_ FP_ATF1$_{Sc}^{opt}$_F:<br>AACGAAATCGACGAAAAAAATCAG<br>BB_BA_FP_ATF1$_{Sc}^{opt}$_R:<br>CATATGTATATCTCCTTCTTATACTTAACTAATATACT | The start codon of ATF1$_{Sc}^{opt}$ in pRSFDuet-1 P$_{T7}$::adhE2$_{Ca}$::fdh$_{Cb}^{opt}$-P$_{T7}$::ATF1$_{Sc}^{opt}$ was replaced with the *nusA* from gDNA of *E. coli* MG1655 by Gibson Assembly method[6]. |
| pRSFDuet-1<br>P$_{T7lac}$::adhE2$_{Ca}$::fdh$_{Cb}^{opt}$-<br>P$_{T7lac}$::trxA_ATF1$_{Sc}^{opt}$ | trxA_ATF1$_{Sc}^{opt}$_F:<br>TATTAGTTAAGTATAAGAAGGAGATATACATATGATGAGCGATAAAATTATTCACCTGAC<br>trxA_ATF1$_{Sc}^{opt}$_R:<br>CTGCTGAACTGGCGCCTGATTTTTTTCGTCGATTTCGTTCGCCAGGTTAGCGTC<br>BB_BA_ FP_ATF1$_{Sc}^{opt}$_F:<br>AACGAAATCGACGAAAAAAATCAG<br>BB_BA_FP_ATF1$_{Sc}^{opt}$_R:<br>CATATGTATATCTCCTTCTTATACTTAACTAATATACT | The start codon of ATF1$_{Sc}^{opt}$ in pRSFDuet-1 P$_{T7}$::adhE2$_{Ca}$::fdh$_{Cb}^{opt}$-P$_{T7}$::ATF1$_{Sc}^{opt}$ was replaced with the *trxA* from gDNA of *E. coli* MG1655 by Gibson Assembly method[6]. |
| pRSFDuet-1<br>P$_{T7lac}$::adhE2$_{Ca}^{opt}$::fdh$_{Cb}^{opt}$-<br>P$_{T7lac}$::trxA_ATF1$_{Sc}^{opt}$ | adhE2$_{Ca}^{opt}$_F:<br>ATGAAAGTGACCAATCAAAAAGAGC<br>adhE2$_{Ca}^{opt}$_R:<br>TTAGAAACTCTTTATGTAAATGTCTTTGAGTTCG<br>BB_BA_adhE2$_{Ca}^{opt}$_F:<br>CAAAGACATTTACATAAAGAGTTTCTAAAAGGAGATATAATGAAAATTGTGCTGGTTTTG<br>BB_BA_adhE2$_{Ca}^{opt}$_R:<br>GTTCAGTTTCTGTTTCAGCTCTTTTTGATTGGTCACTTTCATCGGATCCTGGCTGTGG | adhE2$_{Ca}$ in pRSFDuet-1 P$_{T7}$::adhE2$_{Ca}$::fdh$_{Cb}^{opt}$-P$_{T7}$::trxA_ATF1$_{Sc}$ was replaced with adhE2$_{Ca}^{opt}$ from pET29 P$_{T7}$:: adhE2$_{Ca}^{opt}$ by Gibson Assembly method[6]. |
| pRSFDuet-1<br>P$_{T7lac}$::malE_adhE2$_{Ca}^{opt}$::fdh$_{Cb}^{opt}$-<br>$_{T7lac}$::trxA_ATF1$_{Sc}^{opt}$ | malE_adhE2$_{Ca}^{opt}$_F:<br>GCAGCCATCACCATCATCACCACAGCCAGGATCCGATGAAAATCGAAGAAGGTAAACTGG<br>malE_adhE2$_{Ca}^{opt}$_R:<br>CGTTCAGTTTCTGTTTCAGCTCTTTTTGATTGGTCACTTTCTTGGTGATACGAGTCTGC<br>BB_BA_FP_adhE2$_{Ca}^{opt}$_F:<br>CGGATCCTGGCTGTGG<br>BB_BA_FP_adhE2$_{Ca}^{opt}$_R:<br>AAAGTGACCAATCAAAAAGAGCTG | The start codon of adhE2$_{Ca}^{opt}$ in pRSFDuet-1 P$_{T7lac}$::adhE2$_{Ca}^{opt}$::fdh$_{Cb}^{opt}$-P$_{T7lac}$::trxA_ATF1$_{Sc}^{opt}$ was replaced with the *malE* from gDNA of *E. coli* MG1655 by Gibson Assembly method[6]. |
| pRSFDuet-1<br>P$_{T7lac}$::nusA_adhE2$_{Ca}^{opt}$::fdh$_{Cb}^{opt}$-<br>$_{T7lac}$::trxA_ATF1$_{Sc}^{opt}$ | nusA_adhE2$_{Ca}^{opt}$_F:<br>GCAGCCATCACCATCATCACCACAGCCAGGATCCGATGAACAAAGAAATTTTGGCTGTAG<br>nusA_adhE2$_{Ca}^{opt}$_R:<br>CGTTCAGTTTCTGTTTCAGCTCTTTTTGATTGGTCACTTTCGCTTCGTCACCGAAC<br>BB_BA_FP_adhE2$_{Ca}^{opt}$_F:<br>CGGATCCTGGCTGTGG<br>BB_BA_FP_adhE2$_{Ca}^{opt}$_R:<br>AAAGTGACCAATCAAAAAGAGCTG | The start codon of adhE2$_{Ca}^{opt}$ in pRSFDuet-1 P$_{T7lac}$::adhE2$_{Ca}^{opt}$::fdh$_{Cb}^{opt}$-P$_{T7lac}$::trxA_ATF1$_{Sc}^{opt}$ was replaced with the *nusA* from gDNA of *E. coli* MG1655 by Gibson Assembly method[6]. |
| pRSFDuet-1<br>P$_{T7lac}$::trxA_adhE2$_{Ca}^{opt}$::fdh$_{Cb}^{opt}$-<br>P$_{T7lac}$::trxA_ATF1$_{Sc}^{opt}$ | trxA_adhE2$_{Ca}^{opt}$_F:<br>CAGCCATCACCATCATCACCACAGCCAGGATCCGATGAGCGATAAAATTATTCACCTGAC<br>trxA_adhE2$_{Ca}^{opt}$_R:<br>CGTTCAGTTTCTGTTTCAGCTCTTTTTGATTGGTCACTTTCGCCAGGTTAGCGTC<br>BB_BA_FP_adhE2$_{Ca}^{opt}$_F:<br>CGGATCCTGGCTGTGG<br>BB_BA_FP_adhE2$_{Ca}^{opt}$_R:<br>AAAGTGACCAATCAAAAAGAGCTG | The start codon of adhE2$_{Ca}^{opt}$ in pRSFDuet-1 P$_{T7lac}$::adhE2$_{Ca}^{opt}$::fdh$_{Cb}^{opt}$-P$_{T7lac}$::trxA_ATF1$_{Sc}^{opt}$ was replaced with the *trxA* from gDNA of *E. coli* MG1655 by Gibson Assembly method[6]. |

**Supplementary Table S1.** (Continued)

| **Plasmid** | **Primers (5'→3')** | **Cloning strategies** |
|---|---|---|
| ***Submodule2/3/4 (SM2-SM3-SM4) for EB synthesis: Ethanol+NADH+SAAT*** | | |
| pACYCDuet-1<br>P$_{T7lac}$::pdc$_{Zm}$::adhB$_{Zm}$::fdh$_{Cb}^{opt}$-P$_{T7lac}$::SAAT$_{Fa}$ | EtOH_F:<br>GGCAGCAGCCATCACCATCATCACCACAGCCAGGATCCGATGAGTTATACTGTCGGTACC<br>EtOH_R:<br>CGCATCATACAAAACCAGCACAATTTTCATTATATCTCCTTTTAGAAAGCGCTCAGGAAG<br>fdh$_{Cb}^{opt}$_F:<br>ATGAAAATTGTGCTGGTTTTGTAATGAAAATTGTGCTGGTTTTGTA<br>fdh$_{Cb}^{opt}$_R:<br>TTGTCGACCTGCAGGCGCGCCGAGCTCGAATTCTTATTTTTTATCGTGTTTACCATACGC<br>MCS1_MCS2 linker_F:<br>AAAATAAGAATTCGAGCTCGGC<br>MCS1_MCS2 linker_R:<br>GGAATTTATACTGACCTCAATTTTCTCCATCATATGTATATCTCCTTCTTATACTTAACT<br>SAAT$_{Fa}$_F:<br>AGTTAAGTATAAGAAGGAGATATACATATGATGGAGAAAATTGAGGTCAGTATA<br>SAAT$_{Fa}$_R:<br>AAATTTCGCAGCAGCGGTTTCTTTACCAGACTCGAGTTAAATTAAGGTCTTTGGAGATGC<br>BB_Duet_F:<br>CGGATCCTGGCTGTGG<br>BB_Duet_R:<br>CTCGAGTCTGGTAAAGAAACC | Five PCR fragments including i) pdc$_{Zm}$-adhB$_{Zm}$ from pCT24[5], ii) fdh$_{Cb}^{opt}$ from pET29 P$_{T7lac}$::fdh$_{Cb}^{opt}$, iii) MCS1_MCS2 linker from pACYCDuet-1, iv) SAAT$_{Fa}$ from pDL1[5], and v) plasmid backbone with various copy numbers were assembled by Gibson Assembly method[6]. Specifically, pdc$_{Zm}$-adhB$_{Zm}$-fdh$_{Cb}$ and SAAT$_{Fa}$ were subcloned into the MCS1and MCS2 of the duet vectors, respectively. |
| pETDuet-1<br>P$_{T7lac}$::pdc$_{Zm}$::adhB$_{Zm}$::fdh$_{Cb}^{opt}$-P$_{T7lac}$::SAAT$_{Fa}$ | | |
| pRSFDuet-1<br>P$_{T7lac}$::pdc$_{Zm}$::adhB$_{Zm}$::fdh$_{Cb}^{opt}$-P$_{T7lac}$::SAAT$_{Fa}$ | | |
| ***Submodule2/3/4 (SM2-SM3-SM4) for BB synthesis: Butanol+NADH+SAAT*** | | |
| pACYCDuet-1<br>P$_{T7lac}$::adhE2$_{Ca}$::fdh$_{Cb}^{opt}$-P$_{T7lac}$::SAAT$_{Fa}$ | adhE2$_{Ca}$_F:<br>GCCATCACCATCATCACCACAGCCAGGATCCGATGAAAGTTACAAATCAAAAAGAACTAA<br>adhE2$_{Ca}$_R:<br>TATATCTCCTTTTAAAATGATTTTATATAGATATCCTTAAGTTCAC<br>BB_Duet_F:<br>CGGATCCTGGCTGTGG<br>BB_BB_R:<br>AGGATATCTATATAAAATCATTTTAAAAGGAGATATAATGAAAATTGTGCTGGTTTTGTA | pdc$_{Zm}$-adhB$_{Zm}$ in EB production modules were replaced with adhE2$_{Ca}$ from gDNA of *C. acetobutylicum* by Gibson Assembly method[6]. |
| pETDuet-1<br>P$_{T7lac}$::adhE2$_{Ca}$::fdh$_{Cb}^{opt}$-P$_{T7lac}$::SAAT$_{Fa}$ | | |
| pRSFDuet-1<br>P$_{T7lac}$::adhE2$_{Ca}$::fdh$_{Cb}^{opt}$-P$_{T7lac}$::SAAT$_{Fa}$ | | |
| pRSFDuet-1<br>P$_{T7lac}$::trxA_adhE2$_{Ca}^{opt}$::fdh$_{Cb}^{opt}$-P$_{T7lac}$::SAAT$_{Fa}$ | SAAT$_{Fa}$_F:<br>AGTTAAGTATAAGAAGGAGATATACATATGATGGAGAAAATTGAGGTCAGTATA<br>SAAT$_{Fa}$_R:<br>AAATTTCGCAGCAGCGGTTTCTTTACCAGACTCGAGTTAAATTAAGGTCTTTGGAGATGC<br>BB_trxA_adhE2$_{Ca}^{opt}$_F:<br>ATGTATATCTCCTTCTTATACTTAACTAATATACTAAGATGG<br>BB_Duet_R:<br>CTCGAGTCTGGTAAAGAAACC | trxA_ATF1$_{Sc}^{opt}$ in pRSFDuet-1 P$_{T7lac}$::trxA_adhE2$_{Ca}^{opt}$::fdh$_{Cb}^{opt}$-P$_{T7lac}$::trxA_ATF1$_{Sc}^{opt}$ was replaced with the SAAT$_{Fa}$ from pET29 P$_{T7lac}$::SAAT$_{Fa}$ by Gibson Assembly method[6]. |
| ***Wildtype AATs*** | | |
| pET29 P$_{T7lac}$::ATF1$_{Sc}$ | ATF1$_{Sc}$_F:<br>ATAATTTTGTTTAACTTTAAGAAGGAGATATAGATATGAATGAAATCGATGAGAAAAAT<br>ATF1$_{Sc}$_R:<br>TTTGTTAGCAGCCGGATCTCAGTGGTGGTGGTGGTGGTGGATAGGGCCTAAAAGGAGAG<br>BB_pET29_F:<br>ATCCACCACCACCACC<br>BB_pET29_R:<br>ATCTATATCTCCTTCTTAAAGTTAAACAAAATTATTTCTAG | ATF1$_{Sc}$ in pDL004[7] was subcloned into pET29 by Gibson Assembly method[6]. |

**Supplementary Table S1.** (Continued)

| Plasmid | Primers (5'→3') | Cloning strategies |
|---|---|---|
| pET29 P$_{T7lac}$::SAAT$_{Fa}$ | SAAT$_{Fa}$_F:<br>TAGAAATAATTTTGTTTAACTTTAAGAAGGAGATATAGATATGG<br>AGAAAATTGAGGTCAG<br>SAAT$_{Fa}$_R:<br>GTTAGCAGCCGGATCTCAGTGGTGGTGGTGGTGGTGGATAATTA<br>AGGTCTTTGGAGATGC<br>BB_pET29_F:<br>ATCCACCACCACCACC<br>BB_pET29_R:<br>ATCTATATCTCCTTCTTAAAGTTAAACAAAATTATTTCTAG | SAAT$_{Fa}$ in pDL001[7] was subcloned into pET29 by Gibson Assembly method[6]. |
| ***Codon optimized genes by the U.S. Department of Energy (DOE) Joint Genome Institute (JGI)*** | | |
| pET29 P$_{T7lac}$::ATF1$_{Sc}^{opt}$ | univ_F:<br>CCTCTAGAAATAATTTTGTTTAACTTTAAGAAGGAGA<br>univ_R:<br>CGGATCTCAGTGGTGGTGGT | The plasmids were constructed by JGI. The synthesized genes were amplified and cloned into pET29 plasmid using the EcoRV restriction site. The genes were codon optimized to *E. coli*. |
| pET29 P$_{T7lac}$::SAAT$_{Fa}^{opt}$ | | |
| pET29 P$_{T7lac}$::fdh$_{Cb}^{opt}$ | | |
| pET29 P$_{T7lac}$::adhE2$_{Ca}^{opt}$ | | |
| ***AATs with fusion partners*** | | |
| pET29 P$_{T7lac}$::malE_ATF1$_{Sc}$ | malE_ATF1$_{Sc}$_F:<br>TTTGTTTAACTTTAAGAAGGAGATATAGATATGAAAATCGAAGA<br>AGGTAAACTGGTAATC<br>malE_ATF1$_{Sc}$_R:<br>CTTGTTGCACGGGGGCCTGATTTTTCTCATCGATTTCATTCTTGG<br>TGATACGAGTCTGCG<br>BB_pET29 ATF1$_{Sc}$_F:<br>AATGAAATCGATGAGAAAAATCAGGC<br>BB_pET29 ATF1$_{Sc}$_R:<br>ATCTATATCTCCTTCTTAAAGTTAAACAAAATTATTTCTAG | The start codon of ATF1$_{Sc}$ in pET29 P$_{T7}$::ATF1$_{Sc}$ was replaced with malE from gDNA of *E. coli* MG1655 by Gibson Assembly method[6]. |
| pET29 P$_{T7lac}$::nusA_ATF1$_{Sc}$ | nusA_ATF1$_{Sc}$_F:<br>ATTTTGTTTAACTTTAAGAAGGAGATATAGATATGAACAAAGAA<br>ATTTTGGCTGTAGTTG<br>nusA_ATF1$_{Sc}$_R:<br>ATTCTTGTTGCACGGGGGCCTGATTTTTCTCATCGATTTCATTCG<br>CTTCGTCACCGAACC<br>BB_pET29 ATF1$_{Sc}$_F:<br>AATGAAATCGATGAGAAAAATCAGGC<br>BB_pET29 ATF1$_{Sc}$_R:<br>ATCTATATCTCCTTCTTAAAGTTAAACAAAATTATTTCTAG | The start codon of ATF1$_{Sc}$ in pET29 P$_{T7}$::ATF1$_{Sc}$ was replaced with nusA from gDNA of *E. coli* MG1655 by Gibson Assembly method[6]. |
| pET29 P$_{T7lac}$::trxA_ATF1$_{Sc}$ | trxA_ATF1$_{Sc}$_F:<br>ATTTTGTTTAACTTTAAGAAGGAGATATAGATATGAGCGATAAA<br>ATTATTCACCTGACTG<br>trxA_ATF1$_{Sc}$_R:<br>CATTCTTGTTGCACGGGGGCCTGATTTTTCTCATCGATTTCATTC<br>GCCAGGTTAGCGTCG<br>BB_pET29 ATF1$_{Sc}$_F:<br>AATGAAATCGATGAGAAAAATCAGGC<br>BB_pET29 ATF1$_{Sc}$_R:<br>ATCTATATCTCCTTCTTAAAGTTAAACAAAATTATTTCTAG | The start codon of ATF1$_{Sc}$ in pET29 P$_{T7}$::ATF1$_{Sc}$ was replaced with trxA from gDNA of *E. coli* MG1655 by Gibson Assembly method[6]. |
| pET29 P$_{T7lac}$::malE_SAAT$_{Fa}$ | malE_ATF1$_{Sc}$_F:<br>TTTGTTTAACTTTAAGAAGGAGATATAGATATGAAAATCGAAGA<br>AGGTAAACTGGTAATC<br>malE_ATF1$_{Sc}$_R:<br>TGATGGTGTGTTTGGAATTTATACTGACCTCAATTTTCTCCTTGG<br>TGATACGAGTCTGCG<br>BB_pET29 SAAT$_{Fa}$_F:<br>GAGAAAATTGAGGTCAGTATAAATTCCAAAC<br>BB_pET29 SAAT$_{Fa}$_R:<br>ATCTATATCTCCTTCTTAAAGTTAAACAAAATTATTTCTAG | The start codon of SAAT$_{Fa}$ in pET29 P$_{T7}$::SAAT$_{Fa}$ was replaced with malE from gDNA of *E. coli* MG1655 by Gibson Assembly method[6]. |

**Supplementary Table S1.** (Continued)

| **Plasmid** | **Primers (5'→3')** | **Cloning strategies** |
|---|---|---|
| pET29 P$_{T7lac}$::nusA_SAAT$_{Fa}$ | nusA_ATF1$_{Sc}$_F:<br>ATTTTGTTTAACTTTAAGAAGGAGATATAGATATGAACAAAGAAATTTTGGCTGTAGTTG<br>nusA_ATF1$_{Sc}$_R:<br>GTTTGATGGTGTGTTTGGAATTTATACTGACCTCAATTTTCTCCGCTTCGTCACCGAACC<br>BB_pET29 SAAT$_{Fa}$_F:<br>GAGAAAATTGAGGTCAGTATAAATTCCAAAC<br>BB_pET29 SAAT$_{Fa}$_R:<br>ATCTATATCTCCTTCTTAAAGTTAAACAAAATTATTTCTAG | The start codon of SAAT$_{Fa}$ in pET29 P$_{T7}$::SAAT$_{Fa}$ was replaced with nusA from gDNA of *E. coli* MG1655 by Gibson Assembly method[6]. |
| pET29 P$_{T7lac}$::trxA_SAAT$_{Fa}$ | trxA_ATF1$_{Sc}$_F:<br>ATTTTGTTTAACTTTAAGAAGGAGATATAGATATGAGCGATAAAATTATTCACCTGACTG<br>trxA_ATF1$_{Sc}$_R:<br>GGTTTGATGGTGTGTTTGGAATTTATACTGACCTCAATTTTCTCCGCCAGGTTAGCGTCG<br>BB_pET29 SAAT$_{Fa}$_F:<br>GAGAAAATTGAGGTCAGTATAAATTCCAAAC<br>BB_pET29 SAAT$_{Fa}$_R:<br>ATCTATATCTCCTTCTTAAAGTTAAACAAAATTATTTCTAG | The start codon of SAAT$_{Fa}$ in pET29 P$_{T7}$::SAAT$_{Fa}$ was replaced with trxA from gDNA of *E. coli* MG1655 by Gibson Assembly method[6]. |
| ***Co-expression of chaperones*** | | |
| pACYC P$_{araB}$:tig | - | The chaperone plasmid kit was purchased from TaKaRa Bio Inc. (Cat. #3340). |
| pACYC P$_{araB}$::groES::groEL | - | |
| pACYC P$_{pzt-1}$::groES::groEL::tig | - | |
| pACYC P$_{araB}$::dnaK::dnaJ::grpE | - | |
| pACYC P$_{araB}$::dnaK::dnaJ::grpE-P$_{pzt-1}$::groES::groEL | - | |
| pACYC P$_{araB}$::groES::groEL (Amp$^R$) | Amp$^R$_F:<br>ATTAGTTAAGTATAAGAAGGAGATATACATATGATGAACGAAATCGACGAAAAAAATCAG<br>Amp$^R$_R:<br>TTTCGCAGCAGCGGTTTCTTTACCAGACTCGAGCGGACCCAGTAACAAAGCTTTATATAT<br>BB_groES/EL_F:<br>CTCGAGTCTGGTAAAGAAACC<br>BB_groES/EL_R:<br>CATATGTATATCTCCTTCTTATACTTAACTAATATACT | The chloramphenicol resistance marker of pACYC P$_{araB}$::groES::groEL was replaced with the ampicillin resistance marker from pETDuet-1 by Gibson Assembly method[6]. |

**Supplementary Table S2.** A list of enzymes used in this study.

| #Module | | Enzymes | Enzyme | Organism | Assigned function | #Accession (UniProt) |
|---|---|---|---|---|---|---|
| **1** | Butyryl-CoA synthesis | AtoB$_{Ec}$ | Acetyl-CoA acetyltransferase | *E. coli* | Synthesis of acetoacetyl-CoA from two acetyl-CoA | P76461 |
| | | Hbd$_{Ca}$ | 3-hydroxybutyryl-CoA dehydrogenase | *C. acetobutylicum* | Synthesis of 3-hydroxybutyryl-CoA from acetoacetyl-CoA | P52041 |
| | | Crt$_{Ca}$ | Short-chain-enoyl-CoA hydratase | *C. acetobutylicum* | Synthesis of crotonyl-CoA from 3-hydroxybutyryl-CoA | P52046 |
| | | Ter$_{Td}$ | Trans-2-enoyl-CoA reductase | *T. denticola* | Synthesis of butyryl-CoA from crotonyl-CoA | Q73Q47 |
| **2** | Alcohol synthesis | AdhE2$_{Ca}$ | Aldehyde-alcohol dehydrogenase | *C. acetobutylicum* | Synthesis of butanol from butyryl-CoA | Q9ANR5 |
| | | AdhE2$_{Ca}^{opt}$ | Codon optimized adhE2$_{Ca}$ for expression in *E. coli* | - | Improving soluble expression of adhE2$_{Ca}$ | - |
| | | TrxA_AdhE2$_{Ca}^{opt}$ | adhE2$_{Ca}^{opt}$ tagged with TrxA | - | | - |
| | | Pdc$_{Zm}$ | Pyruvate decarboxylase | *Z. mobilis* | Synthesis of acetaldehyde from pyruvate | P06672 |
| | | AdhB$_{Zm}$ | Alcohol dehydrogenase | *Z. mobilis* | Synthesis of ethanol from acetaldehyde | P0DJA2 |
| **3** | NADH synthesis | Fdh$_{Cb}^{opt}$ | Formate dehydrogenase | *C. boidinii* | Synthesis of NADH from NAD$^+$ | O13437 |
| **4** | Ester synthesis | ATF1$_{Sc}$ | Alcohol O-acetyltransferase | *S. cerevisiae* | Synthesis of an acetate ester from an alcohol and acetyl-CoA. | P40353 |
| | | ATF1$_{Sc}^{opt}$ | Codon optimized ATF1$_{Sc}$ for expression in *E. coli* | - | Improving soluble expression of ATF1$_{Sc}$ | - |
| | | MBP_ATF1$_{Sc}$ | ATF1$_{Sc}$ tagged with MBP without the N-terminus signal sequence | - | | - |
| | | NusA_ATF1$_{Sc}$ | ATF1$_{Sc}$ tagged with NusA | - | | - |
| | | TrxA_ATF1$_{Sc}$ | ATF1$_{Sc}$ tagged with TrxA | - | | - |
| | | TrxA_ATF1$_{Sc}^{opt}$ | ATF1$_{Sc}^{opt}$ tagged with TrxA | - | | - |
| | | SAAT$_{Fa}$ | Alcohol O-acetyltransferase | *F. ananassa* | Synthesis of a butyrate ester from an alcohol and butyryl-CoA. | Q9FVF1 |
| | | SAAT$_{Fa}^{opt}$ | Codon optimized SAAT$_{Fa}$ for expression in *E. coli* | - | Improving soluble expression of SAAT$_{Fa}$ | - |
| | | MBP_SAAT$_{Fa}$ | SAAT$_{Fa}$ tagged with MBP | - | | - |
| | | NusA_SAAT$_{Fa}$ | SAAT$_{Fa}$ tagged with NusA | - | | - |
| | | TrxA_SAAT$_{Fa}$ | SAAT$_{Fa}$ tagged with TrxA | - | | - |
| **etc.** | Fusion tags | MBP | Maltose binding protein without the N-terminus signal sequence (KIKTGARILALSALTTMMFSA SALA) | *E. coli* | Improving the solubility of heterologous proteins[1] | P0AEX9 |
| | | NusA | N-Utilization substrate A | *E. coli* | | P0AFF6 |
| | | TrxA | Thioredoxin 1 | *E. coli* | | P0AA25 |
| | Chaperones | Tf | Trigger factor | *E. coli* | Protein folding. Acts as a mechanical foldase by interacting directly with the newly synthesized proteins[2]. | P0A850 |
| | | DnaK | Chaperone protein | *E. coli* | Unfolding, disaggregation, stabilization of extended chains[3]. | P0A6Y8 |
| | | DnaJ | Chaperone protein | *E. coli* | | P08622 |
| | | GrpE | Protein GrpE | *E. coli* | | P09372 |
| | | GroES | 10 kDa chaperonin | *E. coli* | Protein folding, prevention of aggregation[3]. | P0A6F9 |
| | | GroEL | 60 kDa chaperonin | *E. coli* | | P0A6F5 |

**Supplementary Table S3.** A list of codon optimized gene sequences.

| Gene | Sequences |
|---|---|
| ATF1$_{Sc}^{opt}$ | ATGAACGAAATCGACGAAAAAAATCAGGCGCCAGTTCAGCAGGAGTGTCTGAAAGAAATGATTCAAAATGGTCATGCGCGTAGAATGGGTT<br>CAGTGGAGGATTTGTACGTTGCGCTGAATCGCCAAAACCTTTACCGTAACTTTTGCACGTATGGTGAGCTGTCAGATTACTGCACTCGTGATC<br>AGTTGACGCTGGCGCTCCGTGAAATCTGCTTAAAAAACCCGACTCTTCTGCATATTGTGTTACCGACGCGCTGGCCGAACCACGAAAACTAT<br>TATAGAAGCAGTGAATATTACAGCCGTCCTCACCCTGTGCACGACTACATATCGGTCTTACAGGAATTAAAGCTGTCGGGCGTGGTGTTGAA<br>TGAACAACCTGAATACTCAGCGGTTATGAAACAGATTCTGGAAGAATTCAAAAATAGCAAGGGCAGCTATACGGCTAAGATCTTTAAACTT<br>ACCACCACCCTGACAATCCCATACTTCGGCCCCACAGGCCCATCTTGGCGCTTGATTTGTTTGCCCGAAGAGCACACTGAAAAATGGAAGAA<br>ATTCATATTTGTGAGCAATCATTGCATGTCAGATGGCCGCTCCTCCATTCATTTCTTCCACGATCTGCGTGACGAACTTAACAATATTAAAAC<br>TCCGCCGAAGAAGCTAGATTACATCTTCAAATACGAAGAAGACTACCAGCTGCTGCGTAAACTGCCGGAGCCGATCGAAAAAGTGATTGAT<br>TTTCGTCCGCCGTATCTTTTTATTCCTAAATCCCTCTTATCGGGATTTATATACAATCACCTTCGATTTTCATCAAAAGGGGTATGTATGCGCA<br>TGGATGATGTGGAAAAGACGGACGACGTCGTTACCGAAATCATTAATATTAGCCCGACTGAATTTCAAGCGATAAAAGCAAACATCAAATC<br>GAACATCCAGGGCAAATGTACCATTACTCCCTTTCTCCATGTTTGTTGGTTCGTGTCTCTGCATAAATGGGGCAAGTTCTTTAAACCTCTGAA<br>CTTTGAATGGCTGACTGATATTTTCATCCCTGCGGACTGCCGCTCTCAGTTGCCTGATGACGATGAAATGCGCCAGATGTACCGGTATGGTGC<br>GAATGTGGGCTTCATTGACTTCACTCCGTGGATCAGCGAATTTGACATGAATGATAACAAAGAAAATTTTTGGCCACTGATTGAGCACTATC<br>ATGAGGTGATCAGTGAGGCGCTTCGTAACAAAAAACATCTGCATGGTTTGGGTTTCAACATTCAGGGTTTCGTACAAAAGTACGTGAACATA<br>GATAAAGTCATGTGTGACCGGGCTATTGGTAAACGTCGTGGCGGTACTCTACTCAGCAATGTTGGTCTGTTCAATCAACTTGAAGAACCTGA<br>CGCCAAATATTCTATTTGCGACTTAGCTTTTGGCCAATTCCAGGGCAGCTGGCACCAGGCTTTTAGTCTCGGTGTGTGCTCTACGAATGTTAA<br>AGGTATGAATATCGTCGTCGCGTCGACAAAAAACGTCGTAGGTTCACAGGAGTCCCTGGAAGAGCTTTGTAGCATATATAAAGCTTTGTTAC<br>TGGGTCCG |
| SAAT$_{Fa}^{opt}$ | ATGGAAAAAATCGAAGTCTCAATTAACTCCAAGCATACCATCAAACCCTCTACCTCTAGTACCCCGCTACAACCCTATAAACTGACCTTACT<br>GGATCAGCTGACGCCACCAGCCTATGTCCCGATCGTGTTTTTTTACCCGATCACCGATCACGATTTTAATCTGCCGCAGACGCTGGCGGACTT<br>ACGTCAAGCTCTGTCGGAGACTCTCACGCTGTATTATCCCCTGAGCGGACGTGTTAAAAACAATTTGTATATTGATGACTTTGAGGAAGGCG<br>TGCCGTACTTAGAAGCGCGTGTAAATTGCGATATGACCGATTTTCTGCGTCTGCGCAAAATTGAGTGCCTGAACGAATTTGTACCGATTAAA<br>CCTTTCAGTATGGAAGCAATTAGCGATGAACGTTACCCGTTGCTGGGCGTCCAGGTTAATGTTTTTGATTCGGGCATCGCGATCGGAGTGTC<br>AGTCAGTCATAAATTGATAGATGGTGGCACGGCAGATTGTTTCCTTAAATCTTGGGGCGCCGTGTTTCGTGGCTGCCGTGAAAATATTATCC<br>ACCCGAGTCTGAGTGAGGCTGCGCTGTTGTTCCCCCCACGCGATGACCTGCCAGAAAAATACGTTGACCAAATGGAGGCCCTGTGGTTTGCG<br>GGGAAAAAAGTCGCCACCCGACGCTTTGTTTTCGGTGTGAAAGCGATTAGCAGCATTCAAGACGAAGCAAAATCTGAGAGCGTCCCAAAAC<br>CGTCTCGCGTTCATGCCGTGACAGGCTTCTTATGGAAGCACCTGATAGCTGCGTCCCGCGCATTGACAAGTGGGACTACATCGACCCGTCTG<br>TCAATTGCCGCGCAAGCAGTAAACCTGCGTACTCGCATGAACATGGAAACCGTTTTAGATAACGCTACGGGTAACCTGTTCTGGTGGGCCCA<br>GGCGATCCTAGAACTGTCGCATACCACCCCGGAGATTTCTGACCTTAAACTGTGTGATCTGGTTAACTTGCTGAACGGATCCGTCAAGCAGT<br>GCAATGGCGACTATTTCGAGACCTTTAAAGGTAAAGAAGGCTATGGCCGGATGTGTGAATACTTGGATTTCCAGCGAACAATGTCGTCCATG<br>GAACCGGCTCCGGACATTTATCTGTTTTCATCCTGGACTAACTTTTTTAACCCGTTGGATTTCGGCTGGGGCCGCACCTCCTGGATTGGAGTT<br>GCAGGTAAAATTGAGAGTGCGTCCTGCAAATTTATAATCCTCGTGCCCACACAGTGCGGTTCGGGTATCGAGGCCTGGGTCAACTTGGAGGA<br>AGAAAAAATGGCCATGCTCGAACAGGATCCGCATTTTCTAGCGCTGGCGTCACCTAAAACCCTGATC |
| adhE2$_{Ca}^{opt}$ | ATGAAAGTGACCAATCAAAAAGAGCTGAAACAGAAACTGAACGAACTTCGTGAAGCGCAAAAAAAATTCGCAACCTACACGCAGGAGCAG<br>GTGGATAAAATTTTCAAGCAGTGTGCGATTGCGGCGGCGAAAGAGCGGATTAACCTGGCGAAGCTGGCGGTGGAAGAAACCGGGATTGGG<br>CTGGTGGAGGACAAAATCATCAAGAACCATTTTGCCGCGGAATACATTTATAACAAGTACAAAAACGAAAAGACGTGTGGTATTATTGATC<br>ACGATGACTCCCTGGGCATCACGAAGGTGGCAGAACCGATCGGTATCGTCGCTGCGATCGTTCCGACGACCAATCCCACGTCAACGGCGAT<br>CTTTAAAAGTTTGATTAGCCTGAAAACTCGTAACGCCATCTTCTTTTCCCCCCACCCCCGTGCGAAAAAGAGCACCATCGCGGCCGCGAAAC<br>TTATCTTGGATGCTGCCGTGAAGGCGGGGGCCCCGAAGAACATCATTGGCTGGATCGACGAGCCGTCAATTGAGCTCAGCCAAGACCTGAT<br>GAGCGAAGCGGATATTATTCTGGCGACAGGCGGTCCGAGCATGGTTAAAGCGGCCTACTCGTCTGGCAAACCAGCCATTGGTGTTGGAGCC<br>GGCAACACTCCGGCAATTATTGATGAGAGCGCCGATATCGATATGGCGGTTAGTAGCATCATCCTGTCTAAAACCTACGACAATGGTGTGAT<br>CTGCGCGAGCGAACAAAGTATTCTCGTCATGAACTCTATCTATGAAAAGGTGAAAGAAGAGTTTGTGAAGCGTGGTAGCTATATCCTGAACC<br>AGAACGAAATTGCCAAAATCAAAGAAACAATGTTTAAGAATGGAGCTATCAATGCCGATATCGTTGGTAAAAGTGCCTATATTATCGCGAA<br>AATGGCTGGCATCGAAGTCCCGCAAACTACCAAAATTTTAATTGGTGAGGTTCAGAGCGTGGAAAAATCGGAGTTGTTTTCCCACGAAAAA<br>CTGTCGCCGGTATTAGCAATGTATAAAGTAAAAGATTTTGACGAAGCCCTGAAAAAGGCCCAGCGCTTAATTGAATTAGGCGGCTCTGGTCA<br>TACCTCATCGCTGTACATCGATAGCCAGAACAATAAAGATAAAGTTAAAGAATTTGGCCTGGCCATGAAGACAAGTCGCACGTTTATCAATA<br>TGCCTTCAAGCCAGGGCGCCTCGGGTGATCTTTATAATTTTGCAATCGCGCCGAGCTTCACTCTGGGTTGTGGGACGTGGGGTGGCAATAGC<br>GTCTCACAGAATGTCGAACCGAAGCATTTACTGAACATCAAATCAGTCGCGGAACGCCGTGAAAACATGCTGTGGTTCAAAGTGCCGCAAA<br>AAATCTATTTCAAGTATGGTTGCCTGCGATTCGCACTGAAGGAACTGAAAGATATGAACAAAAAACGTGCGTTTATTGTTACCGATAAGGAC<br>CTGTTTAAACTGGGCTATGTGAACAAAATTACTAAAGTTCTGGATGAAATTGATATTAAATACTCGATCTTCACCGATATTAAAAGTGACCC<br>TACGATCGATAGCGTGAAAAAAGGCGCGAAAGAAATGCTTAACTTTGAACCGGATACCATTATCAGTATTGGGGGTGGTTCTCCGATGGAT<br>GCGGCCAAAGTTATGCATCTGTTGTATGAGTATCCGGAAGCCGAGATTGAGAACCTGGCTATCAATTTTATGGACATCCGAAAGCGGATCTG<br>CAACTTCCCAAAGTTAGGTACGAAAGCCATTAGCGTAGCCATTCCTACCACCGCGGGTACCGGTAGTGAGGCCACGCCGTTCGCGGTGATTA<br>CCAATGATGAGACCGGTATGAAATACCCGTTGACCAGCTATGAATTAACACCAAACATGGCAATTATCGATACGGAACTGATGCTGAACAT<br>GCCTCGTAAACTTACAGCGGCGACTGGGATTGATGCCCTGGTTCACGCGATTGAAGCATACGTGAGTGTGATGGCAACCGATTATACCGATG<br>AGCTGGCCCTGCGCGCTATCAAAATGATCTTCAAATATCTGCCGCGTGCCTACAAAAACGGCACCAATGATATCGAAGCTCGAGAAAAAAT<br>GGCGCATGCCTCAAACATCGCAGGGATGGCATTCGCGAACGCGTTTCTGGGCGTGTGCCACTCTATGGCACACAAACTGGGTGCGATGCACC<br>ATGTGCCACACGGCATTGCATGTGCTGTGTTAATTGAAGAAGTTATTAAATATAATGCGACCGACTGCCCAACCAAACAAACCGCGTTTCCG<br>CAATACAAAAGCCCGAATGCTAAGCGTAAATACGCTGAAATTGCGGAATATCTGAATCTGAAAGGCACCTCTGATACCGAGAAAGTGACCG<br>CCCTGATTGAAGCTATCTCTAAGTTAAAAATTGATCTGAGCATCCCGCAGAACATCTCGGCAGCTGGGATCAATAAAAAAGATTTTTACAAT<br>ACCTTGGATAAAATGTCTGAACTCGCGTTCGACGATCAATGCACAACCGCCAACCCACGCTATCCGCTGATCTCCGAACTCAAAGACATTTA<br>CATAAAGAGTTTC |
| fdh$_{Cb}^{opt}$ | ATGAAAATTGTGCTGGTTTTGTATGATGCGGGAAAGCATGCGGCGGACGAAGAAAAGCTTTATGGGTGTACCGAAAACAAACTGGGCATTG<br>CTAACTGGCTGAAAGACCAGGGTCATGAACTCATTACCACGAGCGATAAAGAAGGCGAAACCTCCGAATTAGACAAGCATATTCCGGATGC<br>GGACATTATTATCACCACTCCGTTTCACCCGGCATACATTACCAAAGAACGTTTAGATAAAGCTAAAAACCTGAAACTGGTCGTCGTCGCCG<br>GTGTCGGCAGTGATCACATTGATTTAGATTATATTAACCAGACGGGCAAAAAAATCTCCGTGTTAGAGGTAACAGGTAGCAACGTCGTCTCA<br>GTGGCGGAACACGTGGTAATGACCATGCTAGTTTTGGTCCGTAACTTTGTTCCGGCACATGAACAAATTATCAATCATGACTGGGAGGTCGC<br>GGCTATCGCGAAAGATGCCTACGATATTGAGGGCAAAACCATTGCGACTATTGGTGCGGGGCGCATCGGTTATCGTGTACTTGAACGCCTTC<br>TTCCGTTTAACCCGAAGGAGCTGTTGTACTACGACTACCAGGCGTTACCAAAAGAGGCTGAAGAAAAAGTGGGCGCGCGCCGTGTTGAGAA<br>TATTGAGGAACTCGTTGCACAGGCAGATATTGTTACCGTGAATGCGCCGCTGCACGCGGGAACCAAAGGTTTGATCAACAAAGAACTACTG<br>AGTAAATTTAAAAAAGGCGCCTGGCTCGTGAATACAGCTCGTGGTGCGATTTGCGTAGCTGAAGATGTGGCCGCAGCACTGGAGAGCGGTC<br>AGCTGCGCGGCTATGGGGGCGACGTGTGGTTCCCCCAACCGGCACCTAAAGATCATCCCTGGCGTGATATGCGTAACAAATACGGCGCAGG<br>CAATGCTATGACCCCGCATTACTCCGGCACAACCTTGGACGCACAAACTCGCTACGCGGAAGGTACTAAGAACATTCTGGAGTCGTTTTTA<br>CGGGCAAGTTTGATTACCGACCGCAGGATATAATTCTTCTGAACGGTGAATACGTGACCAAGGCGTATGGTAAACACGATAAAAAA |

**Supplementary Table S4.** Summary of validation of biosynthesis pathway of BA. Key strain is in bold; Abbreviations: *n.d.*: not detected. *n.a.*: not applicable.

| Strains | | EcJWBA1 | **EcJWBA2** | EcJWBA3 | EcJWBA4 | EcJWBA5 | EcJWBA6 |
|---|---|---|---|---|---|---|---|
| $O.D._{600}$ | | 2.13 ± 0.28 | 2.06 ± 0.05 | 2.34 ± 0.21 | 1.94 ± 0.02 | 1.20 ± 0.08 | 2.50 ± 0.13 |
| Consumed glucose, g/L (mM) | | 5.41 ± 0.43 (30.05 ± 2.37) | 4.20 ± 0.25 (23.31 ± 1.39) | 6.03 ± 0.13 (33.49 ± 0.72) | 5.26 ± 0.10 (29.20 ± 0.56) | 2.57 ± 0.16 (14.28 ± 0.91) | 6.63 ± 0.05 (36.83 ± 0.30) |
| Lactate, g/L (mM) | | 0.30 ± 0.07 (3.37 ± 0.75) | 0.18 ± 0.02 (2.00 ± 0.20) | 0.26 ± 0.06 (2.84 ± 0.62) | 0.18 ± 0.04 (2.03 ± 0.40) | 0.18 ± 0.01 (2.01 ± 0.08) | 0.23 ± 0.06 (2.51 ± 0.63) |
| Formate, g/L (mM) | | 0.51 ± 0.02 (11.10 ± 0.52) | 0.38 ± 0.01 (8.23 ± 0.20) | 0.50 ± 0.02 (10.77 ± 0.33) | 0.54 ± 0.02 (11.77 ± 0.36) | 0.49 ± 0.01 (10.61 ± 0.12) | 0.66 ± 0.04 (14.42 ± 0.94) |
| Acetate, g/L (mM) | | 0.62 ± 0.02 (10.37 ± 0.39) | 0.65 ± 0.00 (10.87 ± 0.02) | 0.54 ± 0.00 (9.02 ± 0.06) | 0.52 ± 0.00 (8.68 ± 0.03) | 0.40 ± 0.00 (6.74 ± 0.02) | 0.59 ± 0.02 (9.90 ± 0.32) |
| Ethanol, g/L (mM) | | 2.46 ± 0.12 (53.45 ± 2.70) | 2.41 ± 0.03 (52.38 ± 0.69) | 3.00 ± 0.08 (65.17 ± 1.82) | 2.54 ± 0.08 (55.21 ± 1.63) | 2.38 ± 0.03 (51.65 ± 0.68) | 2.46 ± 0.02 (53.34 ± 0.54) |
| Butanol, g/L (mM) | | 0.04 ± 0.01 (0.58 ± 0.12) | 0.06 ± 0.00 (0.84 ± 0.01) | 0.03 ± 0.00 (0.45 ± 0.01) | 0.03 ± 0.00 (0.42 ± 0.01) | 0.11 ± 0.01 (1.43 ± 0.13) | 0.05 ± 0.01 (0.67 ± 0.08) |
| Ester titers, mg/L (mM) | EA | 5.7 ± 1.3 (0.06 ± 0.01) | 11.9 ± 1.9 (0.13 ± 0.02) | 3.7 ± 0.5 (0.04 ± 0.01) | 0.8 ± 0.1 (0.01 ± 0.00) | 0.2 ± 0.0 (0.00 ± 0.00) | 1.3 ± 0.5 (0.01 ± 0.01) |
| | EB | *n.d.* | *n.d.* | *n.d.* | *n.d.* | *n.d.* | *n.d.* |
| | **BA** | 12.9 ± 5.9 (0.11 ± 0.05) | **34.2 ± 6.6 (0.29 ± 0.06)** | 3.8 ± 0.8 (0.03 ± 0.01) | 2.1 ± 0.3 (0.02 ± 0.00) | 0.7 ± 0.2 (0.01 ± 0.00) | 10.5 ± 4.6 (0.09 ± 0.04) |
| | BB | *n.d.* | *n.d.* | *n.d.* | *n.d.* | *n.d.* | *n.d.* |
| | Ester total | 18.6 ± 7.1 (0.18 ± 0.06) | 46.1 ± 8.5 (0.43 ± 0.08) | 7.5 ± 1.3 (0.07 ± 0.01) | 2.9 ± 0.4 (0.03 ± 0.00) | 0.9 ± 0.2 (0.01 ± 0.00) | 11.8 ± 5.0 (0.11 ± 0.04) |
| Specific ester productivity, mg/gDCW/h (μM/gDCW/h) | EA | 0.2 ± 0.0 (2.59 ± 0.27) | 0.5 ± 0.1 (5.62 ± 0.82) | 0.1 ± 0.0 (1.57 ± 0.33) | 0.0 ± 0.0 (0.40 ± 0.04) | 0.0 ± 0.0 (0.16 ± 0.01) | 0.1 ± 0.0 (0.51 ± 0.18) |
| | EB | *n.a.* | *n.a.* | *n.a.* | *n.a.* | *n.a.* | *n.a.* |
| | **BA** | 0.5 ± 0.2 (4.37 ± 1.36) | **1.4 ± 0.3 (12.31 ± 2.21)** | 0.1 ± 0.0 (1.22 ± 0.37) | 0.1 ± 0.0 (0.78 ± 0.12) | 0.1 ± 0.0 (0.42 ± 0.08) | 0.4 ± 0.2 (3.12 ± 1.33) |
| | BB | *n.a.* | *n.a.* | *n.a.* | *n.a.* | *n.a.* | *n.a.* |
| | Ester total | 0.7 ± 0.2 (6.95 ± 1.60) | 1.9 ± 0.3 (17.93 ± 3.04) | 0.3 ± 0.1 (2.79 ± 0.69) | 0.1 ± 0.0 (1.18 ± 0.16) | 0.1 ± 0.0 (0.58 ± 0.08) | 0.4 ± 0.2 (3.63 ± 1.50) |
| Ester yields, mg/g glucose (mM/M) | EA | 1.0 ± 0.2 (2.13 ± 0.32) | 2.8 ± 0.6 (5.81 ± 1.23) | 0.6 ± 0.1 (1.26 ± 0.18) | 0.2 ± 0.0 (0.31 ± 0.04) | 0.1 ± 0.0 (0.16 ± 0.03) | 0.2 ± 0.1 (0.40 ± 0.14) |
| | EB | *n.a.* | *n.a.* | *n.a.* | *n.a.* | *n.a.* | *n.a.* |
| | **BA** | 2.3 ± 0.9 (3.62 ± 1.33) | **8.2 ± 2.0 (12.75 ± 3.13)** | 0.6 ± 0.2 (0.98 ± 0.24) | 0.4 ± 0.1 (0.60 ± 0.11) | 0.3 ± 0.1 (0.42 ± 0.13) | 1.6 ± 0.7 (2.46 ± 1.08) |
| | BB | *n.a.* | *n.a.* | *n.a.* | *n.a.* | *n.a.* | *n.a.* |
| | Ester total | 3.4 ± 1.0 (5.75 ± 1.63) | 11.1 ± 2.6 (18.56 ± 4.36) | 1.3 ± 0.2 (2.24 ± 0.41) | 0.5 ± 0.1 (0.92 ± 0.15) | 0.4 ± 0.1 (0.58 ± 0.15) | 1.8 ± 0.8 (2.86 ± 1.23) |
| Culture time (h) | | 24 | 24 | 24 | 24 | 24 | 24 |

**Supplementary Table S5.** Summary of validation of biosynthesis pathway of EB. Key strain is in bold; Abbreviations: *n.d.*: not detected. *n.a.*: not applicable.

| Strains | | EcJWEB1 | **EcJWEB2** | EcJWEB3 | EcJWEB4 | EcJWEB5 | EcJWEB6 |
|---|---|---|---|---|---|---|---|
| $O.D._{600}$ | | 2.12 ± 0.03 | 3.39 ± 0.26 | 3.86 ± 0.24 | 3.02 ± 0.08 | 0.93 ± 0.02 | 2.79 ± 0.06 |
| Consumed glucose, g/L (mM) | | 7.53 ± 0.13 (41.79 ± 0.72) | 7.21 ± 0.39 (40.02 ± 2.19) | 19.61 ± 0.20 (108.86 ± 1.10) | 13.40 ± 0.22 (74.40 ± 1.24) | 2.77 ± 0.37 (15.37 ± 2.05) | 9.30 ± 0.39 (51.62 ± 2.16) |
| Lactate, g/L (mM) | | 0.02 ± 0.03 (0.22 ± 0.34) | *n.d.* | 0.19 ± 0.01 (2.10 ± 0.16) | 0.18 ± 0.00 (1.95 ± 0.05) | *n.d.* | 0.06 ± 0.00 (0.62 ± 0.01) |
| Formate, g/L (mM) | | 0.44 ± 0.01 (9.63 ± 0.16) | 0.17 ± 0.01 (3.79 ± 0.12) | 0.61 ± 0.03 (13.32 ± 0.57) | 0.69 ± 0.02 (14.96 ± 0.39) | 0.18 ± 0.02 (3.93 ± 0.46) | 0.56 ± 0.01 (12.23 ± 0.32) |
| Acetate, g/L (mM) | | 0.48 ± 0.01 (7.94 ± 0.12) | 0.37 ± 0.00 (6.10 ± 0.04) | 0.56 ± 0.01 (9.25 ± 0.09) | 0.53 ± 0.00 (8.80 ± 0.07) | 0.36 ± 0.00 (6.03 ± 0.06) | 0.49 ± 0.01 (8.16 ± 0.11) |
| Ethanol, g/L (mM) | | 4.92 ± 0.11 (106.72 ± 2.32) | 5.16 ± 0.14 (111.92 ± 2.96) | 10.89 ± 0.10 (236.41 ± 2.13) | 7.25 ± 0.04 (157.34 ± 0.90) | 2.84 ± 0.04 (61.66 ± 0.89) | 5.29 ± 0.15 (114.85 ± 3.27) |
| Butanol, g/L (mM) | | *n.d.* | *n.d.* | 0.02 ± 0.00 (0.25 ± 0.02) | *n.d.* | 0.03 ± 0.00 (0.40 ± 0.02) | *n.d.* |
| Ester titers, mg/L (mM) | EA | 0.5 ± 0.2 (0.01 ± 0.00) | 4.9 ± 0.5 (0.06 ± 0.01) | 7.4 ± 1.7 (0.08 ± 0.02) | 2.5 ± 0.4 (0.03 ± 0.00) | 0.1 ± 0.1 (0.00 ± 0.00) | 1.8 ± 0.3 (0.02 ± 0.00) |
| | **EB** | 17.1 ± 3.4 (0.15 ± 0.03) | **71.0 ± 6.6 (0.61 ± 0.06)** | 21.7 ± 2.9 (0.19 ± 0.03) | 8.4 ± 1.0 (0.07 ± 0.01) | 1.4 ± 0.2 (0.01 ± 0.00) | 15.8 ± 1.3 (0.14 ± 0.01) |
| | BA | *n.d.* | *n.d.* | *n.d.* | *n.d.* | *n.d.* | *n.d.* |
| | BB | *n.d.* | 1.3 ± 0.7 (0.01 ± 0.00) | *n.d.* | *n.d.* | *n.d.* | *n.d.* |
| | Ester total | 17.6 ± 3.2 (0.15 ± 0.03) | 77.2 ± 7.7 (0.68 ± 0.07) | 29.1 ± 4.7 (0.27 ± 0.04) | 10.9 ± 1.3 (0.10 ± 0.01) | 1.5 ± 0.1 (0.01 ± 0.00) | 17.6 ± 1.6 (0.16 ± 0.01) |
| Specific ester productivity, mg/gDCW/h (µM/gDCW/h) | EA | 0.0 ± 0.0 (0.22 ± 0.10) | 0.1 ± 0.0 (1.42 ± 0.17) | 0.2 ± 0.1 (1.88 ± 0.57) | 0.1 ± 0.0 (0.82 ± 0.14) | 0.0 ± 0.0 (0.07 ± 0.07) | 0.1 ± 0.0 (0.61 ± 0.11) |
| | **EB** | 0.7 ± 0.1 (5.98 ± 1.10) | **1.8 ± 0.2 (15.60 ± 1.95)** | 0.5 ± 0.1 (4.20 ± 0.85) | 0.2 ± 0.0 (2.05 ± 0.20) | 0.1 ± 0.0 (1.09 ± 0.12) | 0.5 ± 0.1 (4.20 ± 0.43) |
| | BA | *n.a.* | *n.a.* | *n.a.* | *n.a.* | *n.a.* | *n.a.* |
| | BB | *n.a.* | 0.0 ± 0.0 (0.23 ± 0.12) | *n.a.* | *n.a.* | *n.a.* | *n.a.* |
| | Ester total | 0.7 ± 0.1 (6.19 ± 1.00) | 2.0 ± 0.3 (17.25 ± 2.24) | 0.7 ± 0.2 (6.08 ± 1.41) | 0.3 ± 0.0 (2.87 ± 0.31) | 0.1 ± 0.0 (1.16 ± 0.05) | 0.5 ± 0.1 (4.81 ± 0.53) |
| Ester yields, mg/g glucose (mM/M) | EA | 0.1 ± 0.0 (0.13 ± 0.06) | 0.7 ± 0.0 (1.39 ± 0.06) | 0.4 ± 0.1 (0.77 ± 0.19) | 0.2 ± 0.0 (0.39 ± 0.07) | 0.0 ± 0.0 (0.05 ± 0.04) | 0.2 ± 0.0 (0.39 ± 0.07) |
| | **EB** | 2.3 ± 0.4 (3.53 ± 0.66) | **9.8 ± 0.4 (15.26 ± 0.65)** | 1.1 ± 0.2 (1.72 ± 0.25) | 0.6 ± 0.1 (0.97 ± 0.12) | 0.5 ± 0.1 (0.79 ± 0.21) | 1.7 ± 0.2 (2.65 ± 0.33) |
| | BA | *n.a.* | *n.a.* | *n.a.* | *n.a.* | *n.a.* | *n.a.* |
| | BB | *n.a.* | 0.2 ± 0.1 (0.22 ± 0.10) | *n.a.* | *n.a.* | *n.a.* | *n.a.* |
| | Ester total | 2.3 ± 0.4 (3.65 ± 0.60) | 10.7 ± 0.5 (16.87 ± 0.81) | 1.5 ± 0.3 (2.49 ± 0.44) | 0.8 ± 0.1 (1.35 ± 0.18) | 0.5 ± 0.1 (0.84 ± 0.17) | 1.9 ± 0.2 (3.03 ± 0.40) |
| Culture time (h) | | 24 | 24 | 24 | 24 | 24 | 24 |

**Supplementary Table S6.** Summary of validation of biosynthesis pathway of BB. Key strain is in bold; Abbreviations: *n.d.*: not detected. *n.a.*: not applicable.

| Strains | | EcJWBB1 | **EcJWBB2** | EcJWBB3 | EcJWBB4 | EcJWBB5 | EcJWBB6 |
|---|---|---|---|---|---|---|---|
| O.D.$_{600}$ | | 2.31 ± 0.10 | 1.96 ± 0.02 | 2.61 ± 0.08 | 2.31 ± 0.06 | 1.24 ± 0.05 | 2.01 ± 0.07 |
| Consumed glucose, g/L (mM) | | 6.98 ± 0.17 (38.72 ± 0.96) | 3.90 ± 0.26 (21.63 ± 1.43) | 10.46 ± 0.15 (58.04 ± 0.86) | 8.90 ± 1.05 (49.42 ± 5.82) | 1.75 ± 0.04 (9.71 ± 0.20) | 5.15 ± 0.21 (28.60 ± 1.18) |
| Lactate, g/L (mM) | | 0.21 ± 0.01 (2.32 ± 0.14) | 0.09 ± 0.01 (1.05 ± 0.06) | 0.17 ± 0.01 (1.87 ± 0.15) | 0.16 ± 0.02 (1.76 ± 0.17) | 0.10 ± 0.01 (1.15 ± 0.13) | 0.17 ± 0.01 (1.87 ± 0.11) |
| Formate, g/L (mM) | | 0.49 ± 0.01 (10.56 ± 0.14) | 0.36 ± 0.01 (7.82 ± 0.17) | 0.56 ± 0.02 (12.25 ± 0.50) | 0.63 ± 0.02 (13.70 ± 0.40) | 0.40 ± 0.05 (8.67 ± 1.19) | 0.61 ± 0.01 (13.27 ± 0.15) |
| Acetate, g/L (mM) | | 0.54 ± 0.00 (8.93 ± 0.03) | 0.46 ± 0.00 (7.71 ± 0.05) | 0.53 ± 0.01 (8.87 ± 0.10) | 0.52 ± 0.02 (8.59 ± 0.32) | 0.40 ± 0.01 (6.64 ± 0.09) | 0.49 ± 0.00 (8.15 ± 0.04) |
| Ethanol, g/L (mM) | | 3.04 ± 0.07 (65.93 ± 1.53) | 2.66 ± 0.09 (57.82 ± 1.88) | 3.68 ± 0.08 (79.96 ± 1.74) | 2.91 ± 0.16 (63.23 ± 3.44) | 2.47 ± 0.30 (53.58 ± 6.42) | 2.50 ± 0.03 (54.21 ± 0.71) |
| Butanol, g/L (mM) | | 0.11 ± 0.01 (1.54 ± 0.15) | 0.13 ± 0.00 (1.71 ± 0.06) | 0.02 ± 0.00 (0.24 ± 0.00) | 0.04 ± 0.03 (0.54 ± 0.43) | 0.09 ± 0.02 (1.27 ± 0.29) | 0.13 ± 0.01 (1.76 ± 0.12) |
| Ester titers, mg/L (mM) | EA | 0.7 ± 0.1 (0.01 ± 0.00) | 1.5 ± 0.3 (0.02 ± 0.00) | 4.6 ± 0.5 (0.05 ± 0.01) | 1.1 ± 0.2 (0.01 ± 0.00) | *n.d.* | 0.4 ± 0.6 (0.00 ± 0.01) |
| | EB | 43.6 ± 6.1 (0.38 ± 0.05) | 126.2 ± 11.5 (1.09 ± 0.10) | 5.3 ± 0.1 (0.05 ± 0.00) | 1.6 ± 0.2 (0.01 ± 0.00) | 0.4 ± 0.7 (0.00 ± 0.01) | 4.6 ± 3.1 (0.04 ± 0.03) |
| | BA | 0.3 ± 0.1 (0.00 ± 0.00) | 0.3 ± 0.0 (0.00 ± 0.00) | 0.2 ± 0.1 (0.00 ± 0.00) | 0.2 ± 0.1 (0.00 ± 0.00) | *n.d.* | 0.0 ± 0.1 (0.00 ± 0.00) |
| | **BB** | 16.0 ± 3.8 (0.11 ± 0.03) | **33.5 ± 2.9 (0.23 ± 0.02)** | *n.d.* | 1.0 ± 0.4 (0.01 ± 0.00) | *n.d.* | 11.5 ± 1.0 (0.08 ± 0.01) |
| | Ester total | 60.5 ± 9.9 (0.50 ± 0.08) | 161.4 ± 14.4 (1.34 ± 0.12) | 10.0 ± 0.6 (0.10 ± 0.01) | 4.0 ± 0.4 (0.04 ± 0.00) | 0.4 ± 0.7 (0.00 ± 0.01) | 16.5 ± 1.7 (0.12 ± 0.01) |
| Specific ester productivity, mg/gDCW/h (µM/gDCW/h) | EA | 0.0 ± 0.0 (0.29 ± 0.03) | 0.1 ± 0.0 (0.75 ± 0.14) | 0.2 ± 0.0 (1.71 ± 0.24) | 0.0 ± 0.0 (0.48 ± 0.09) | *n.a.* | 0.0 ± 0.0 (0.19 ± 0.32) |
| | EB | 1.6 ± 0.2 (13.95 ± 1.71) | 5.5 ± 0.5 (47.66 ± 4.16) | 0.2 ± 0.0 (1.50 ± 0.07) | 0.1 ± 0.0 (0.52 ± 0.05) | 0.0 ± 0.1 (0.24 ± 0.42) | 0.2 ± 0.1 (1.68 ± 1.10) |
| | BA | 0.0 ± 0.0 (0.08 ± 0.03) | 0.0 ± 0.0 (0.11 ± 0.01) | 0.0 ± 0.0 (0.05 ± 0.02) | 0.0 ± 0.0 (0.07 ± 0.03) | *n.a.* | 0.0 ± 0.0 (0.02 ± 0.03) |
| | **BB** | 0.6 ± 0.1 (4.11 ± 0.86) | **1.5 ± 0.1 (10.19 ± 0.89)** | *n.a.* | 0.0 ± 0.0 (0.26 ± 0.09) | *n.a.* | 0.5 ± 0.1 (3.42 ± 0.37) |
| | Ester total | 2.3 ± 0.3 (18.43 ± 2.57) | 7.1 ± 0.6 (58.71 ± 5.10) | 0.3 ± 0.0 (3.26 ± 0.32) | 0.2 ± 0.0 (1.33 ± 0.07) | 0.0 ± 0.1 (0.24 ± 0.42) | 0.7 ± 0.1 (5.30 ± 0.44) |
| Ester yields, mg/g glucose (mM/M) | EA | 0.1 ± 0.0 (0.20 ± 0.01) | 0.4 ± 0.1 (0.80 ± 0.21) | 0.4 ± 0.1 (0.89 ± 0.10) | 0.1 ± 0.0 (0.26 ± 0.03) | *n.a.* | 0.1 ± 0.1 (0.14 ± 0.25) |
| | EB | 6.2 ± 0.8 (9.68 ± 1.16) | 32.6 ± 5.2 (50.57 ± 8.06) | 0.5 ± 0.0 (0.78 ± 0.02) | 0.2 ± 0.0 (0.29 ± 0.06) | 0.2 ± 0.4 (0.36 ± 0.62) | 0.9 ± 0.6 (1.40 ± 0.95) |
| | BA | 0.0 ± 0.0 (0.06 ± 0.02) | 0.1 ± 0.0 (0.11 ± 0.02) | 0.0 ± 0.0 (0.03 ± 0.01) | 0.0 ± 0.0 (0.04 ± 0.01) | *n.a.* | 0.0 ± 0.0 (0.01 ± 0.02) |
| | **BB** | 2.3 ± 0.5 (2.86 ± 0.63) | **8.7 ± 1.3 (10.81 ± 1.65)** | *n.a.* | 0.1 ± 0.1 (0.15 ± 0.07) | *n.a.* | 2.2 ± 0.2 (2.79 ± 0.28) |
| | Ester total | 8.7 ± 1.3 (12.80 ± 1.80) | 41.7 ± 6.6 (62.29 ± 9.88) | 1.0 ± 0.1 (1.71 ± 0.13) | 0.5 ± 0.1 (0.74 ± 0.14) | 0.2 ± 0.4 (0.36 ± 0.62) | 3.2 ± 0.4 (4.34 ± 0.64) |
| Culture time (h) | | 24 | 24 | 24 | 24 | 24 | 24 |

**Supplementary Table S7.** Summary of induction conditions optimization for biosynthesis pathway of BA. The best culture condition is in bold; Abbreviations: *n.d.*: not detected. *n.a.*: not applicable.

| Strain | | EcJWBA2 | | | | | |
|---|---|---|---|---|---|---|---|
| Temperature | | **28'C** | | | 37'C | | |
| IPTG concentration, mM | | 0.01 | **0.1** | 1.0 | 0.01 | 0.1 | 1.0 |
| O.D.$_{600}$ | | 6.55 ± 0.12 | 3.80 ± 0.02 | 1.62 ± 0.08 | 8.95 ± 0.23 | 2.50 ± 0.07 | 1.78 ± 0.06 |
| Consumed glucose, g/L (mM) | | 7.77 ± 0.16 (43.14 ± 0.90) | 4.39 ± 0.19 (24.35 ± 1.03) | 2.02 ± 0.02 (11.23 ± 0.11) | 12.17 ± 0.40 (67.53 ± 2.23) | 3.82 ± 0.36 (21.21 ± 2.00) | 3.54 ± 0.51 (19.63 ± 2.84) |
| Lactate, g/L (mM) | | 0.26 ± 0.01 (2.88 ± 0.15) | 0.27 ± 0.06 (2.95 ± 0.64) | 0.01 ± 0.02 (0.15 ± 0.18) | 0.84 ± 0.02 (9.38 ± 0.17) | 0.11 ± 0.00 (1.22 ± 0.02) | 0.08 ± 0.01 (0.91 ± 0.10) |
| Formate, g/L (mM) | | 2.32 ± 0.06 (50.34 ± 1.27) | 0.86 ± 0.35 (18.68 ± 7.50) | 0.23 ± 0.02 (4.95 ± 0.37) | 2.35 ± 0.07 (51.15 ± 1.42) | 0.51 ± 0.01 (11.17 ± 0.17) | 0.61 ± 0.02 (13.18 ± 0.49) |
| Acetate, g/L (mM) | | 0.94 ± 0.02 (15.62 ± 0.26) | 0.76 ± 0.01 (12.68 ± 0.09) | 0.41 ± 0.00 (6.80 ± 0.07) | 0.87 ± 0.01 (14.42 ± 0.16) | 0.55 ± 0.00 (9.15 ± 0.04) | 0.43 ± 0.01 (7.09 ± 0.17) |
| Ethanol, g/L (mM) | | 3.56 ± 0.11 (77.24 ± 2.28) | 2.30 ± 0.23 (49.97 ± 5.07) | 1.69 ± 0.02 (36.62 ± 0.44) | 3.06 ± 0.08 (66.42 ± 1.65) | 2.39 ± 0.04 (51.82 ± 0.79) | 2.69 ± 0.17 (58.31 ± 3.71) |
| Butanol, g/L (mM) | | 0.17 ± 0.03 (2.33 ± 0.43) | 0.09 ± 0.04 (1.19 ± 0.58) | 0.06 ± 0.01 (0.86 ± 0.07) | 0.07 ± 0.00 (0.88 ± 0.05) | 0.10 ± 0.01 (1.32 ± 0.07) | 0.11 ± 0.00 (1.45 ± 0.06) |
| Ester titers, mg/L (mM) | EA | 10.1 ± 2.4 (0.11 ± 0.03) | 8.6 ± 2.6 (0.10 ± 0.03) | 2.4 ± 0.9 (0.03 ± 0.01) | 13.4 ± 5.1 (0.15 ± 0.06) | 5.3 ± 1.1 (0.06 ± 0.01) | 2.1 ± 0.8 (0.02 ± 0.01) |
| | EB | *n.d.* | *n.d.* | *n.d.* | *n.d.* | *n.d.* | *n.d.* |
| | BA | 35.9 ± 5.5 (0.31 ± 0.05) | **48.0 ± 7.1 (0.41 ± 0.06)** | 8.0 ± 3.8 (0.07 ± 0.03) | 19.5 ± 7.3 (0.17 ± 0.06) | 15.0 ± 1.8 (0.13 ± 0.02) | 8.9 ± 1.5 (0.08 ± 0.01) |
| | BB | *n.d.* | *n.d.* | *n.d.* | *n.d.* | *n.d.* | *n.d.* |
| | Ester total | 46.0 ± 5.6 (0.42 ± 0.05) | 56.6 ± 9.7 (0.51 ± 0.09) | 10.3 ± 4.7 (0.10 ± 0.04) | 32.8 ± 12.4 (0.32 ± 0.12) | 20.3 ± 2.4 (0.19 ± 0.02) | 11.0 ± 2.2 (0.10 ± 0.02) |
| Specific ester productivity, mg/gDCW/h (µM/gDCW/h) | EA | 0.1 ± 0.0 (1.51 ± 0.33) | 0.2 ± 0.1 (2.21 ± 0.69) | 0.1 ± 0.1 (1.43 ± 0.58) | 0.1 ± 0.1 (1.47 ± 0.59) | 0.2 ± 0.0 (2.06 ± 0.39) | 0.1 ± 0.0 (1.15 ± 0.37) |
| | EB | *n.a.* | *n.a.* | *n.a.* | *n.a.* | *n.a.* | *n.a.* |
| | BA | 0.5 ± 0.1 (4.06 ± 0.61) | **1.1 ± 0.2 (9.35 ± 1.46)** | 0.4 ± 0.2 (3.68 ± 1.80) | 0.2 ± 0.1 (1.62 ± 0.64) | 0.5 ± 0.1 (4.44 ± 0.41) | 0.4 ± 0.1 (3.70 ± 0.54) |
| | BB | *n.a.* | *n.a.* | *n.a.* | *n.a.* | *n.a.* | *n.a.* |
| | Ester total | 0.6 ± 0.1 (5.57 ± 0.59) | 1.3 ± 0.2 (11.57 ± 2.14) | 0.6 ± 0.3 (5.11 ± 2.37) | 0.3 ± 0.1 (3.09 ± 1.23) | 0.7 ± 0.1 (6.50 ± 0.63) | 0.5 ± 0.1 (4.85 ± 0.85) |
| Ester yields, mg/g glucose (mM/M) | EA | 1.3 ± 0.3 (2.67 ± 0.67) | 1.9 ± 0.5 (3.97 ± 0.98) | 1.1 ± 0.6 (2.29 ± 1.26) | 1.1 ± 0.4 (2.24 ± 0.80) | 1.4 ± 0.2 (2.81 ± 0.37) | 0.6 ± 0.1 (1.20 ± 0.25) |
| | EB | *n.a.* | *n.a.* | *n.a.* | *n.a.* | *n.a.* | *n.a.* |
| | BA | 4.6 ± 0.6 (7.16 ± 0.97) | **10.9 ± 1.0 (16.88 ± 1.51)** | 4.0 ± 2.6 (6.23 ± 4.07) | 1.6 ± 0.6 (2.47 ± 0.88) | 4.0 ± 0.6 (6.13 ± 0.99) | 2.5 ± 0.3 (3.91 ± 0.42) |
| | BB | *n.a.* | *n.a.* | *n.a.* | *n.a.* | *n.a.* | *n.a.* |
| | Ester total | 5.9 ± 0.7 (9.83 ± 1.08) | 9.0 ± 0.6 (14.72 ± 0.92) | 5.1 ± 3.2 (8.52 ± 5.32) | 2.7 ± 1.0 (4.73 ± 1.66) | 5.3 ± 0.7 (8.94 ± 1.08) | 3.1 ± 0.3 (5.10 ± 0.47) |
| Culture time (h) | | 24 | 24 | 24 | 24 | 24 | 24 |

**Supplementary Table S8.** Summary of induction conditions optimization for biosynthesis pathway of EB. The best culture condition is in bold; Abbreviations: *n.d.*: not detected. *n.a.*: not applicable.

| Strain | | EcJWEB2 | | | | | |
|---|---|---|---|---|---|---|---|
| Temperature | | **28’C** | | | 37’C | | |
| IPTG concentration, mM | | 0.01 | **0.1** | 1.0 | 0.01 | 0.1 | 1.0 |
| O.D.$_{600}$ | | 5.90 ± 0.21 | 3.88 ± 0.10 | 2.07 ± 0.14 | 17.37 ± 0.74 | 3.03 ± 0.03 | 3.50 ± 0.51 |
| Consumed glucose, g/L (mM) | | 18.69 ± 0.26 (103.73 ± 1.46) | 9.84 ± 0.24 (54.63 ± 1.35) | 3.32 ± 0.46 (18.44 ± 2.54) | 40.80 ± 2.00 (226.47 ± 11.10) | 8.14 ± 0.23 (45.19 ± 1.26) | 7.71 ± 1.38 (42.82 ± 7.64) |
| Lactate, g/L (mM) | | 0.02 ± 0.00 (0.17 ± 0.00) | *n.d.* | 0.03 ± 0.00 (0.29 ± 0.00) | 1.37 ± 0.08 (15.17 ± 0.86) | 0.03 ± 0.00 (0.39 ± 0.02) | *n.d.* |
| Formate, g/L (mM) | | 0.11 ± 0.01 (2.44 ± 0.16) | 0.12 ± 0.00 (2.60 ± 0.11) | 0.04 ± 0.00 (0.80 ± 0.06) | 0.42 ± 0.15 (9.23 ± 3.28) | 0.38 ± 0.01 (8.33 ± 0.25) | 0.31 ± 0.10 (6.77 ± 2.09) |
| Acetate, g/L (mM) | | 0.37 ± 0.00 (6.16 ± 0.03) | 0.27 ± 0.00 (4.48 ± 0.03) | 0.25 ± 0.00 (4.09 ± 0.04) | 0.63 ± 0.00 (10.44 ± 0.07) | 0.28 ± 0.00 (4.66 ± 0.00) | 0.26 ± 0.02 (4.40 ± 0.26) |
| Ethanol, g/L (mM) | | 9.52 ± 0.21 (206.66 ± 4.61) | 5.64 ± 0.09 (122.33 ± 1.88) | 2.35 ± 0.19 (50.91 ± 4.10) | 18.55 ± 0.30 (402.67 ± 6.44) | 5.14 ± 0.08 (111.56 ± 1.67) | 5.18 ± 0.39 (112.35 ± 8.41) |
| Butanol, g/L (mM) | | 0.05 ± 0.02 (0.69 ± 0.22) | *n.d.* | *n.d.* | 0.23 ± 0.00 (3.11 ± 0.02) | *n.d.* | *n.d.* |
| Ester titers, mg/L (mM) | EA | 14.5 ± 4.7 (0.17 ± 0.05) | 18.2 ± 1.6 (0.21 ± 0.02) | 2.3 ± 1.1 (0.03 ± 0.01) | 8.29 ± 0.95 (0.09 ± 0.01) | 4.50 ± 0.29 (0.05 ± 0.00) | 5.75 ± 1.59 (0.07 ± 0.02) |
| | EB | 123.4 ± 37.2 (1.06 ± 0.32) | **200.4 ± 9.4 (1.73 ± 0.08)** | 23.9 ± 20.0 (0.21 ± 0.17) | 131.46 ± 12.45 (1.13 ± 0.11) | 113.41 ± 34.13 (0.98 ± 0.29) | 48.29 ± 19.57 (0.42 ± 0.17) |
| | BA | 0.1 ± 0.2 (0.00 ± 0.00) | *n.d.* | *n.d.* | 0.45 ± 0.06 (0.00 ± 0.00) | *n.d.* | *n.d.* |
| | BB | 6.7 ± 1.7 (0.05 ± 0.01) | 5.1 ± 0.3 (0.04 ± 0.00) | *n.d.* | 14.25 ± 2.47 (0.10 ± 0.02) | 1.67 ± 1.47 (0.01 ± 0.01) | 0.94 ± 1.33 (0.01 ± 0.01) |
| | Ester total | 144.8 ± 42.7 (1.27 ± 0.38) | 223.7 ± 10.7 (1.97 ± 0.10) | 26.2 ± 21.1 (0.23 ± 0.18) | 154.46 ± 15.93 (1.33 ± 0.14) | 119.58 ± 32.68 (1.04 ± 0.28) | 54.99 ± 19.31 (0.49 ± 0.16) |
| Specific ester productivity, mg/gDCW/h (μM/gDCW/h) | EA | 0.2 ± 0.1 (2.43 ± 0.85) | 0.4 ± 0.0 (4.58 ± 0.25) | 0.1 ± 0.1 (1.10 ± 0.57) | 0.04 ± 0.00 (0.46 ± 0.03) | 0.13 ± 0.01 (1.45 ± 0.08) | 0.09 ± 0.09 (1.07 ± 1.06) |
| | EB | 1.8 ± 0.6 (15.58 ± 5.06) | **4.4 ± 0.1 (38.19 ± 0.48)** | 1.0 ± 0.9 (8.68 ± 7.42) | 0.64 ± 0.03 (5.51 ± 0.28) | 3.23 ± 1.00 (27.80 ± 8.63) | 0.75 ± 0.67 (6.43 ± 5.80) |
| | BA | 0.0 ± 0.0 (0.01 ± 0.02) | *n.a.* | *n.a.* | 0.00 ± 0.00 (0.02 ± 0.00) | *n.a.* | *n.a.* |
| | BB | 0.1 ± 0.0 (0.68 ± 0.18) | 0.1 ± 0.0 (0.78 ± 0.08) | *n.a.* | 0.07 ± 0.01 (0.48 ± 0.06) | 0.05 ± 0.04 (0.33 ± 0.29) | 0.01 ± 0.02 (0.09 ± 0.16) |
| | Ester total | 2.1 ± 0.7 (18.70 ± 6.00) | 5.0 ± 0.1 (43.55 ± 0.65) | 1.1 ± 0.9 (9.78 ± 7.98) | 0.75 ± 0.04 (6.47 ± 0.38) | 3.40 ± 0.96 (29.57 ± 8.31) | 0.86 ± 0.76 (7.60 ± 6.69) |
| Ester yields, mg/g glucose (mM/M) | EA | 0.8 ± 0.2 (1.59 ± 0.49) | 1.9 ± 0.1 (3.83 ± 0.27) | 0.7 ± 0.3 (1.41 ± 0.65) | 0.14 ± 0.12 (0.28 ± 0.24) | 0.55 ± 0.05 (1.13 ± 0.10) | 0.69 ± 0.07 (1.41 ± 0.15) |
| | EB | 6.6 ± 2.0 (10.25 ± 3.10) | **20.6 ± 0.6 (31.96 ± 0.91)** | 7.1 ± 5.7 (11.03 ± 8.87) | 2.17 ± 1.88 (3.37 ± 2.92) | 13.88 ± 3.92 (21.52 ± 6.08) | 6.16 ± 3.45 (9.55 ± 5.35) |
| | BA | 0.0 ± 0.0 (0.01 ± 0.01) | *n.a.* | *n.a.* | 0.01 ± 0.01 (0.01 ± 0.01) | *n.a.* | *n.a.* |
| | BB | 0.4 ± 0.1 (0.45 ± 0.11) | 0.5 ± 0.0 (0.66 ± 0.06) | *n.a.* | 0.24 ± 0.21 (0.29 ± 0.26) | 0.21 ± 0.18 (0.26 ± 0.23) | 0.13 ± 0.18 (0.16 ± 0.23) |
| | Ester total | 7.8 ± 2.3 (12.30 ± 3.63) | 23.0 ± 0.7 (36.45 ± 1.12) | 7.8 ± 6.0 (12.44 ± 9.52) | 2.55 ± 2.21 (3.96 ± 3.43) | 14.63 ± 3.72 (22.91 ± 5.79) | 6.98 ± 3.57 (11.13 ± 5.44) |
| Culture time (h) | | 24 | 24 | 24 | 24 | 24 | 24 |

**Supplementary Table S9.** Summary of induction conditions optimization for biosynthesis pathway of BB. The best culture condition is in bold; Abbreviations: *n.d.*: not detected. *n.a.*: not applicable.

| Strain | | EcJWBB2 | | | | | |
|---|---|---|---|---|---|---|---|
| Temperature | | 28'C | | | **37'C** | | |
| IPTG concentration, mM | | 0.01 | 0.1 | 1.0 | 0.01 | **0.1** | 1.0 |
| O.D.$_{600}$ | | 6.65 ± 0.23 | 4.37 ± 0.22 | 2.20 ± 0.18 | 8.47 ± 0.24 | 3.29 ± 0.11 | 1.78 ± 0.05 |
| Consumed glucose, g/L (mM) | | 9.02 ± 0.19 (50.05 ± 1.03) | 6.89 ± 0.14 (38.23 ± 0.76) | 3.75 ± 0.07 (20.80 ± 0.37) | 13.07 ± 0.16 (72.56 ± 0.90) | 4.89 ± 0.31 (27.14 ± 1.72) | 3.99 ± 0.21 (22.15 ± 1.18) |
| Lactate, g/L (mM) | | 0.31 ± 0.05 (3.46 ± 0.61) | 0.44 ± 0.03 (4.86 ± 0.32) | 0.10 ± 0.02 (1.14 ± 0.18) | 1.01 ± 0.02 (11.25 ± 0.20) | 0.14 ± 0.00 (1.52 ± 0.02) | 0.05 ± 0.00 (0.52 ± 0.02) |
| Formate, g/L (mM) | | 2.33 ± 0.12 (50.66 ± 2.60) | 1.66 ± 0.03 (36.17 ± 0.71) | 0.55 ± 0.06 (11.94 ± 1.32) | 2.54 ± 0.03 (55.28 ± 0.70) | 1.16 ± 0.01 (25.11 ± 0.16) | 0.72 ± 0.01 (15.71 ± 0.13) |
| Acetate, g/L (mM) | | 0.53 ± 0.01 (8.79 ± 0.11) | 0.41 ± 0.00 (6.75 ± 0.04) | 0.31 ± 0.00 (5.21 ± 0.01) | 0.62 ± 0.01 (10.35 ± 0.08) | 0.50 ± 0.00 (8.40 ± 0.07) | 0.34 ± 0.00 (5.66 ± 0.02) |
| Ethanol, g/L (mM) | | 3.83 ± 0.21 (83.05 ± 4.53) | 3.65 ± 0.11 (79.19 ± 2.48) | 2.63 ± 0.07 (57.00 ± 1.56) | 3.18 ± 0.01 (69.12 ± 0.17) | 3.48 ± 0.03 (75.63 ± 0.65) | 3.07 ± 0.05 (66.64 ± 1.11) |
| Butanol, g/L (mM) | | 0.36 ± 0.04 (4.92 ± 0.53) | 0.14 ± 0.05 (1.89 ± 0.62) | 0.09 ± 0.04 (1.24 ± 0.52) | 0.22 ± 0.00 (2.97 ± 0.04) | 0.15 ± 0.00 (1.99 ± 0.04) | 0.14 ± 0.00 (1.89 ± 0.02) |
| Ester titers, mg/L (mM) | EA | 2.7 ± 1.1 (0.03 ± 0.01) | 2.4 ± 0.0 (0.03 ± 0.00) | 1.1 ± 1.0 (0.01 ± 0.01) | 1.8 ± 0.5 (0.02 ± 0.01) | 1.3 ± 0.6 (0.01 ± 0.01) | 0.6 ± 0.2 (0.01 ± 0.00) |
| | EB | 77.9 ± 24.3 (0.67 ± 0.21) | 229.8 ± 72.7 (1.98 ± 0.63) | 175.6 ± 12.8 (1.51 ± 0.11) | 43.7 ± 4.6 (0.38 ± 0.04) | 246.2 ± 72.6 (2.12 ± 0.63) | 121.1 ± 4.4 (1.04 ± 0.04) |
| | BA | 0.8 ± 0.2 (0.01 ± 0.00) | 0.5 ± 0.1 (0.00 ± 0.00) | *n.d.* | 0.4 ± 0.6 (0.00 ± 0.00) | 0.2 ± 0.3 (0.00 ± 0.00) | *n.d.* |
| | BB | 36.2 ± 12.7 (0.25 ± 0.09) | 89.1 ± 28.7 (0.62 ± 0.20) | 47.0 ± 10.0 (0.33 ± 0.07) | 21.4 ± 2.5 (0.15 ± 0.02) | **127.4 ± 32.5 (0.88 ± 0.23)** | 57.9 ± 1.6 (0.40 ± 0.01) |
| | Ester total | 117.6 ± 38.3 (0.96 ± 0.31) | 321.8 ± 101.6 (2.63 ± 0.83) | 223.7 ± 16.9 (1.85 ± 0.13) | 67.2 ± 6.0 (0.55 ± 0.05) | 375.1 ± 105.5 (3.02 ± 0.86) | 179.6 ± 4.9 (1.45 ± 0.04) |
| Specific ester productivity, mg/gDCW/h (µM/gDCW/h) | EA | 0.0 ± 0.0 (0.39 ± 0.17) | 0.1 ± 0.0 (0.52 ± 0.01) | 0.0 ± 0.0 (0.50 ± 0.45) | 0.0 ± 0.0 (0.21 ± 0.06) | 0.0 ± 0.0 (0.39 ± 0.17) | 0.0 ± 0.0 (0.32 ± 0.10) |
| | EB | 1.0 ± 0.3 (8.66 ± 2.69) | 4.4 ± 1.4 (37.92 ± 12.40) | 6.9 ± 0.9 (59.55 ± 7.68) | 0.5 ± 0.1 (3.88 ± 0.41) | 6.4 ± 1.9 (55.42 ± 16.55) | 5.9 ± 0.3 (50.34 ± 2.96) |
| | BA | 0.0 ± 0.0 (0.09 ± 0.03) | 0.0 ± 0.0 (0.09 ± 0.02) | *n.a.* | 0.0 ± 0.0 (0.04 ± 0.05) | 0.0 ± 0.0 (0.04 ± 0.06) | *n.a.* |
| | BB | 0.5 ± 0.2 (3.25 ± 1.15) | 1.7 ± 0.6 (11.84 ± 3.94) | 1.8 ± 0.3 (12.68 ± 1.72) | 0.2 ± 0.0 (1.53 ± 0.18) | **3.3 ± 0.9 (23.08 ± 5.89)** | 2.8 ± 0.2 (19.38 ± 1.02) |
| | Ester total | 1.5 ± 0.5 (12.40 ± 4.02) | 6.2 ± 2.0 (50.37 ± 16.37) | 8.8 ± 0.9 (72.73 ± 7.68) | 0.7 ± 0.1 (5.65 ± 0.48) | 9.8 ± 2.8 (78.93 ± 22.58) | 8.7 ± 0.5 (70.04 ± 3.82) |
| Ester yields, mg/g glucose (mM/M) | EA | 0.3 ± 0.1 (0.61 ± 0.26) | 0.4 ± 0.0 (0.71 ± 0.00) | 0.3 ± 0.3 (0.59 ± 0.52) | 0.1 ± 0.0 (0.28 ± 0.08) | 0.3 ± 0.1 (0.55 ± 0.27) | 0.2 ± 0.0 (0.30 ± 0.08) |
| | EB | 8.7 ± 2.8 (13.45 ± 4.41) | 33.0 ± 9.9 (51.13 ± 15.42) | 46.9 ± 2.9 (72.68 ± 4.51) | 3.3 ± 0.3 (5.18 ± 0.46) | 51.0 ± 17.9 (79.03 ± 27.69) | 30.4 ± 1.3 (47.13 ± 2.03) |
| | BA | 0.1 ± 0.0 (0.14 ± 0.04) | 0.1 ± 0.0 (0.12 ± 0.03) | *n.a.* | 0.0 ± 0.0 (0.05 ± 0.07) | 0.0 ± 0.1 (0.05 ± 0.09) | *n.a.* |
| | BB | 4.0 ± 1.5 (5.05 ± 1.85) | 12.8 ± 3.9 (15.96 ± 4.90) | 12.5 ± 2.5 (15.63 ± 3.08) | 1.6 ± 0.2 (2.05 ± 0.20) | **26.3 ± 8.0 (32.87 ± 10.04)** | 14.6 ± 1.2 (18.18 ± 1.43) |
| | Ester total | 13.1 ± 4.5 (19.25 ± 6.55) | 46.2 ± 13.9 (67.93 ± 20.34) | 59.7 ± 3.5 (88.90 ± 5.05) | 5.2 ± 0.4 (7.55 ± 0.51) | 77.6 ± 26.0 (112.50 ± 37.96) | 45.1 ± 2.4 (65.61 ± 3.37) |
| Culture time (h) | | 24 | 24 | 24 | 24 | 24 | 24 |

**Supplementary Table S10.** Summary of solubility engineering of ATF1$_{Sc}$. Key strategies are in bold; Abbreviations: *n.d.*: not detected.

| Target products | Expressed enzymes/pathways | Final O.D.$_{600}$ (24h) | Ester titer (mg/L (mM)) | | | Residual alcohols (g/L (mM)) | Conversion (%) |
|---|---|---|---|---|---|---|---|
| | | | EA | BA | Total | | |
| EA (Ethanol doping) | ATF1$_{Sc}$ | 6.17 ± 0.36 | 10.4 ± 0.9 (0.12 ± 0.01) | *n.d.* | 10.4 ± 0.9 (0.12 ± 0.01) | 4.84 ± 0.26 (105.13 ± 5.58)[a] | 0.1[b] |
| | **ATF1$_{Sc}^{opt}$** | 6.34 ± 0.20 | 188.1 ± 25.0 (2.13 ± 0.28) | *n.d.* | 188.1 ± 25.0 (2.13 ± 0.28) | 4.18 ± 0.05 (90.73 ± 1.02)[a] | **2.3[b]** |
| | **MBP_ATF1$_{Sc}$** | 6.93 ± 0.33 | 84.5 ± 11.5 (0.96 ± 0.13) | *n.d.* | 84.5 ± 11.5 (0.96 ± 0.13) | 4.32 ± 0.18 (93.80 ± 3.83)[a] | **1.0[b]** |
| | **NusA_ATF1$_{Sc}$** | 6.28 ± 0.25 | 104.4 ± 3.0 (1.19 ± 0.03) | *n.d.* | 104.4 ± 3.0 (1.19 ± 0.03) | 4.12 ± 0.24 (89.53 ± 5.18)[a] | **1.3[b]** |
| | **TrxA_ATF1$_{Sc}$** | 6.87 ± 0.08 | 110.6 ± 8.1 (1.25 ± 0.09) | *n.d.* | 110.6 ± 8.1 (1.25 ± 0.09) | 4.32 ± 0.22 (93.84 ± 4.86)[a] | **1.3[b]** |
| | ATF1$_{Sc}$+Tf | 6.26 ± 0.17 | 14.6 ± 0.8 (0.17 ± 0.01) | *n.d.* | 14.6 ± 0.8 (0.17 ± 0.01) | 5.23 ± 0.16 (113.52 ± 3.46)[a] | 0.2[b] |
| | ATF1$_{Sc}$+GroES/GroEL | 6.91 ± 0.23 | 17.6 ± 5.3 (0.20 ± 0.06) | *n.d.* | 17.6 ± 5.3 (0.20 ± 0.06) | 5.39 ± 0.23 (117.05 ± 5.04)[a] | 0.2[b] |
| | ATF1$_{Sc}$+GroES/GroEL/Tf | 6.25 ± 0.25 | 18.9 ± 4.1 (0.21 ± 0.05) | *n.d.* | 18.9 ± 4.1 (0.21 ± 0.05) | 5.11 ± 0.31 (110.98 ± 6.76)[a] | 0.2[b] |
| | ATF1$_{Sc}$+DnaK/DnaJ/GrpE | 5.69 ± 0.37 | 12.5 ± 2.4 (0.14 ± 0.03) | *n.d.* | 12.5 ± 2.4 (0.14 ± 0.03) | 5.27 ± 0.16 (114.36 ± 3.48)[a] | 0.1[b] |
| | ATF1$_{Sc}$+DnaK/DnaJ/GrpE+ /GroES/GroEL | 5.44 ± 0.13 | 12.4 ± 1.9 (0.14 ± 0.02) | *n.d.* | 12.4 ± 1.9 (0.14 ± 0.02) | 5.05 ± 0.04 (109.52 ± 0.81)[a] | 0.1[b] |
| BA (Butanol doping) | ATF1$_{Sc}$ | 5.41 ± 0.20 | 3.2 ± 1.4 (0.04 ± 0.02) | 370.7 ± 42.7 (3.19 ± 0.37) | 373.9 ± 44.0 (3.23 ± 0.38) | 1.73 ± 0.18 (23.33 ± 2.49)[c] | 12.0[d] |
| | **ATF1$_{Sc}^{opt}$** | 5.66 ± 0.26 | 44.7 ± 12.3 (0.51 ± 0.14) | 2,256.1 ± 222.8 (19.42 ± 1.92) | 2,300.8 ± 228.3 (19.93 ± 1.98) | 0.24 ± 0.08 (3.19 ± 1.03)[c] | **85.9[d]** |
| | **MBP_ATF1$_{Sc}$** | 6.25 ± 0.03 | 25.7 ± 8.9 (0.29 ± 0.09) | 2,062.3 ± 165.3 (17.75 ± 1.42) | 2,088.0 ± 173.5 (18.05 ± 1.52) | 0.45 ± 0.13 (6.06 ± 1.72)[c] | **74.6[d]** |
| | **NusA_ATF1$_{Sc}$** | 5.63 ± 0.31 | 31.5 ± 8.2 (0.36 ± 0.09) | 2,081.2 ± 97.5 (17.92 ± 0.84) | 2,112.6 ± 95.6 (18.27 ± 0.82) | 0.34 ± 0.15 (4.61 ± 1.99)[c] | **79.5[d]** |
| | **TrxA_ATF1$_{Sc}$** | 6.49 ± 0.20 | 33.1 ± 12.5 (0.38 ± 0.14) | 2,095.3 ± 191.9 (18.04 ± 1.65) | 2,128.3 ± 187.9 (18.41 ± 1.61) | 0.21 ± 0.02 (2.79 ± 0.20)[c] | **86.6[d]** |
| | ATF1$_{Sc}$+Tf | 5.49 ± 0.09 | 7.1 ± 1.2 (0.08 ± 0.01) | 480.4 ± 73.8 (4.14 ± 0.64) | 487.5 ± 74.8 (4.22 ± 0.65) | 1.71 ± 0.09 (23.09 ± 1.16)[c] | 15.2[d] |
| | ATF1$_{Sc}$+GroES/GroEL | 5.94 ± 0.16 | 8.0 ± 0.7 (0.09 ± 0.01) | 635.1 ± 165.3 (5.47 ± 1.42) | 643.0 ± 164.6 (5.56 ± 1.42) | 1.66 ± 0.10 (22.35 ± 1.36)[c] | 19.7[d] |
| | ATF1$_{Sc}$+GroES/GroEL/Tf | 5.72 ± 0.50 | 6.3 ± 0.6 (0.07 ± 0.01) | 380.3 ± 67.0 (3.27 ± 0.58) | 386.6 ± 66.4 (3.35 ± 0.57) | 1.96 ± 0.10 (26.50 ± 1.36)[c] | 11.0[d] |
| | ATF1$_{Sc}$+DnaK/DnaJ/GrpE | 5.27 ± 0.26 | 4.5 ± 1.1 (0.05 ± 0.01) | 500.1 ± 178.3 (4.31 ± 1.53) | 504.7 ± 177.3 (4.36 ± 1.52) | 1.93 ± 0.30 (25.99 ± 4.07)[c] | 14.2[d] |
| | ATF1$_{Sc}$+DnaK/DnaJ/GrpE+ /GroES/GroEL | 4.68 ± 0.21 | 4.1 ± 1.5 (0.05 ± 0.02) | 484.1 ± 17.1 (4.17 ± 0.15) | 488.2 ± 18.6 (4.21 ± 0.16) | 1.98 ± 0.17 (26.74 ± 2.35)[c] | 13.5[d] |

[a]Concentration of residual ethanol; [b]Conversion of ethanol into EA: (EA produced)/(EA produced + ethanol remained)*100, (mole/mole).
[c]Concentration of residual butanol; [d]Conversion of butanol into BA: (BA produced)/(BA produced + butanol remained)*100, (mole/mole).

**Supplementary Table S11.** Summary of solubility engineering of SAAT$_{Fa}$. Key strategy is in bold.

| Target products | Expressed enzymes/pathways | Final O.D.$_{600}$ (24h) | Ester titer (mg/L (mM)) | | | | | Residual alcohols (g/L (mM)) | Conversion (%) |
|---|---|---|---|---|---|---|---|---|---|
| | | | EA | EB | BA | BB | Total | | |
| EB (Ethanol doping) | BCoA+SAAT$_{Fa}$ | 6.87 ± 0.01 | 1.8 ± 0.2<br>(0.02 ± 0.00) | 46.5 ± 5.3<br>(0.40 ± 0.05) | 0.3 ± 0.1<br>(0.00 ± 0.00) | 11.4 ± 1.9<br>(0.08 ± 0.01) | 59.9 ± 7.3<br>(0.50 ± 0.06) | 5.98 ± 0.15<br>(129.88 ± 3.36)[a] | 0.3[b] |
| | BCoA+SAAT$_{Fa}^{opt}$ | 5.52 ± 0.21 | 1.0 ± 0.7<br>(0.01 ± 0.01) | 55.2 ± 16.3<br>(0.47 ± 0.14) | 0.2 ± 0.0<br>(0.00 ± 0.00) | 14.0 ± 5.4<br>(0.10 ± 0.04) | 70.3 ± 21.1<br>(0.58 ± 0.17) | 6.48 ± 0.04<br>(140.76 ± 0.90)[a] | 0.3[b] |
| | BCoA+MBP_SAAT$_{Fa}$ | 6.33 ± 0.06 | 1.4 ± 0.1<br>(0.02 ± 0.00) | 24.0 ± 1.6<br>(0.21 ± 0.01) | 0.1 ± 0.0<br>(0.00 ± 0.00) | 6.5 ± 1.0<br>(0.05 ± 0.01) | 32.0 ± 2.6<br>(0.27 ± 0.02) | 6.32 ± 0.20<br>(137.17 ± 4.27)[a] | 0.2[b] |
| | BCoA+NusA_SAAT$_{Fa}$ | 6.54 ± 0.25 | 1.5 ± 0.1<br>(0.02 ± 0.00) | 13.0 ± 1.0<br>(0.11 ± 0.01) | 0.2 ± 0.1<br>(0.00 ± 0.00) | 3.0 ± 0.5<br>(0.02 ± 0.00) | 17.7 ± 1.5<br>(0.15 ± 0.01) | 6.43 ± 0.09<br>(139.50 ± 1.88)[a] | 0.1[b] |
| | BCoA+TrxA_SAAT$_{Fa}$ | 6.22 ± 0.19 | 1.8 ± 0.3<br>(0.02 ± 0.00) | 30.8 ± 2.9<br>(0.27 ± 0.02) | 0.2 ± 0.0<br>(0.00 ± 0.00) | 5.8 ± 0.8<br>(0.04 ± 0.01) | 38.6 ± 3.9<br>(0.33 ± 0.03) | 6.23 ± 0.23<br>(135.27 ± 5.09)[a] | 0.2[b] |
| | BCoA+SAAT$_{Fa}$+Tf | 6.62 ± 0.19 | 4.1 ± 0.8<br>(0.05 ± 0.01) | 25.2 ± 4.7<br>(0.22 ± 0.04) | 0.3 ± 0.1<br>(0.00 ± 0.00) | 3.0 ± 0.0<br>(0.02 ± 0.00) | 32.6 ± 5.4<br>(0.29 ± 0.05) | 7.15 ± 0.49<br>(155.11 ± 10.69)[a] | 0.1[b] |
| | **BCoA+SAAT$_{Fa}$+GroES/GroEL** | 6.82 ± 0.21 | 6.3 ± 1.3<br>(0.07 ± 0.01) | 464.2 ± 8.9<br>(4.00 ± 0.08) | 0.4 ± 0.0<br>(0.00 ± 0.00) | 66.7 ± 1.6<br>(0.46 ± 0.01) | 537.6 ± 7.4<br>(4.53 ± 0.07) | 6.90 ± 0.32<br>(149.78 ± 6.91)[a] | **2.6[b]** |
| | BCoA+SAAT$_{Fa}$+GroES/GroEL/Tf | 6.43 ± 0.14 | 9.0 ± 1.2<br>(0.10 ± 0.01) | 250.2 ± 35.8<br>(2.15 ± 0.31) | 0.7 ± 0.1<br>(0.01 ± 0.00) | 39.1 ± 5.6<br>(0.27 ± 0.04) | 299.0 ± 42.6<br>(2.53 ± 0.36) | 6.53 ± 0.07<br>(141.82 ± 1.45)[a] | 1.5[b] |
| | BCoA+SAAT$_{Fa}$+DnaK/DnaJ/GrpE | 5.04 ± 0.12 | 2.7 ± 0.9<br>(0.03 ± 0.01) | 10.3 ± 1.1<br>(0.09 ± 0.01) | 0.3 ± 0.0<br>(0.00 ± 0.00) | 1.5 ± 0.2<br>(0.01 ± 0.00) | 14.8 ± 2.1<br>(0.13 ± 0.02) | 5.64 ± 0.14<br>(122.39 ± 3.03)[a] | 0.1[b] |
| | BCoA+SAAT$_{Fa}$+DnaK/DnaJ/GrpE+<br>/GroES/GroEL | 5.68 ± 0.04 | 4.8 ± 1.2<br>(0.05 ± 0.01) | 330.5 ± 44.4<br>(2.85 ± 0.38) | 0.3 ± 0.0<br>(0.00 ± 0.00) | 44.8 ± 6.1<br>(0.31 ± 0.04) | 380.4 ± 48.6<br>(3.21 ± 0.41) | 6.53 ± 0.35<br>(141.69 ± 7.68)[a] | 2.0[b] |
| BB (Butanol doping) | BCoA+SAAT$_{Fa}$ | 6.21 ± 0.20 | 0.2 ± 0.0<br>(0.00 ± 0.00) | 10.5 ± 0.9<br>(0.09 ± 0.01) | 17.9 ± 1.3<br>(0.15 ± 0.01) | 207.8 ± 5.1<br>(1.44 ± 0.04) | 236.5 ± 5.4<br>(1.69 ± 0.04) | 2.51 ± 0.16<br>(33.93 ± 2.21)[c] | 4.1[d] |
| | BCoA+SAAT$_{Fa}^{opt}$ | 4.92 ± 0.15 | 0.0 ± 0.1<br>(0.00 ± 0.00) | 7.2 ± 2.7<br>(0.06 ± 0.02) | 21.1 ± 9.3<br>(0.18 ± 0.08) | 344.2 ± 171.6<br>(2.39 ± 1.19) | 372.5 ± 183.6<br>(2.63 ± 1.29) | 2.54 ± 0.14<br>(34.31 ± 1.86)[c] | 6.5[d] |
| | BCoA+MBP_SAAT$_{Fa}$ | 5.78 ± 0.36 | 0.2 ± 0.0<br>(0.00 ± 0.00) | 6.4 ± 0.4<br>(0.06 ± 0.00) | 16.5 ± 0.7<br>(0.14 ± 0.01) | 147.1 ± 14.6<br>(1.02 ± 0.10) | 170.2 ± 14.9<br>(1.22 ± 0.10) | 2.45 ± 0.27<br>(33.09 ± 3.70)[c] | 3.0[d] |
| | BCoA+NusA_SAAT$_{Fa}$ | 5.71 ± 0.12 | 0.1 ± 0.0<br>(0.00 ± 0.00) | 2.9 ± 0.2<br>(0.02 ± 0.00) | 14.2 ± 0.2<br>(0.12 ± 0.00) | 81.5 ± 7.2<br>(0.57 ± 0.05) | 98.7 ± 7.0<br>(0.71 ± 0.05) | 2.53 ± 0.21<br>(34.13 ± 2.79)[c] | 1.6[d] |
| | BCoA+TrxA_SAAT$_{Fa}$ | 5.87 ± 0.52 | 0.2 ± 0.1<br>(0.00 ± 0.00) | 10.8 ± 4.4<br>(0.09 ± 0.04) | 20.2 ± 4.5<br>(0.17 ± 0.04) | 189.2 ± 64.4<br>(1.31 ± 0.45) | 220.4 ± 73.0<br>(1.58 ± 0.52) | 2.26 ± 0.21<br>(30.43 ± 2.78)[c] | 4.1[d] |
| | BCoA+SAAT$_{Fa}$+Tf | 6.03 ± 0.39 | 0.9 ± 0.4<br>(0.01 ± 0.00) | 11.0 ± 0.9<br>(0.09 ± 0.01) | 59.1 ± 14.8<br>(0.51 ± 0.13) | 152.0 ± 28.8<br>(1.05 ± 0.20) | 223.0 ± 44.3<br>(1.67 ± 0.33) | 2.52 ± 0.41<br>(33.97 ± 5.47)[c] | 3.0[d] |
| | **BCoA+SAAT$_{Fa}$+GroES/GroEL** | 5.66 ± 0.34 | 10.5 ± 3.1<br>(0.12 ± 0.04) | 195.7 ± 15.4<br>(1.69 ± 0.13) | 244.0 ± 101.4<br>(2.10 ± 0.87) | 1,710.1 ± 256.6<br>(11.86 ± 1.78) | 2,160.4 ± 349.7<br>(15.76 ± 2.59) | 1.51 ± 0.33<br>(20.35 ± 4.48)[c] | **36.8[d]** |
| | BCoA+SAAT$_{Fa}$+GroES/GroEL/Tf | 5.57 ± 0.10 | 3.1 ± 0.3<br>(0.04 ± 0.00) | 81.3 ± 6.2<br>(0.70 ± 0.05) | 120.3 ± 16.9<br>(1.04 ± 0.15) | 822.1 ± 109.0<br>(5.70 ± 0.76) | 1,026.9 ± 131.0<br>(7.47 ± 0.95) | 1.79 ± 0.12<br>(24.09 ± 1.62)[c] | 19.1[d] |
| | BCoA+SAAT$_{Fa}$+DnaK/DnaJ/GrpE | 4.65 ± 0.06 | 0.6 ± 0.1<br>(0.01 ± 0.00) | 3.5 ± 0.4<br>(0.03 ± 0.00) | 39.5 ± 2.4<br>(0.34 ± 0.02) | 48.3 ± 5.3<br>(0.34 ± 0.04) | 91.9 ± 7.6<br>(0.71 ± 0.06) | 2.40 ± 0.11<br>(32.43 ± 1.44)[c] | 1.0[d] |
| | BCoA+SAAT$_{Fa}$+DnaK/DnaJ/GrpE+<br>/GroES/GroEL | 5.16 ± 0.11 | 1.6 ± 0.1<br>(0.02 ± 0.00) | 52.0 ± 8.0<br>(0.45 ± 0.07) | 56.6 ± 7.5<br>(0.49 ± 0.06) | 565.0 ± 50.4<br>(3.92 ± 0.35) | 675.1 ± 52.7<br>(4.87 ± 0.37) | 2.16 ± 0.21<br>(29.12 ± 2.80)[c] | 11.9[d] |

[a]Concentration of residual ethanol; [b]Conversion of ethanol into EB: (EB produced)/(EB produced + ethanol remained)*100, (mole/mole).
[c]Concentration of residual butanol; [d]Conversion of butanol into BB: (BB produced)/(BB produced + butanol remained)*100, (mole/mole).

**Supplementary Table S12.** Summary of BA production with strategies for improving solubility of ATF1$_{Sc}$. Key strain is in bold; Abbreviations: *n.d.*: not detected. *n.a.*: not applicable.

| Target product | | BA | | | |
|---|---|---|---|---|---|
| Strains | | EcJWBA7 | EcJWBA8 | EcJWBA9 | **EcJWBA10** |
| Strategies for improving solubility of ATF1$_{Sc}$ | | Codon optimization | Codon opt. +MBP-tag (N'-terminus) | Codon opt. +NusA-tag (N'-terminus) | **Codon opt. +TrxA-tag (N'-terminus)** |
| O.D.$_{600}$ | | 5.35 ± 0.18 | 5.57 ± 0.05 | 4.45 ± 0.11 | 5.87 ± 0.23 |
| Consumed glucose, g/L (mM) | | 5.68 ± 0.14 (31.51 ± 0.76) | 6.54 ± 0.12 (36.29 ± 0.67) | 5.31 ± 0.18 (29.50 ± 1.01) | 5.36 ± 0.06 (29.75 ± 0.35) |
| Lactate, g/L (mM) | | 0.14 ± 0.01 (1.56 ± 0.06) | 0.21 ± 0.03 (2.28 ± 0.36) | 0.06 ± 0.00 (0.62 ± 0.03) | 0.14 ± 0.05 (1.51 ± 0.51) |
| Formate, g/L (mM) | | 1.60 ± 0.05 (34.72 ± 0.99) | 1.78 ± 0.09 (38.75 ± 1.86) | 1.81 ± 0.04 (39.38 ± 0.88) | 1.51 ± 0.56 (32.89 ± 12.25) |
| Acetate, g/L (mM) | | 1.09 ± 0.02 (18.11 ± 0.28) | 0.79 ± 0.00 (13.21 ± 0.04) | 0.66 ± 0.01 (11.00 ± 0.10) | 0.88 ± 0.13 (14.62 ± 2.21) |
| Ethanol, g/L (mM) | | 2.17 ± 0.06 (47.08 ± 1.32) | 3.09 ± 0.06 (67.12 ± 1.37) | 3.23 ± 0.06 (70.22 ± 1.40) | 2.49 ± 0.26 (54.07 ± 5.54) |
| Butanol, g/L (mM) | | 0.06 ± 0.00 (0.82 ± 0.02) | 0.17 ± 0.01 (2.27 ± 0.11) | 0.24 ± 0.01 (3.24 ± 0.20) | 0.11 ± 0.02 (1.48 ± 0.20) |
| Ester titers, mg/L (mM) | EA | 18.7 ± 1.5 (0.21 ± 0.02) | 11.6 ± 2.0 (0.13 ± 0.02) | 6.4 ± 0.6 (0.07 ± 0.01) | 16.7 ± 1.8 (0.19 ± 0.02) |
| | EB | *n.d.* | *n.d.* | *n.d.* | *n.d.* |
| | BA | 51.7 ± 7.1 (0.44 ± 0.06) | 79.3 ± 9.8 (0.68 ± 0.08) | 76.1 ± 6.2 (0.66 ± 0.05) | **89.5 ± 14.8 (0.77 ± 0.13)** |
| | BB | *n.d.* | *n.d.* | *n.d.* | *n.d.* |
| | Ester total | 70.4 ± 8.8 (0.66 ± 0.08) | 90.9 ± 10.8 (0.81 ± 0.10) | 82.5 ± 6.3 (0.73 ± 0.05) | 106.1 ± 16.0 (0.96 ± 0.14) |
| Specific ester productivity, mg/gDCW/h (µM/gDCW/h) | EA | 0.3 ± 0.0 (0.00 ± 0.00) | 0.2 ± 0.0 (0.00 ± 0.00) | 0.1 ± 0.0 (0.00 ± 0.00) | 0.2 ± 0.0 (0.00 ± 0.00) |
| | EB | *n.a.* | *n.a.* | *n.a.* | *n.a.* |
| | BA | 0.8 ± 0.1 (0.01 ± 0.00) | 1.2 ± 0.1 (0.01 ± 0.00) | 1.5 ± 0.1 (0.01 ± 0.00) | **1.3 ± 0.2 (0.01 ± 0.00)** |
| | BB | *n.a.* | *n.a.* | *n.a.* | *n.a.* |
| | Ester total | 1.1 ± 0.2 (0.01 ± 0.00) | 1.4 ± 0.2 (0.01 ± 0.00) | 1.6 ± 0.1 (0.01 ± 0.00) | 1.6 ± 0.2 (0.01 ± 0.00) |
| Ester yields, mg/g glucose (mM/M) | EA | 3.3 ± 0.3 (6.74 ± 0.69) | 1.8 ± 0.3 (3.62 ± 0.55) | 1.2 ± 0.1 (2.46 ± 0.24) | 3.1 ± 0.3 (6.35 ± 0.63) |
| | EB | *n.a.* | *n.a.* | *n.a.* | *n.a.* |
| | BA | 9.1 ± 1.3 (14.12 ± 1.94) | 12.1 ± 1.4 (18.80 ± 2.23) | 14.3 ± 0.7 (22.19 ± 1.10) | **16.7 ± 2.8 (25.88 ± 4.26)** |
| | BB | *n.a.* | *n.a.* | *n.a.* | *n.a.* |
| | Ester total | 12.4 ± 1.6 (20.86 ± 2.59) | 13.9 ± 1.6 (22.42 ± 2.46) | 15.5 ± 0.7 (24.65 ± 1.01) | 19.8 ± 3.0 (32.23 ± 4.68) |
| Culture time (h) | | 24 | 24 | 24 | 24 |

**Supplementary Table S13.** Summary of production of EB with strategy for improving solubility of $SAAT_{Fa}$. The best culture condition is in bold; Abbreviations: *n.d.*: not detected. *n.a.*: not applicable.

| Target product | | EB | | | |
|---|---|---|---|---|---|
| Strains | | EcJWEB7 | | | |
| Strategy for improving solubility of $SAAT_{Fa}$ | | Co-expression of GroES/EL | | | |
| Arabinose conc. for inducing expression of GroES/EL | | 0 mg/ml | 1 mg/ml | **5 mg/ml** | 50 mg/ml |
| $O.D._{600}$ | | 3.85 ± 0.13 | 3.86 ± 0.34 | 3.47 ± 0.23 | 3.69 ± 0.10 |
| Consumed glucose, g/L (mM) | | 9.83 ± 0.10 (54.57 ± 0.54) | 9.78 ± 0.08 (54.28 ± 0.45) | 9.49 ± 0.19 (52.68 ± 1.07) | 7.71 ± 0.16 (42.81 ± 0.91) |
| Lactate, g/L (mM) | | 0.08 ± 0.00 (0.87 ± 0.01) | 0.08 ± 0.00 (0.88 ± 0.01) | 0.08 ± 0.00 (0.87 ± 0.01) | 0.09 ± 0.00 (0.97 ± 0.02) |
| Formate, g/L (mM) | | 0.26 ± 0.01 (5.74 ± 0.16) | 0.25 ± 0.01 (5.42 ± 0.15) | 0.25 ± 0.01 (5.41 ± 0.23) | 0.24 ± 0.00 (5.24 ± 0.05) |
| Acetate, g/L (mM) | | 0.24 ± 0.00 (4.03 ± 0.01) | 0.24 ± 0.00 (4.04 ± 0.01) | 0.24 ± 0.00 (4.01 ± 0.01) | 0.25 ± 0.00 (4.14 ± 0.04) |
| Ethanol, g/L (mM) | | 5.17 ± 0.03 (112.22 ± 0.60) | 5.12 ± 0.05 (111.15 ± 0.99) | 5.04 ± 0.09 (109.31 ± 2.05) | 5.45 ± 0.07 (118.26 ± 1.52) |
| Butanol, g/L (mM) | | 0.03 ± 0.00 (0.47 ± 0.04) | 0.03 ± 0.00 (0.40 ± 0.06) | 0.03 ± 0.00 (0.34 ± 0.01) | 0.03 ± 0.01 (0.43 ± 0.07) |
| Ester titers, mg/L (mM) | EA | 15.4 ± 1.1 (0.18 ± 0.01) | 29.8 ± 6.4 (0.34 ± 0.07) | 19.6 ± 11.2 (0.22 ± 0.13) | 41.0 ± 4.5 (0.47 ± 0.05) |
| | EB | 263.9 ± 51.8 (2.27 ± 0.45) | 337.6 ± 46.1 (2.91 ± 0.40) | **365.7 ± 69.2 (3.15 ± 0.60)** | 346.2 ± 59.8 (2.98 ± 0.51) |
| | BA | *n.d.* | *n.d.* | *n.d.* | *n.d.* |
| | BB | 6.4 ± 1.9 (0.04 ± 0.01) | 7.5 ± 1.3 (0.05 ± 0.01) | 8.4 ± 2.4 (0.06 ± 0.02) | 7.6 ± 1.2 (0.05 ± 0.01) |
| | Ester total | 285.7 ± 53.8 (2.49 ± 0.46) | 374.9 ± 51.1 (3.30 ± 0.45) | 393.6 ± 72.7 (3.43 ± 0.63) | 394.7 ± 62.0 (3.50 ± 0.54) |
| Specific ester productivity, mg/gDCW/h (µM/gDCW/h) | EA | 0.3 ± 0.0 (3.83 ± 0.11) | 0.7 ± 0.2 (6.15 ± 2.02) | 0.5 ± 0.3 (5.45 ± 2.90) | 1.0 ± 0.1 (9.46 ± 1.63) |
| | EB | 5.7 ± 1.5 (49.30 ± 13.11) | 6.7 ± 0.5 (57.64 ± 4.25) | **8.8 ± 1.6 (75.70 ± 14.01)** | 8.8 ± 1.4 (75.74 ± 12.07) |
| | BA | *n.a.* | *n.a.* | *n.a.* | *n.a.* |
| | BB | 0.1 ± 0.1 (0.91 ± 0.33) | 0.2 ± 0.0 (1.08 ± 0.07) | 0.2 ± 0.1 (1.41 ± 0.44) | 0.2 ± 0.0 (1.37 ± 0.13) |
| | Ester total | 6.2 ± 1.6 (54.04 ± 13.33) | 7.4 ± 0.6 (64.86 ± 5.48) | 9.5 ± 1.5 (82.56 ± 12.81) | 9.8 ± 1.4 (86.56 ± 12.44) |
| Ester yields, mg/g glucose (mM/M) | EA | 1.6 ± 0.1 (3.21 ± 0.25) | 3.1 ± 0.6 (6.23 ± 1.29) | 2.1 ± 1.2 (4.20 ± 2.35) | 5.3 ± 0.7 (10.88 ± 1.39) |
| | EB | 26.8 ± 5.1 (41.58 ± 7.83) | 34.5 ± 4.7 (53.55 ± 7.32) | **38.5 ± 6.7 (59.64 ± 10.42)** | 44.9 ± 7.5 (69.58 ± 11.59) |
| | BA | *n.a.* | *n.a.* | *n.a.* | *n.a.* |
| | BB | 0.7 ± 0.2 (0.81 ± 0.23) | 0.8 ± 0.1 (0.95 ± 0.17) | 0.9 ± 0.3 (1.10 ± 0.31) | 1.0 ± 0.2 (1.22 ± 0.18) |
| | Ester total | 29.0 ± 5.2 (45.60 ± 8.08) | 38.3 ± 5.2 (60.73 ± 8.22) | 41.4 ± 7.0 (64.94 ± 10.78) | 51.2 ± 7.8 (81.69 ± 12.14) |
| Culture time (h) | | 24 | 24 | 24 | 24 |

**Supplementary Table S14.** Summary of production of BB with strategies for improving solubility of $SAAT_{Fa}$.

| Target product | | BB | | | |
|---|---|---|---|---|---|
| Strains | | EcJWBB7 | | | |
| Strategy for improving solubility of $SAAT_{Fa}$ | | Co-expression of GroES/EL | | | |
| Arabinose conc. for inducing expression of GroES/EL | | 0 mg/ml | 1 mg/ml | 5 mg/ml | 50 mg/ml |
| $O.D._{600}$ | | 2.97 ± 0.47 | 2.09 ± 0.08 | 2.27 ± 0.10 | 3.14 ± 0.16 |
| Consumed glucose, g/L (mM) | | 3.72 ± 2.80 (20.67 ± 15.52) | 1.58 ± 0.98 (8.77 ± 5.43) | 0.95 ± 0.87 (5.26 ± 4.83) | 1.10 ± 0.61 (6.08 ± 3.36) |
| Lactate, g/L (mM) | | 0.19 ± 0.02 (2.08 ± 0.23) | 0.12 ± 0.00 (1.29 ± 0.05) | 0.08 ± 0.00 (0.85 ± 0.02) | 0.14 ± 0.00 (1.60 ± 0.03) |
| Formate, g/L (mM) | | 1.03 ± 0.35 (22.47 ± 7.54) | 0.76 ± 0.02 (16.49 ± 0.36) | 0.81 ± 0.01 (17.58 ± 0.26) | 0.93 ± 0.03 (20.30 ± 0.59) |
| Acetate, g/L (mM) | | 0.48 ± 0.03 (8.05 ± 0.55) | 0.41 ± 0.01 (6.75 ± 0.17) | 0.44 ± 0.01 (7.40 ± 0.15) | 0.49 ± 0.01 (8.18 ± 0.24) |
| Ethanol, g/L (mM) | | 2.71 ± 0.23 (58.92 ± 4.99) | 1.99 ± 0.02 (43.14 ± 0.54) | 2.11 ± 0.08 (45.70 ± 1.67) | 2.85 ± 0.09 (61.81 ± 1.90) |
| Butanol, g/L (mM) | | 0.02 ± 0.01 (0.31 ± 0.13) | 0.06 ± 0.04 (0.75 ± 0.48) | 0.07 ± 0.01 (0.90 ± 0.12) | 0.04 ± 0.04 (0.53 ± 0.52) |
| Ester titers, mg/L (mM) | EA | 6.0 ± 1.6 (0.07 ± 0.02) | 6.1 ± 1.0 (0.07 ± 0.01) | 8.22 ± 1.6 (0.09 ± 0.02) | 6.9 ± 2.6 (0.08 ± 0.03) |
| | EB | 155.8 ± 7.7 (1.34 ± 0.07) | 147.3 ± 15.1 (1.27 ± 0.13) | 240.4 ± 14.4 (2.07 ± 0.12) | 224.2 ± 14.0 (1.93 ± 0.12) |
| | BA | 0.3 ± 0.1 (0.00 ± 0.00) | 0.6 ± 0.1 (0.01 ± 0.00) | 1.9 ± 0.1 (0.02 ± 0.00) | 1.5 ± 0.1 (0.01 ± 0.00) |
| | BB | 25.6 ± 10.7 (0.18 ± 0.07) | 30.0 ± 4.8 (0.21 ± 0.03) | 51.4 ± 3.4 (0.36 ± 0.02) | 42.1± 4.1 (0.29 ± 0.03) |
| | Ester total | 187.7 ± 16.2 (1.59 ± 0.12) | 184.0 ± 20.3 (1.55 ± 0.17) | 302.0 ± 18.7 (2.54 ± 0.16) | 274.7 ± 15.8 (2.31 ± 0.13) |
| Specific ester productivity, mg/gDCW/h (μM/gDCW/h) | EA | 0.2 ± 0.1 (2.02 ± 0.66) | 0.3 ± 0.1 (2.84 ± 0.54) | 0.3 ± 0.1 (3.53 ± 0.61) | 0.2 ± 0.1 (2.17 ± 0.89) |
| | EB | 4.6 ± 0.5 (39.32 ± 4.32) | 6.1 ± 0.8 (52.28 ± 7.06) | 9.1 ± 0.2 (78.52 ± 1.82) | 6.2 ± 0.6 (53.10 ± 5.45) |
| | BA | 0.0 ± 0.0 (0.07 ± 0.02) | 0.0 ± 0.0 (0.22 ± 0.03) | 0.1 ± 0.0 (0.61 ± 0.03) | 0.0 ± 0.0 (0.35 ± 0.01) |
| | BB | 0.7 ± 0.3 (5.13 ± 2.01) | 1.2 ± 0.2 (8.58 ± 1.63) | 2.0 ± 0.1 (13.53 ± 0.35) | 1.2 ± 0.1 (8.02 ± 0.90) |
| | Ester total | 5.5 ± 0.6 (46.54 ± 5.15) | 7.6 ± 1.1 (63.92 ± 9.00) | 11.5 ± 0.3 (96.19 ± 2.04) | 7.6 ± 0.7 (63.64 ± 6.20) |
| Ester yields, mg/g glucose (mM/M) | EA | 1.2 ± 0.2 (2.46 ± 0.42) | 2.6 ± 0.3 (5.29 ± 0.54) | 5.4 ± 0.9 (11.06 ± 1.77) | 6.5 ± 3.8 (13.26 ± 7.70) |
| | EB | 28.9 ± 4.5 (44.74 ± 6.98) | 67.7 ± 2.0 (104.94 ± 3.10) | 161.5 ± 17.0 (250.51 ± 26.32) | 166.5 ± 62.1 (258.27 ± 96.38) |
| | BA | 0.0 ± 0.0 (0.07 ± 0.03) | 0.3 ± 0.0 (0.45 ± 0.01) | 1.3 ± 0.2 (1.94 ± 0.27) | 1.1 ± 0.3 (1.76 ± 0.41) |
| | BB | 4.2 ± 2.7 (5.28 ± 3.37) | 13.5 ± 1.5 (16.91 ± 1.85) | 34.4 ± 3.4 (43.02 ± 4.24) | 30.1 ± 9.0 (37.66 ± 11.29) |
| | Ester total | 34.3 ± 7.0 (52.54 ± 9.96) | 84.1 ± 3.2 (127.59 ± 4.42) | 202.6 ± 19.7 (306.54 ± 29.06) | 204.3 ± 75.2 (310.95 ± 115.78) |
| Culture time (h) | | 24 | 24 | 24 | 24 |

**Supplementary Table S15.** Summary of BA and BB production with strategies for improving solubility of AdhE2$_{Ca}$. Key strain is in bold; Abbreviations: *n.d.*: not detected. *n.a.*: not applicable.

| Target product | | BA | | | | BB |
|---|---|---|---|---|---|---|
| Strains | | EcJWBA11 | EcJWBA12 | EcJWBA13 | **EcJWBA14** | **EcJWBB8** |
| Strategies for improving solubility of AdhE2$_{Ca}$ | | Codon optimization | Codon opt. +MBP-tag (N'-terminus) | Codon opt. +NusA-tag (N'-terminus) | **Codon opt. +TrxA-tag (N'-terminus)** | **Codon opt. +TrxA-tag (N'-terminus)** |
| O.D.$_{600}$ | | 4.19 ± 0.14 | 3.70 ± 0.19 | 3.85 ± 0.20 | 4.28 ± 0.19 | 4.97 ± 0.26 |
| Consumed glucose, g/L (mM) | | 8.23 ± 0.71 (45.67 ± 3.95) | 2.13 ± 1.55 (11.82 ± 8.62) | 2.21 ± 0.04 (12.25 ± 0.19) | 2.34 ± 0.06 (13.00 ± 0.35) | 4.00 ± 2.37 (22.20 ± 13.13) |
| Lactate, g/L (mM) | | 0.04 ± 0.00 (0.42 ± 0.02) | 0.06 ± 0.01 (0.65 ± 0.09) | 0.03 ± 0.00 (0.37 ± 0.03) | 0.05 ± 0.00 (0.51 ± 0.04) | 0.10 ± 0.01 (1.15 ± 0.07) |
| Formate, g/L (mM) | | 0.60 ± 0.12 (12.98 ± 2.63) | 0.67 ± 0.07 (14.62 ± 1.63) | 0.63 ± 0.03 (13.78 ± 0.62) | 0.69 ± 0.02 (15.08 ± 0.34) | 0.92 ± 0.09 (19.89 ± 1.96) |
| Acetate, g/L (mM) | | 0.69 ± 0.05 (11.43 ± 0.75) | 0.75 ± 0.01 (12.57 ± 0.16) | 0.80 ± 0.01 (13.24 ± 0.24) | 0.78 ± 0.01 (12.99 ± 0.13) | 0.47 ± 0.03 (7.77 ± 0.54) |
| Ethanol, g/L (mM) | | 1.15 ± 0.03 (24.90 ± 0.71) | 1.60 ± 0.09 (34.83 ± 1.86) | 1.33 ± 0.02 (28.81 ± 0.37) | 1.40 ± 0.02 (30.33 ± 0.42) | 2.53 ± 0.05 (54.84 ± 1.02) |
| Butanol, g/L (mM) | | 0.10 ± 0.00 (1.30 ± 0.05) | 0.14 ± 0.00 (1.93 ± 0.06) | 0.16 ± 0.01 (2.11 ± 0.15) | 0.20 ± 0.01 (2.76 ± 0.13) | 0.18 ± 0.00 (2.40 ± 0.02) |
| Ester titers, mg/L (mM) | EA | 12.4 ± 1.5 (0.14 ± 0.02) | 9.2 ± 0.5 (0.10 ± 0.01) | 9.7 ± 4.1 (0.11 ± 0.05) | 2.0 ± 0.4 (0.02 ± 0.00) | 2.7 ± 1.0 (0.03 ± 0.01) |
| | EB | *n.d.* | *n.d.* | *n.d.* | *n.d.* | 156.3 ± 22.4 (1.35 ± 0.19) |
| | BA | 80.3 ± 9.0 (0.69 ± 0.08) | 54.2 ± 4.9 (0.47 ± 0.04) | 82.8 ± 15.5 (0.71 ± 0.13) | **203.0 ± 5.7 (1.75 ± 0.05)** | 4.1 ± 0.5 (0.04 ± 0.00) |
| | BB | *n.d.* | *n.d.* | *n.d.* | *n.d.* | **167.3 ± 18.2 (1.16 ± 0.13)** |
| | Ester total | 92.8 ± 10.5 (0.83 ± 0.09) | 63.3 ± 4.8 (0.57 ± 0.04) | 92.5 ± 19.6 (0.82 ± 0.18) | 205.0 ± 6.1 (1.77 ± 0.05) | 330.5 ± 16.3 (2.57 ± 0.14) |
| Specific ester productivity, mg/gDCW/h (μM/gDCW/h) | EA | 0.3 ± 0.0 (0.00 ± 0.00) | 0.2 ± 0.0 (0.00 ± 0.00) | 0.2 ± 0.1 (0.00 ± 0.00) | 0.0 ± 0.0 (0.00 ± 0.00) | 0.1 ± 0.0 (0.54 ± 0.21) |
| | EB | *n.a.* | *n.a.* | *n.a.* | *n.a.* | 2.7 ± 0.5 (23.40 ± 4.24) |
| | BA | 1.7 ± 0.2 (0.01 ± 0.00) | 1.3 ± 0.1 (0.01 ± 0.00) | 1.9 ± 0.3 (0.02 ± 0.00) | **4.2 ± 0.1 (0.04 ± 0.00)** | 0.1 ± 0.0 (0.62 ± 0.08) |
| | BB | *n.a.* | *n.a.* | *n.a.* | *n.a.* | **2.9 ± 0.4 (20.10 ± 2.42)** |
| | Ester total | 1.9 ± 0.3 (0.02 ± 0.00) | 1.5 ± 0.1 (0.01 ± 0.00) | 2.1 ± 0.4 (0.02 ± 0.00) | 4.2 ± 0.1 (0.04 ± 0.00) | 5.7 ± 0.6 (44.65 ± 4.70) |
| Ester yields, mg/g glucose (mM/M) | EA | 1.5 ± 0.3 (3.13 ± 0.66) | 8.9 ± 9.7 (18.09 ± 19.82) | 4.3 ± 1.8 (8.85 ± 3.62) | 0.9 ± 0.2 (1.79 ± 0.41) | 0.6 ± 0.3 (1.32 ± 0.70) |
| | EB | *n.a.* | *n.a.* | *n.a.* | *n.a.* | 42.5 ± 21.0 (65.97 ± 32.56) |
| | BA | 9.9 ± 2.0 (15.32 ± 3.16) | 56.9 ± 67.4 (88.27 ± 104.51) | 37.2 ± 6.4 (57.72 ± 9.96) | **86.8 ± 5.8 (134.61 ± 8.96)** | 1.2 ± 0.7 (1.81 ± 1.05) |
| | BB | *n.a.* | *n.a.* | *n.a.* | *n.a.* | **53.3 ± 30.1 (66.57 ± 37.58)** |
| | Ester total | 11.4 ± 2.4 (18.45 ± 3.82) | 65.8 ± 77.1 (106.36 ± 124.32) | 41.5 ± 8.2 (66.57 ± 13.58) | 87.7 ± 6.0 (136.40 ± 9.38) | 97.6 ± 52.1 (135.68 ± 71.90) |
| Culture time (h) | | 24 | 24 | 24 | 24 | 24 |

**Supplementary Table S16.** Summary of butyryl-CoA-derived esters production in EcJWBA14, EcJWEB7, and EcJWBB8 in anaerobic bottles with pH-adjustment. Abbreviations: *n.d.*: not detected. *n.a.*: not applicable.

| Strains | | EcJWBA14 | EcJWEB7 | EcJWBB8 |
|---|---|---|---|---|
| $O.D._{600}$ | | 3.76 ± 0.20 | 4.36 ± 0.26 | 4.22 ± 0.20 |
| Consumed glucose, g/L (mM) | | 11.85 ± 0.91<br>(65.80 ± 5.04) | 29.71 ± 0.23<br>(164.90 ± 1.27) | 41.40 ± 1.50<br>(229.78 ± 8.32) |
| Lactate, g/L (mM) | | 0.06 ± 0.00<br>(0.68 ± 0.02) | 0.17 ± 0.01<br>(1.93 ± 0.15) | 1.05 ± 0.17<br>(11.71 ± 1.85) |
| Formate, g/L (mM) | | 2.14 ± 0.06<br>(46.55 ± 1.20) | 0.18 ± 0.01<br>(3.89 ± 0.18) | 3.58 ± 0.57<br>(77.81 ± 12.33) |
| Acetate, g/L (mM) | | 1.03 ± 0.01<br>(17.17 ± 0.23) | 0.22 ± 0.00<br>(3.61 ± 0.04) | 0.40 ± 0.02<br>(6.65 ± 0.38) |
| Ethanol, g/L (mM) | | 2.05 ± 0.02<br>(44.56 ± 0.51) | 12.52 ± 0.10<br>(271.74 ± 2.25) | 9.88 ± 0.28<br>(214.41 ± 5.98) |
| Butanol, g/L (mM) | | 0.16 ± 0.01<br>(2.11 ± 0.17) | 0.03 ± 0.00<br>(0.46 ± 0.03) | 0.20 ± 0.02<br>(2.76 ± 0.33) |
| Ester titers, mg/L (mM) | EA | 39.8 ± 7.6<br>(0.45 ± 0.09) | 58.3 ± 9.3<br>(0.66 ± 0.11) | 1.2 ± 0.5<br>(0.01 ± 0.01) |
| | EB | *n.d.* | **408.9 ± 44.3**<br>**(3.52 ± 0.38)** | 381.9 ± 24.7<br>(3.29 ± 0.21) |
| | BA | **441.4 ± 40.9**<br>**(3.80 ± 0.35)** | 0.9 ± 0.1<br>(0.01 ± 0.00) | 7.4 ± 1.2<br>(0.06 ± 0.01) |
| | BB | *n.d.* | 10.1 ± 1.0<br>(0.07 ± 0.01) | **449.6 ± 43.0**<br>**(3.12 ± 0.30)** |
| | Ester total | 481.1 ± 48.4<br>(4.25 ± 0.44) | 478.2 ± 54.8<br>(4.26 ± 0.50) | 839.9 ± 68.4<br>6.48 ± 0.52) |
| Specific ester productivity, mg/gDCW/h (μM/gDCW/h) | EA | 0.4 ± 0.1<br>(0.01 ± 0.00) | 0.3 ± 0.0<br>(0.00 ± 0.00) | 0.0 ± 0.0<br>(0.00 ± 0.00) |
| | EB | *n.a.* | 2.0 ± 0.1<br>(0.02 ± 0.00) | 1.9 ± 0.2<br>(0.02 ± 0.00) |
| | BA | 4.9 ± 0.3<br>(0.04 ± 0.00) | 0.0 ± 0.0<br>(0.00 ± 0.00) | 0.0 ± 0.0<br>(0.00 ± 0.00) |
| | BB | *n.a.* | 0.1 ± 0.0<br>(0.00 ± 0.00) | 2.3 ± 0.3<br>(0.02 ± 0.00) |
| | Ester total | 5.4 ± 0.4<br>(0.05 ± 0.00) | 2.4 ± 0.1<br>(0.02 ± 0.00) | 4.2 ± 0.6<br>(0.03 ± 0.00) |
| Ester yields, mg/g glucose (mM/M) | EA | 3.34 ± 0.38<br>(0.01 ± 0.00) | 2.0 ± 0.3<br>(0.00 ± 0.00) | 0.0 ± 0.0<br>(0.00 ± 0.00) |
| | EB | *n.a.* | 13.7 ± 1.4<br>(0.02 ± 0.00) | 9.4 ± 0.6<br>(0.01 ± 0.00) |
| | BA | 37.2 ± 0.6<br>(0.06 ± 0.00) | 0.0 ± 0.0<br>(0.00 ± 0.00) | 0.2 ± 0.0<br>(0.00 ± 0.00) |
| | BB | *n.a.* | 0.3 ± 0.0<br>(0.00 ± 0.00) | 11.1 ± 1.1<br>(0.01 ± 0.00) |
| | Ester total | 40.6 ± 1.0<br>(0.06 ± 0.00) | 16.0 ± 1.7<br>(0.03 ± 0.00) | 20.7 ± 1.7<br>(0.03 ± 0.00) |
| Culture time (h) | | 48 | 96 | 96 |

**Supplementary Table S17.** Summary of BA production by EcJWBA15, and EcJWBA16. Abbreviations: *n.d.*: not detected. *n.a.*: not applicable.

| Target product | | BA | |
|---|---|---|---|
| Strains | | EcJWBA15 | |
| Culture type | | Tubes | Anaerobic bottles with pH-adjustment |
| $O.D._{600}$ | | 3.80 ± 0.38 | 3.37 ± 0.15 |
| Consumed glucose, g/L (mM) | | 3.26 ± 0.27 (18.11 ± 1.51) | 6.45 ± 0.27 (35.81 ± 1.48) |
| Lactate, g/L (mM) | | 0.13 ± 0.06 (1.49 ± 0.64) | 0.10 ± 0.02 (1.10 ± 0.22) |
| Formate, g/L (mM) | | 1.47 ± 0.29 (31.91 ± 6.25) | 2.37 ± 0.06 (51.52 ± 1.36) |
| Acetate, g/L (mM) | | 0.86 ± 0.05 (14.26 ± 0.83) | 1.14 ± 0.01 (18.91 ± 0.22) |
| Ethanol, g/L (mM) | | 1.42 ± 0.17 (30.93 ± 3.67) | 1.41 ± 0.05 (30.63 ± 1.18) |
| Butanol, g/L (mM) | | 0.18 ± 0.03 (2.39 ± 0.44) | 0.07 ± 0.01 (0.88 ± 0.09) |
| Ester titers, mg/L (mM) | EA | 14.4 ± 0.8 (0.16 ± 0.01) | 28.7 ± 2.6 (0.3 ± 0.0) |
| | EB | *n.d.* | *n.d.* |
| | BA | 259.5 ± 11.6 (2.23 ± 0.10) | 636.3 ± 43.8 (5.5 ± 0.4) |
| | BB | *n.d.* | *n.d.* |
| | Ester total | 273.8 ± 12.4 (2.40 ± 0.11) | 665.0 ± 44.3 (5.8 ± 0.4) |
| Specific ester productivity, mg/gDCW/h (μM/gDCW/h) | EA | 0.3 ± 0.0 (0.00 ± 0.00) | 0.2 ± 0.0 (0.00 ± 0.00) |
| | EB | *n.a.* | *n.a.* |
| | BA | 6.1 ± 0.5 (0.05 ± 0.00) | 4.1 ± 0.2 (0.03 ± 0.00) |
| | BB | *n.a.* | *n.a.* |
| | Ester total | 6.4 ± 0.6 (0.06 ± 0.00) | 4.2 ± 0.2 (0.04 ± 0.00) |
| Ester yields, mg/g glucose (mM/M) | EA | 4.2 ± 0.2 (0.01 ± 0.00) | 4.5 ± 0.3 (0.01 ± 0.00) |
| | EB | *n.a.* | *n.a.* |
| | BA | 75.9 ± 2.2 (0.12 ± 0.00) | 98.9 ± 10.4 (0.15 ± 0.02) |
| | BB | *n.a.* | *n.a.* |
| | Ester total | 80.1 ± 2.4 (0.13 ± 0.00) | 103.3 ± 10.5 (0.16 ± 0.02) |
| Culture time (h) | | 24 | 96 |

**Supplementary Table S18.** Summary of direct fermentative production of butyryl-CoA-derived esters from sugar.

| Products | Strain ID | Host strain | AATs | Engineering Strategies | Titers (mg/L) | Volumetric productivity (mg/L/h) | Yields (mg ester/ g glucose) | Selectivity (%) | *Maximum Theoretical yield (%) | Ref. |
|---|---|---|---|---|---|---|---|---|---|---|
| BA | *E. coli* | *E. coli* | $SAAT_{Fa}$ | - | 0.5 | 0.005 | 0.01 | 0.3 | 0.002 | [5] |
| | CaSAAT | *C. acetobutylicum* | $SAAT_{Fa}^{opt}$ | - | 8.4 | 0.12 | 0.2 | 14.2 | 0.07 | [8] |
| | EcJWBA2 | *E. coli* | $ATF1_{Sc}$ | - | 34.2 | 1.43 | 8.2 | 74.3 | 1.9 | This study |
| | EcJWBA2 | *E. coli* | $ATF1_{Sc}$ | Optimizing induction conditions | 48.0 | 2.00 | 10.9 | 83.1 | 2.5 | This study |
| | EcJWBA10 | *E. coli* | $TrxA\_ATF1_{Sc}^{opt}$ | +Improving soluble expression of $ATF1_{Sc}$ | 89.5 | 3.73 | 16.7 | 84.3 | 3.9 | This study |
| | EcJWBA14 | *E. coli* | $TrxA\_ATF1_{Sc}^{opt}$ | +Improving soluble expression of $AdhE2_{Ca}$ | 203.0 | 8.46 | 86.8 | 99.0 | 20.2 | This study |
| | EcJWBA15 | *E. coli* | $TrxA\_ATF1_{Sc}^{opt}$ | +Using an *adhE* deletion strain | 259.5 | 10.81 | 75.9 | 94.8 | 17.7 | This study |
| | EcJWBA15 | *E. coli* | $TrxA\_ATF1_{Sc}^{opt}$ | +Culturing the strain under anaerobic conditions with pH-adjustment | 636.3 | 6.63 | 98.9 | 95.7 | 23.0 | This study |
| EB | EcDL204 | *E. coli* | $SAAT_{Fa}$ | - | 134.0 | 1.40 | 3.3 | 78.2 | 0.8 | [5] |
| | CaSAAT | *C. acetobutylicum* | $SAAT_{Fa}^{opt}$ | - | 0.6 | 0.008 | 0.02 | 1.0 | 0.005 | [8] |
| | EcJWEB2 | *E. coli* | $SAAT_{Fa}$ | - | 71.0 | 2.96 | 9.8 | 92.0 | 2.3 | This study |
| | EcJWEB2 | *E. coli* | $SAAT_{Fa}$ | Optimizing induction conditions | 200.4 | 8.35 | 20.6 | 89.6 | 4.8 | This study |
| | EcJWEB7 | *E. coli* | $SAAT_{Fa}$ + GroES/EL | +Improving soluble expression of $SAAT_{Fa}$ | 365.7 | 15.23 | 38.5 | 92.9 | 8.9 | This study |
| | EcJWEB7 | *E. coli* | $SAAT_{Fa}$ + GroES/EL | +Culturing the strain under anaerobic conditions with pH-adjustment | 408.9 | 4.26 | 36.5 | 85.5 | 8.5 | This study |
| BB | EcDL204 | *E. coli* | $SAAT_{Fa}$ | - | 36.8 | 0.38 | 0.9 | 21.5 | 0.2 | [5] |
| | CaSAAT | *C. acetobutylicum* | $SAAT_{Fa}^{opt}$ | - | 50.1 | 0.70 | 1.7 | 84.8 | 0.4 | [8] |
| | EcJWBB2 | *E. coli* | $SAAT_{Fa}$ | - | 33.5 | 1.40 | 8.7 | 20.8 | 2.2 | This study |
| | EcJWBB2 | *E. coli* | $SAAT_{Fa}$ | Optimizing induction conditions | 127.4 | 5.31 | 26.3 | 34.0 | 6.6 | This study |
| | EcJWBB7 | *E. coli* | $SAAT_{Fa}$ + GroES/EL | +Improving soluble expression of $SAAT_{Fa}$ | 167.3 | 7.00 | 53.3 | 50.6 | 13.3 | This study |
| | EcJWBB8 | *E. coli* | $SAAT_{Fa}$ + $TrxA\_adhE2^{opt}$ | +Improving soluble expression of $AdhE2_{Ca}$ +Culturing the strain under anaerobic conditions with pH-adjustment | 449.6 | 4.68 | 40.1 | 53.5 | 10.0 | This study |

*Calculated based on consumed glucose.

**Supplementary Figure S1.** SDS-PAGE analysis results of (**a**) EcJWBA1-6. (**b**) EcJWEB1-6. (**c**) EcJWBB1-6. Red arrows indicate the expected size of overexpressed enzymes. Abbreviations: L, protein ladder; T, total fractions; S, Soluble fractions. (Induction conditions: 37℃, 0.5 mM IPTG)

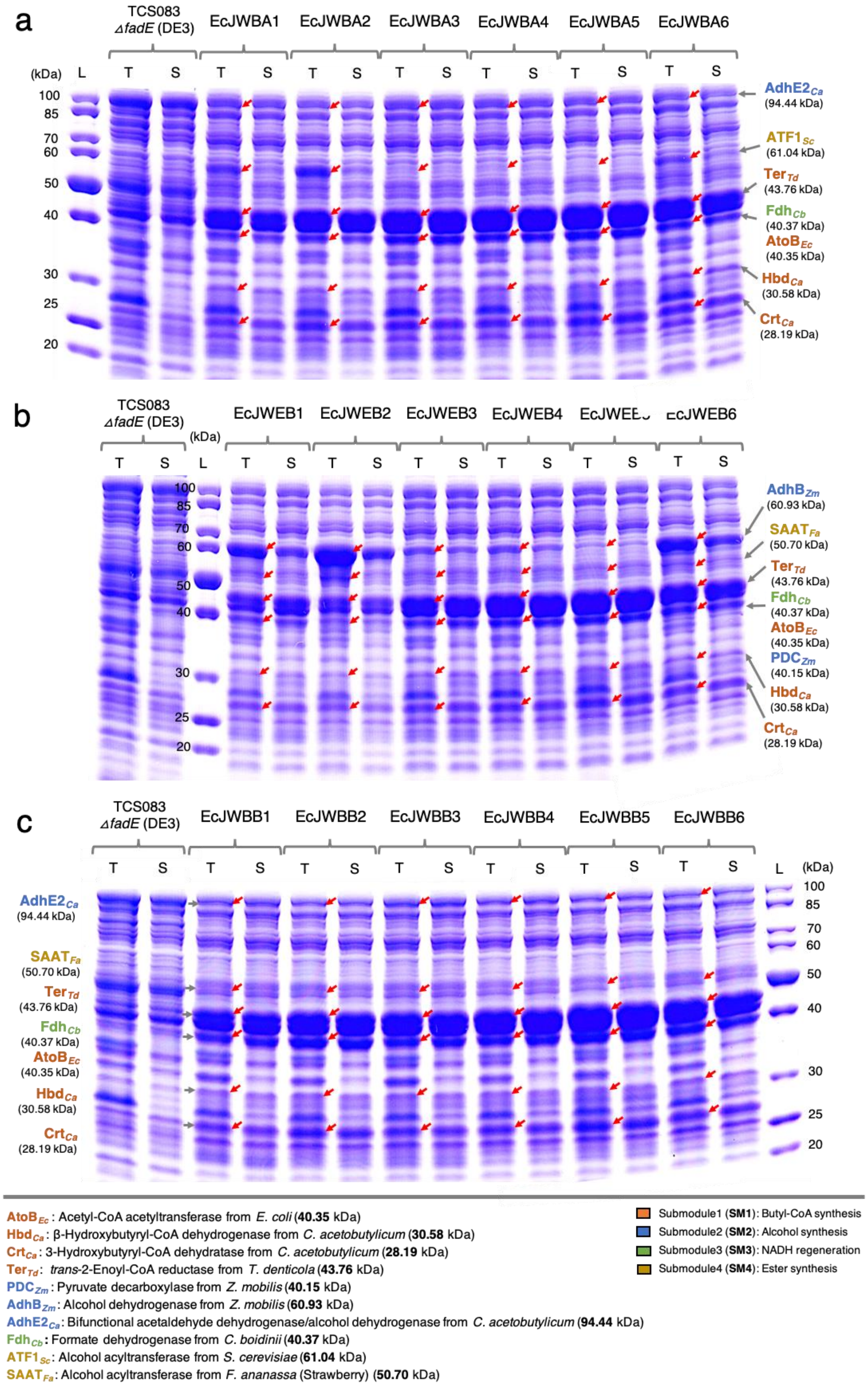

**Supplementary Figure S2.** Results of induction conditions optimization. (**a**) BA production in EcJWBA2. (**b**) EB production in EcJWEB2. (**c**) BB production in EcJWBB2. Abbreviations: EA, ethyl acetate; EB, ethyl butyrate; BA, butyl acetate; and BB, butyl butyrate.

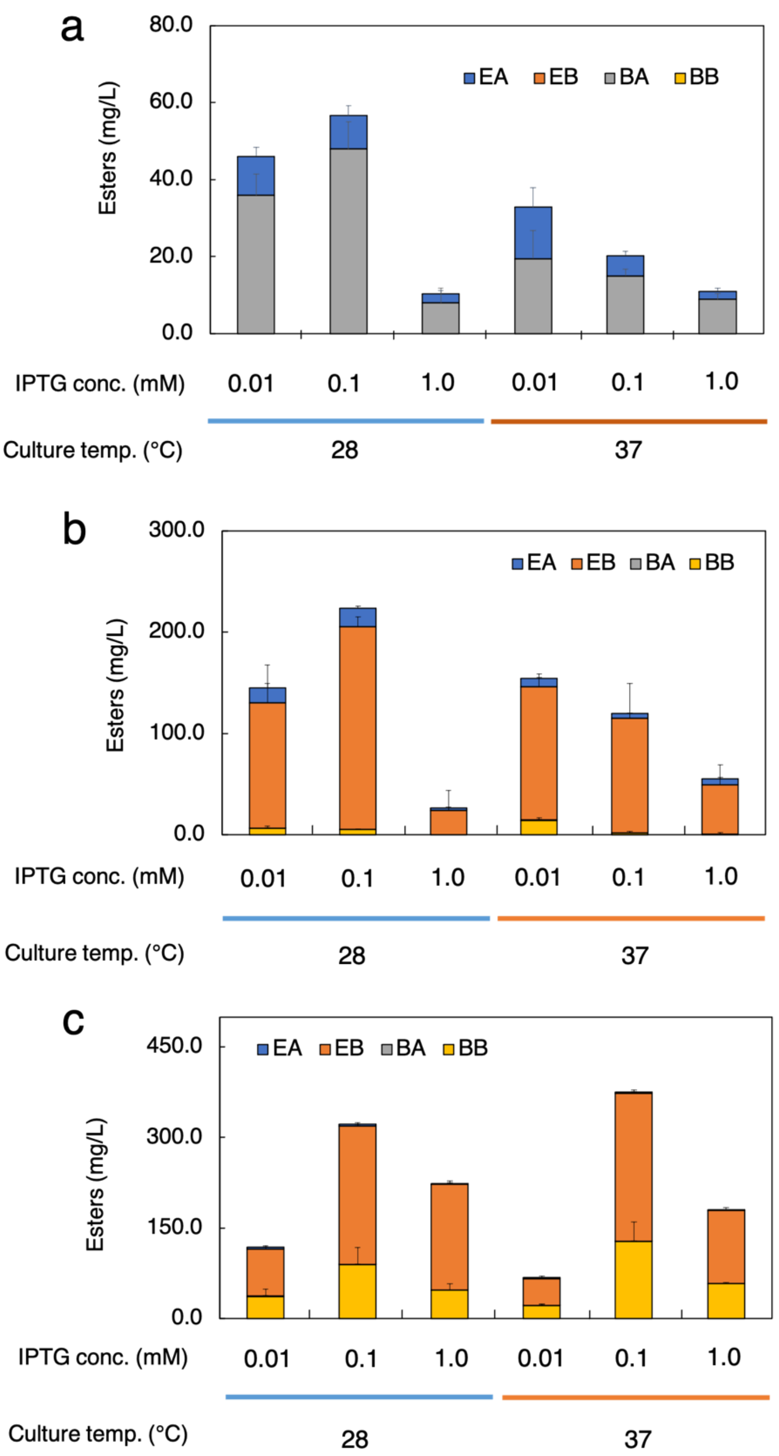

**Supplementary Figure S3.** SDS-PAGE analysis results of EcJWBA2, EcJWEB2, and EcJWBB2 with various induction conditions. (**a**) EcJWBA2. (**b**) EcJWEB2. (**c**) EcJWBB2. Red arrows indicate the expected size of overexpressed enzymes. Yellow stars indicate the conditions result in the highest esters production among the evaluated conditions. Abbreviations: L, protein ladder; T, total fractions; S, Soluble fractions. (Induction conditions: 28/37°C, 0.01/0.1/1.0 mM IPTG)

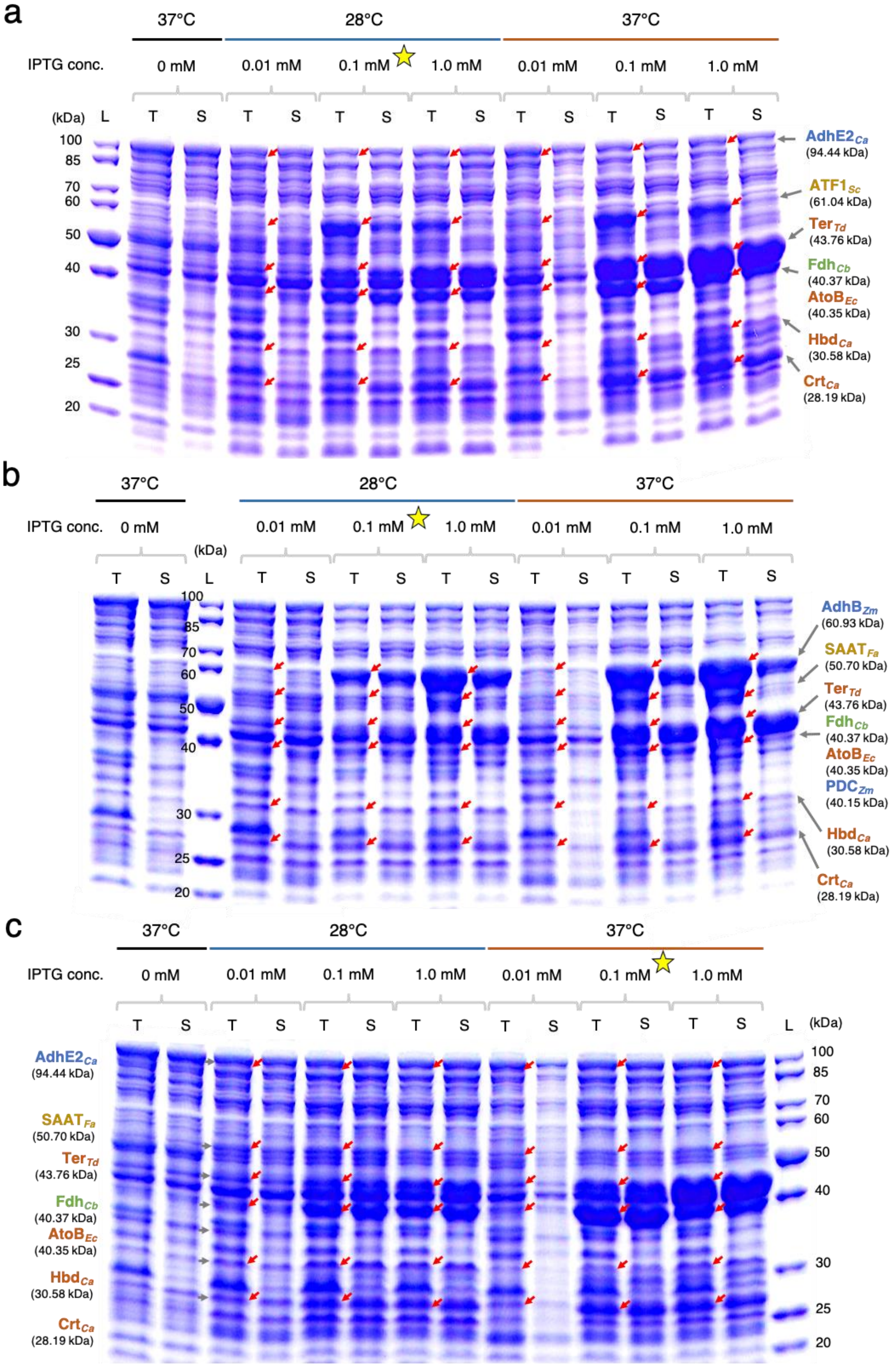

**Supplementary Figure S4.** SDS-PAGE analysis results of butyryl-CoA-derived esters producing strains with various induction conditions. (**a**) EcJWBA2, and EcJWBA7-10. (**b**) EcJWEB7. (**c**) EcJWBB7 Red arrows indicate the expected size of overexpressed enzymes. Abbreviations: L, protein ladder; T, total fractions; S, Soluble fractions. (Induction conditions: EcJWBA7-10 and EcJWEB7: 28℃, EcJWBB7: 37℃, 0.1 mM IPTG).

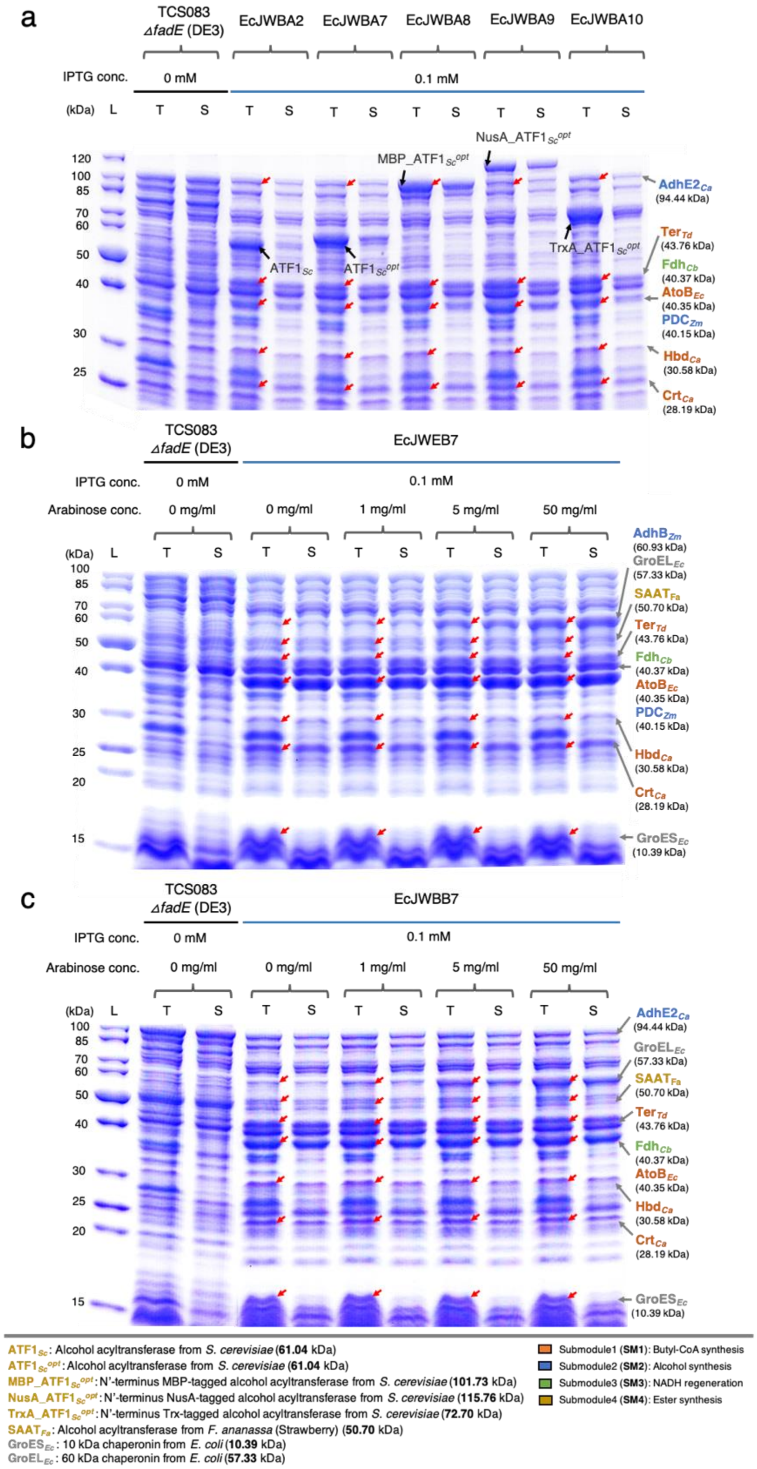

**Supplementary Figure S5.** SDS-PAGE analysis results of EcJWBA10-14. Red arrows indicate the expected size of overexpressed enzymes. Abbreviations: L, protein ladder; T, total fractions; S, Soluble fractions. (Induction conditions: 28℃, 0.1 mM IPTG).

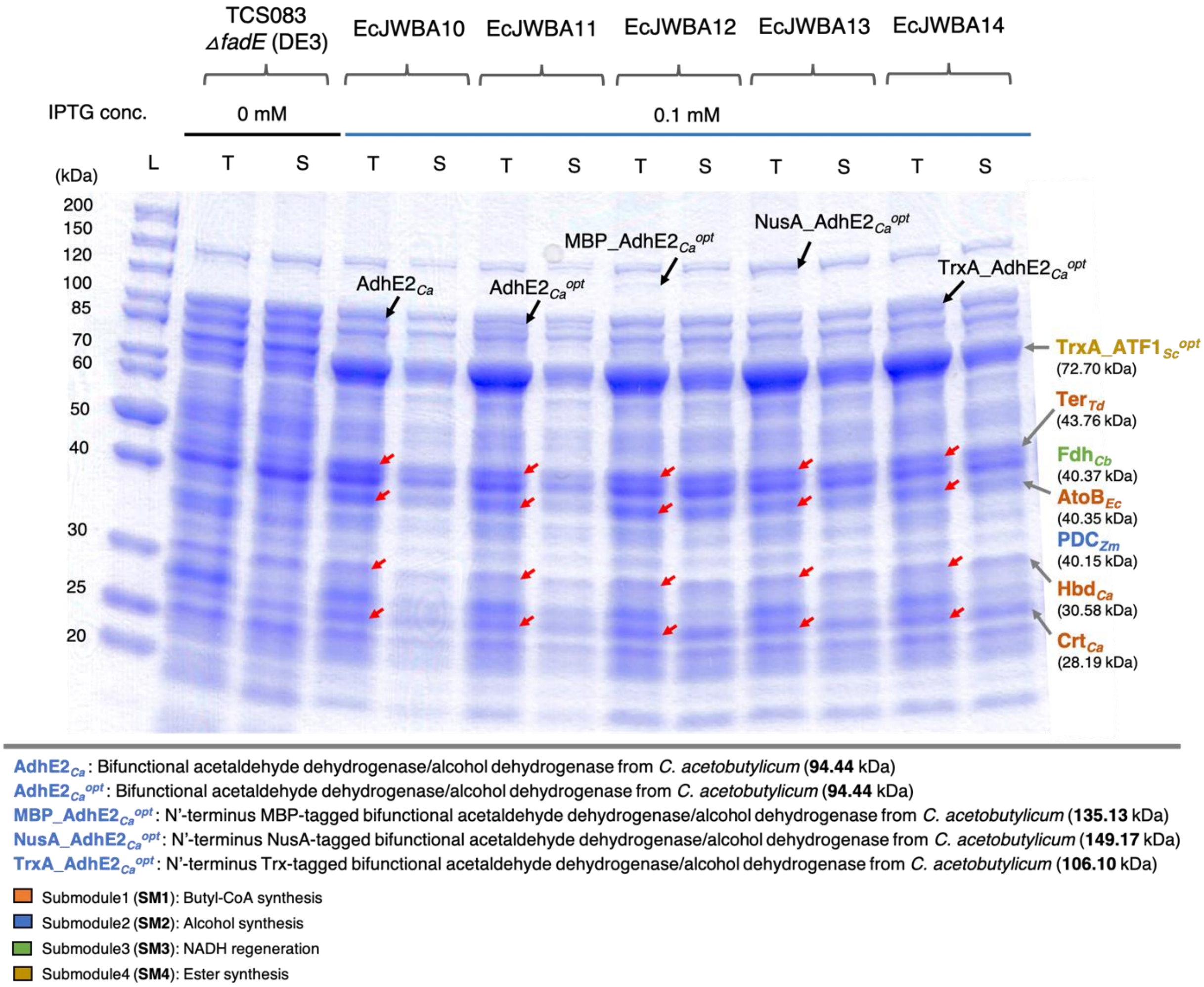

**Supplementary Figure S6.** Predicted solubility of enzymes used in this study. The solubility of enzymes were predicted with Protein-Sol[9] (https://protein-sol.manchester.ac.uk) from the amino acid sequences. The AVER value (0.45) indicates the population average for the experimental dataset[10] (in grey). Thus, while the solubility values greater than 0.45 is predicted to have a higher solubility than the average soluble *E. coli* protein (in orange), the values lower than 0.45 is predicted to be less soluble (in blue).

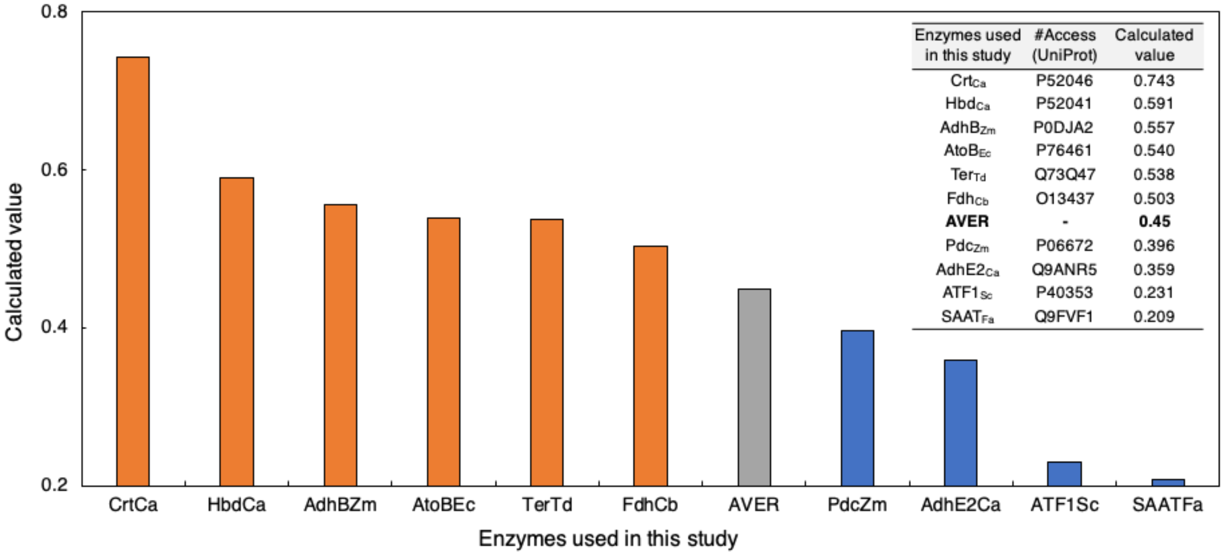

**Supplementary Figure S7.** Stoichiometric balance for synthesis of butyryl-CoA-derived esters. (**a**) Reactions in modular biosynthetic pathways of butyryl-CoA-derived esters from glucose. (**b**) Net reactions for biosynthesis of butyl acetate (BA), ethyl butyrate (EB), or butyl butyrate (BB) from glucose. *Ester yield on glucose.

a

| | Pathways | Reactions | #Modules |
|---|---|---|---|
| A | Glycolysis | Glucose → 2 Pyruvate + 2 NADH | Endogenous |
| B | Pfl reaction | Pyruvate → Acetyl-CoA + Formate | Endogenous |
| C | Fdh reaction | Formate → $CO_2$ + NADH | 3 |
| D | Butyl-CoA synthesis | 2 Acetyl-CoA + 2 NADH → Butyl-CoA | 1 |
| E | AdhE2 reaction | Butyl-CoA + 2 NADH → Butanol | 2 |
| F | PDC/adhE reaction | Pyruvate + NADH → Ethanol | 2 |
| G | BA synthesis | Butanol + Acetyl-CoA → Butyl acetate | 4 |
| H | EB synthesis | Ethanol + Butyl-CoA → Ethyl butyrate | 4 |
| I | BB synthesis | Butanol + Butyl-CoA → Butyl butyrate | 4 |

b

| Products | Stoichiometric equations | Net Reactions | Maximum theoretical yield* | |
|---|---|---|---|---|
| | | | Cmol/Cmol | g/g |
| BA | 1.5A + 3B + 3C + D + E + G | 1.5 Glucose + 2 $NAD^+$ → Butyl acetate + 3 $CO_2$ + 2 NADH | 0.67 | 0.43 |
| EB | 1.5A + 2B + 2C + D + F + H | 1.5 Glucose + 2 $NAD^+$ → Ethyl butyrate + 3 $CO_2$ + 2 NADH | 0.67 | 0.43 |
| BB | 2A + 4B + 4C + 2D + E + I | 2 Glucose + 2 $NAD^+$ → Butyl butyrate + 4 $CO_2$ + 2 NADH | 0.50 | 0.40 |
